# Tracing genetic connections of ancient Hungarians to the 6-14^th^ century populations of the Volga-Ural region

**DOI:** 10.1101/2022.02.04.478947

**Authors:** Bea Szeifert, Dániel Gerber, Veronika Csáky, Péter Langó, Dmitrii A. Stashenkov, Aleksandr A. Khokhlov, Ayrat G. Sitdikov, Ilgizar R. Gazimzyanov, Elizaveta V. Volkova, Natalia P. Matveeva, Alexander S. Zelenkov, Olga E. Poshekhonova, Anastasiia V. Sleptsova, Konstantin G. Karacharov, Viktoria V. Ilyushina, Boris A. Konikov, Flarit A. Sungatov, Alexander G. Kolonskikh, Sergei G. Botalov, Ivan V. Grudochko, Oleksii Komar, Balázs Egyed, Balázs G. Mende, Attila Türk, Anna Szécsényi-Nagy

## Abstract

Most of the early Hungarian tribes originated from the Volga-Kama and South-Ural regions, where they were composed of a mixed population based on historical, philological, and archaeological data. We present here the uniparental genetic makeup of the medieval era of these regions that served as a melting pot for ethnic groups with different linguistic and historical backgrounds. Representing diverse cultural contexts, the new genetic data originates from ancient proto-Ob-Ugric people from Western Siberia (6^th^-13^th^ century), the pre-Conquest period, and subsisting Hungarians from the Volga-Ural region (6^th^-14^th^ century) and their neighbours. By examining the eastern archaeology traits of Hungarian prehistory, we also study their genetic composition and origin in an interdisciplinary framework.

We analysed 110 deep-sequenced mitogenomes and 42 Y-chromosome haplotypes from 18 archaeological sites in Russia. The results support the studied groups’ genetic relationships regardless of geographical distances, suggesting large-scale mobility. We detected long-lasting genetic connections between the sites representing the Kushnarenkovo and Chiyalik cultures and the Carpathian Basin Hungarians and confirmed the Uralic transmission of several East-Eurasian uniparental lineages in their genepool. Based on phylogenetics, we demonstrate and model the connections and splits of the studied Volga-Ural and conqueror groups.

Early Hungarians and their alliances conquered the Carpathian Basin around 890 AD. Re-analysis of the Hungarian conquerors’ maternal genepool reveals numerous surviving maternal relationships in both sexes; therefore, we conclude that men and women came to the Carpathian Basin together, and although they were subsequently genetically fused into the local population, certain eastern lineages survived for centuries.

## Introduction

The Hungarians are the sole Uralic-speaking people in Central Europe today. The earliest known settlement area that can be associated with their ancestors is the territory bordered by the Rivers Tobol, Irtysh, and Ishim in the Trans-Urals and the western zone of south-western Siberia. Their artefacts appear among the finds left by the descendants of the Iron Age Sargat cultures, on both sides of the Urals, mainly in the distribution areas of the early medieval Bakal and Potchevash cultures (1) (2) (3) (4). In the 6^th^-8^th^ centuries the early Hungarians (together with other groups) lived most probably in the southern Ural region, and their archaeological remains were a substantial part of the Kushnarenkovo-Karayakupovo culture (2) (5) (6). At the beginning of the 9^th^ century AD, the ancestors of the Hungarians crossed the River Volga and moved to the territory lying to the north of the Black Sea (Subbotsi-type sites), where they became the neighbours of the Khazars and Slavic-speaking peoples

(2) (7) (8) (9). Later, leaving the Khazar Khaganate along with Kabars, they settled in the Carpathian Basin around 890 AD (10) (6) (2) (9). Meanwhile, Hungarians who remained in the Volga–Ural region were reported in the middle of the 13^th^ century (11), whose tangible heritage was associated with the Chiyalik culture in archaeological research, belonging to the area known as *Magna Hungaria* (5) (12) (13) (14) (15). Historical and linguistic data suggest that a part of the Hungarians conquering the Carpathian Basin came from the Southern Urals and Trans-Urals, which is also supported by the findings of archaeological research (2) (6) (8) (9) (16).

The Volga–Ural region witnessed several waves of migration, and its population was extremely complex in both historical and genetic terms. There is little genetic information about the medieval period of Central Eurasia but studies on its Iron and Bronze Age populations (17) (18) (19) serve as important reference points. The current residents of the area have been mainly studied genetically in the context of the Uralic language family. In most cases, similarities could be pointed out among the groups belonging to the Uralic language family living in the Volga-Ural region. Further connections could also be detected with their geographical neighbours, the Bashkir, Chuvash and Tatar groups (20) (21). The Kushnarenkovo culture (from the Trans-Uralic Uyelgi site) and those representing the Lomovatovo and Nevolino cultures associated populations (Cis-Ural is the western foreland of the Urals) show extensive genetic connections to the conquering Hungarians (22).

The composition of the uniparental genetic lines (23) (24) (25) (26) of the Hungarians living today in the Carpathian Basin is similar to that of other European peoples (20), but the maternal (22) (27) (28) (29) (30) (31) (32) (33) and paternal (32) (34) (35) lineages of the population of the archaeological Conquest period in Carpathian Basin in the late 9^th^-10^th^ centuries (henceforth: conqueror Hungarians, in short conquerors) show a different picture. Their maternal lineages are similar to those of modern Tatars, with a significant Eastern Eurasian component. On this basis, relationships with the Potapovka, Poltavka, and Srubnaya cultures’ associated populations were suggested, and connections with the Scythians and Huns were raised (28). Due to the substantial time gap, however, the connections with these groups are rather indirect. Conqueror paternal lineage composition is most similar to today’s Bashkirs (34)(35). Fóthi et al. traced the origins of these paternal lineages to three areas between the Lake Baikal and the Altai Mountains, between Western Siberia and the Southern Urals, and to the regions of the Black Sea and the North Caucasus (34), but due to deep genetic divergence dates, firm conclusions on tribal origins cannot be drawn from this observation. The remains of King Bela III (ruled between 1172–1196 AD) have been genetically analysed recently. He is the sole member of the first Hungarian dynasty whose grave was found *in-situ* in Székesfehérvár, Hungary. Based on previous autosomal analysis he shares most of his genetic ancestry with the representatives of ancient and modern-day Europeans (36). His Y-chromosome belongs to the R1a-Z2123 haplogroup, which has the highest frequency in Central Asia today but is also found in the Volga-Ural region, which is in line with earlier observations and the significance of the Volga-Ural region in the formation of the early Hungarian elite (37) (38).

Previous genetic studies on the 10^th^ to 12^th^ century Carpathian Basin population divided individuals into two groups of “elite” and “common” people, based on the funerary furnishing and burial customs (27) (28) (29) (30) (31) (32) (33) (34) (35). As these concepts have become archaeologically incomprehensible and vague today, a division with a well-defined description focusing on the number of graves discovered in the cemeteries and the chronology of their usage was introduced (39) (40) (for a summary of this classification, see Supplementary Material, Chapter A, 8.). Our analyses adopted this approach by dividing the conqueror dataset into three main groups based on Kovács (39): KL-IV (largely corresponding to the former “elite” group), which is characterised by small 10^th^ century cemeteries of the camps that were the primary settlements of the nomadic Hungarians; KL-V and KL-VI (largely corresponding to former “commoners” group). KL-V represents 10^th-^century cemeteries of villages with a large number of burials and KL-VI contains large village cemeteries opened in the 10^th^ century and used until the 11^th^ and 12^th^ centuries.

Our study focuses on regions in present-day Russia that were important as the genesis of several Turkic (e.g., Bashkirs, Tatars) and Uralic speaking groups, such as the Maris, Khantys and the Hungarians as well. We analyse the uniparental markers of 112 individuals (6^th^–14^th^ centuries) representing 13 sites in the region of the Volga and Southern Urals (collectively the Volga-Ural region), as well as five sites associated with early Ob-Ugric people (6^th^-13^th^ century) living in Western Siberia (for more information about the studied sites, see Fig. 1, Methods and Supplementary Material, Chapter A and Table S1). Our objective has been to characterise genetically the cultural and ethnic hub of the Volga-Ural region, focusing on sites that can be linked to the early Hungarians archaeologically or their neighbours in a geographical sense, and to determine whether the possible ancestors of the Hungarians show biological link to their neighbours and the conquerors of the Carpathian Basin. We also examined how much of the genetic makeup of the Volga-Ural region was preserved by the population of the Carpathian Basin between the 10^th^ and 12^th^ centuries.

**Figure 1:**
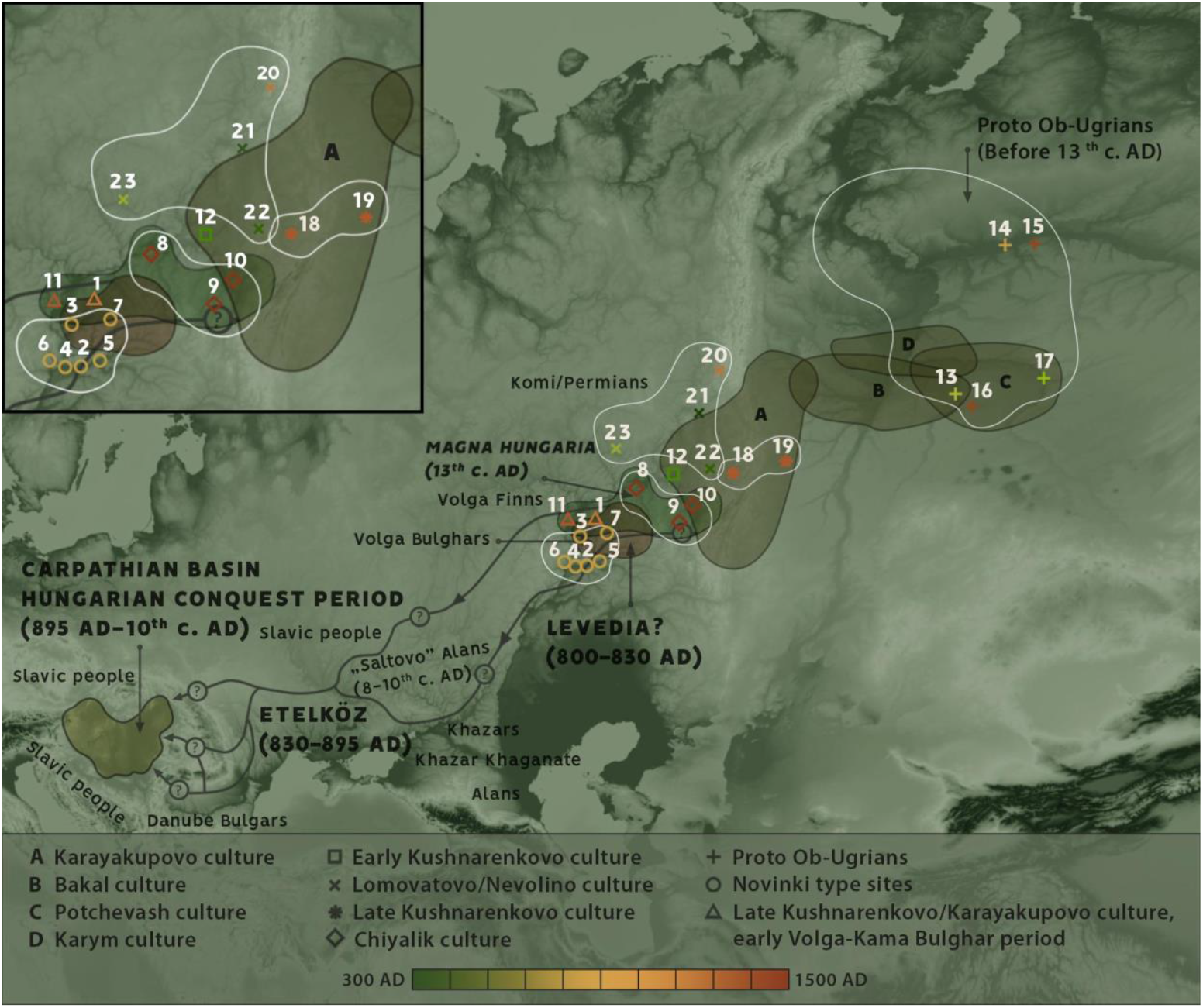
The supposed migration route of the early Hungarians (arrows) and the regions which could be linked to them (uppercase letters). The sites were grouped according to archaeological and chronological aspects; the formed groups are marked with white outlines. The investigated sites and the groups formed from them: <u>Bolshie Tigani</u> [1]; <u>Novinki group</u>: Novinki [2], Mulovka [3], Brusyany [4], Lebyazhinka [5], Malaya Ryazan [6], Shilovka [7]; <u>Chiyalik group</u>: Gulyukovo [8], Novo Hozyatovo [9], Gornovo [10]; <u>Tankeevka</u> [11]; <u>Bustanaevo</u> [12]; <u>Proto-Ob-Ugric group</u>: Vikulovo [13], Barshov Gorodok [14], Ivanov Mis [15], Panovo [16], Ust-Tara [17]; <u>Uyelgi+Karanayevo group</u>: Karanayevo [18], Uyelgi [19]; <u>Cis-Ural group</u>: Bayanovo [20], Brody [21], Bartim [22], Sukhoy Log [23] (source of 19-23.: Csáky et al 2020 (22)). Source of the map: Qgis v3.16.0 Topographic WMS-by terrestris (https://ows.terrestris.de/osm/service?). Modifications were made in Adobe Photoshop CS6 and Adobe Photoshop 2020.

In our study, we analyse 10 groups (Table 1) formed according to archaeological, chronological and geographic aspects. In addition to the investigated sites, we used the database data of the conquerors from the Carpathian Basin (22) (27) (28) (29) (KL-IV–VI), the Cis-Ural group (Bayanovo, Brody, Bartim, Sukhoy Log) and the Uyelgi site (22).

**Table 1.**
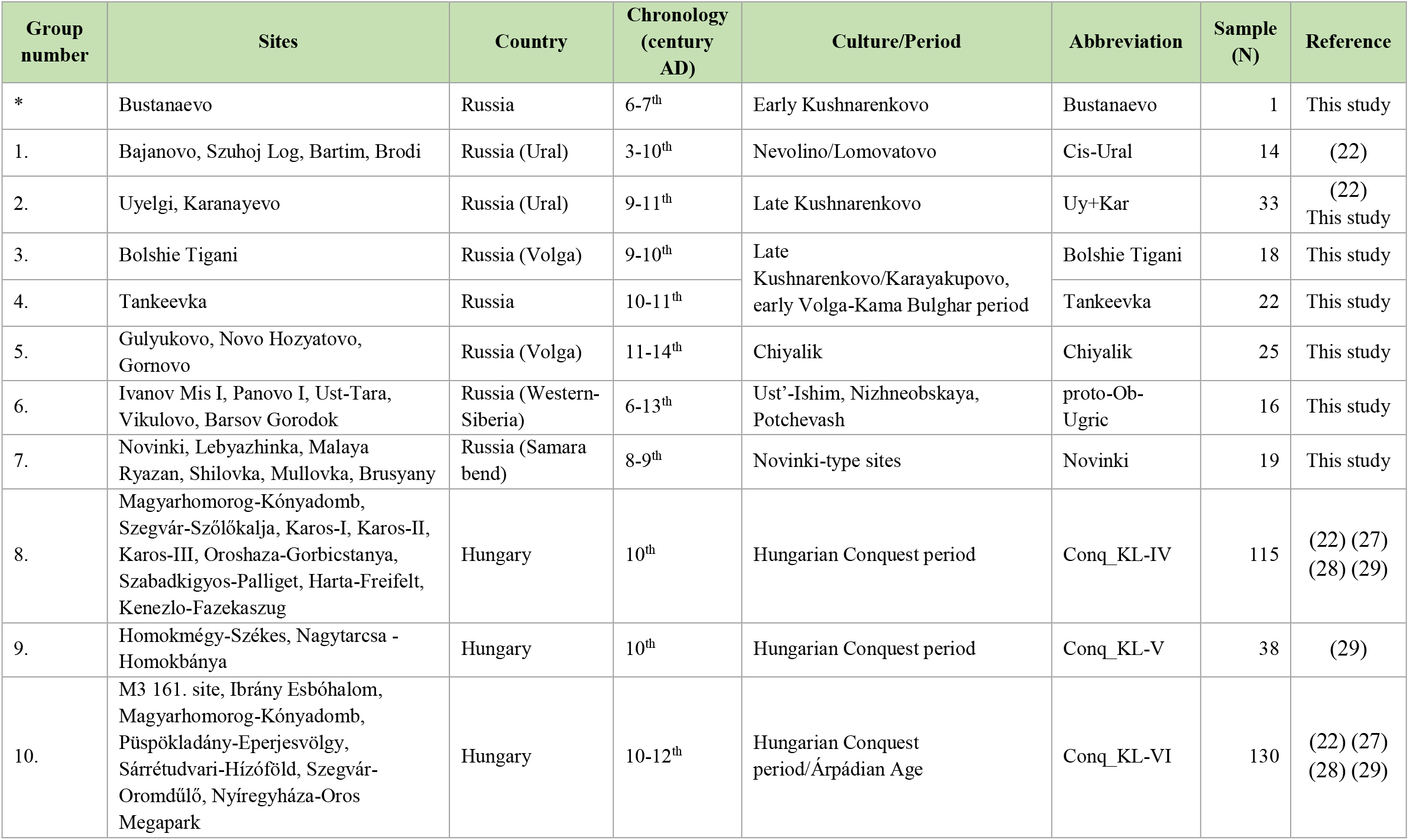
Information about groups formed in this study. The sites were grouped according to archaeological and chronological aspects. We investigated only one individual from the Bustanaevo site that was not used in population genetic analysis due to its outlying chronology. For other information see Supplementary Material, Chapter A, Table S1-S2 and Table S7.

## Results

### Primary observations and paternal lineages

Seventy-three different haplotypes could be detected based on the complete mitochondrial genome sequences obtained from 112 newly examined individuals (Supplementary Material, Table S2). These belong to 15 macro-haplogroups (A, B, C, D, F, G, H, J, K, M, N, T, U, V, Z), which – except Z – were also described in the conquerors (Supplementary Material, Table S7). Other macrohaplogroups (I, W, X, Y) are also present in conqueror groups, which may have reached them from other sources or could not be detected in the studied eastern groups due to their small sample sizes. Eastern-Western geographical distribution of mitochondrial macrohaplogroups’ frequency observable today is also reflected in the study groups, as e.g. proto-Ob-Ugric associated individuals mostly possess eastern type (A, B, C, D, G, M), and Tankeevka mostly western type (H, T, U) lineages (Fig. 2, Supplementary Material, Table S2 and Table S7).

**Figure 2:**
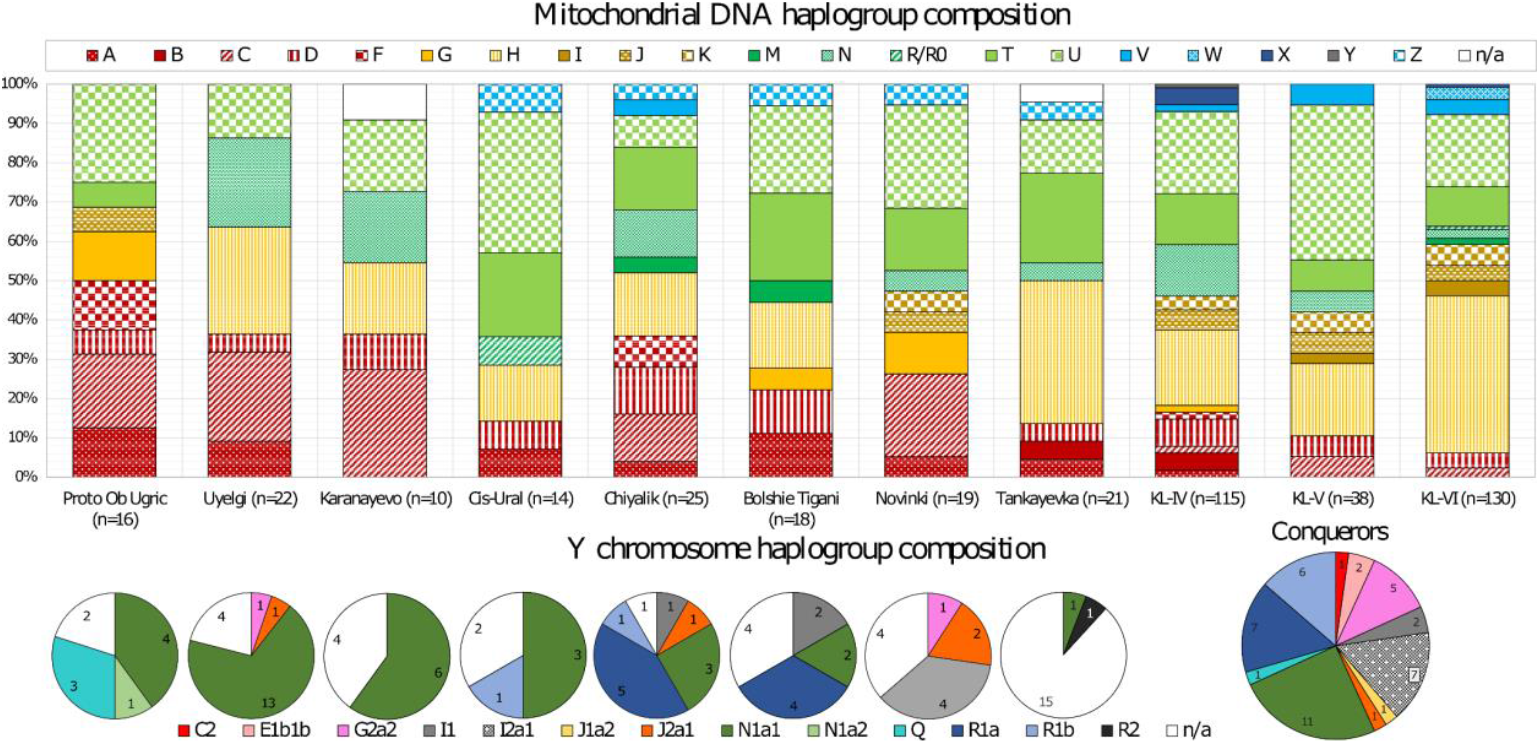
The mitochondrial (bar graph) and Y-chromosome (pie charts) haplogroup compositions of the analysed groups. In the case of the Hungarian conquerors, most of the Y-chromosomal data are known from the KL-IV group (34) (35). Due to the underrepresentation of KL-V and KL-VI groups in the Y-chromosomal dataset, the conquerors were merged into one group in this figure.

Forty-two males out of 72 yielded identifiable Y-chromosome haplotypes (see Methods and Supplementary Material, Table S2 and Table S10). The paternal lineages of the examined individuals can be classified into eight major haplogroups (G2a2, I1, J2a1, N1a1, N1a2, Q, R1a, R1b). Most of them also appear at several sites associated with the Hungarians, as well as among the Hungarian conquerors (Fig. 2). The Q-L330 subgroup (ISOGG 15.73 Q1b1a3) was detected only in the proto-Ob-Ugric group, which corresponds to its Altai or Siberian origin (41). Haplogroup N1a1-M46 is present at all sites associated with the early Hungarians and among the conquerors but could not be detected in the Novinki group. 36.8% of the known paternal lineages of the Hungarian conquerors belong to the haplogroup N, although on average it is rare (1–2%) among today’s Hungarians, compared to other members of the Uralic language family (20) (42) (43). Interestingly, this proportion is somewhat higher (6.1%) in geographically isolated regions in Hungary (44). Within N1a1-M46, the N1a-Z1936 (ISOGG v15.73 N1a1a1a1a2) subclade potentially represents the relationship between the representatives of the Uralic language family, which subgroup has also been described in the conquerors (34) (35).

Based on the STR network analysis (Fig. 3), individuals from Uyelgi+Karanayevo (Karanayevo) and Chiyalik groups (Gornovo) belong to one of the subgroups of N1a1a1a1a2a1c (ISOGG v15.73; SNP: B539/PH3340) fitting in the genetic composition of the Volga–Ural region, and they are clustered together with the samples from Uyelgi, and present-day Khantys, Mansis and Hungarians (all belonging to the Ob-Ugric branch of Uralic languages), as well as with the Bashkirs and Tatars of the Volga–Ural region (22) (26). Because the Y-SNP results from literature and the Y-STR networks are not always compatible, we cannot define more downstream SNPs and subgroups, than N1a-B539/PH3340.

**Figure 3:**
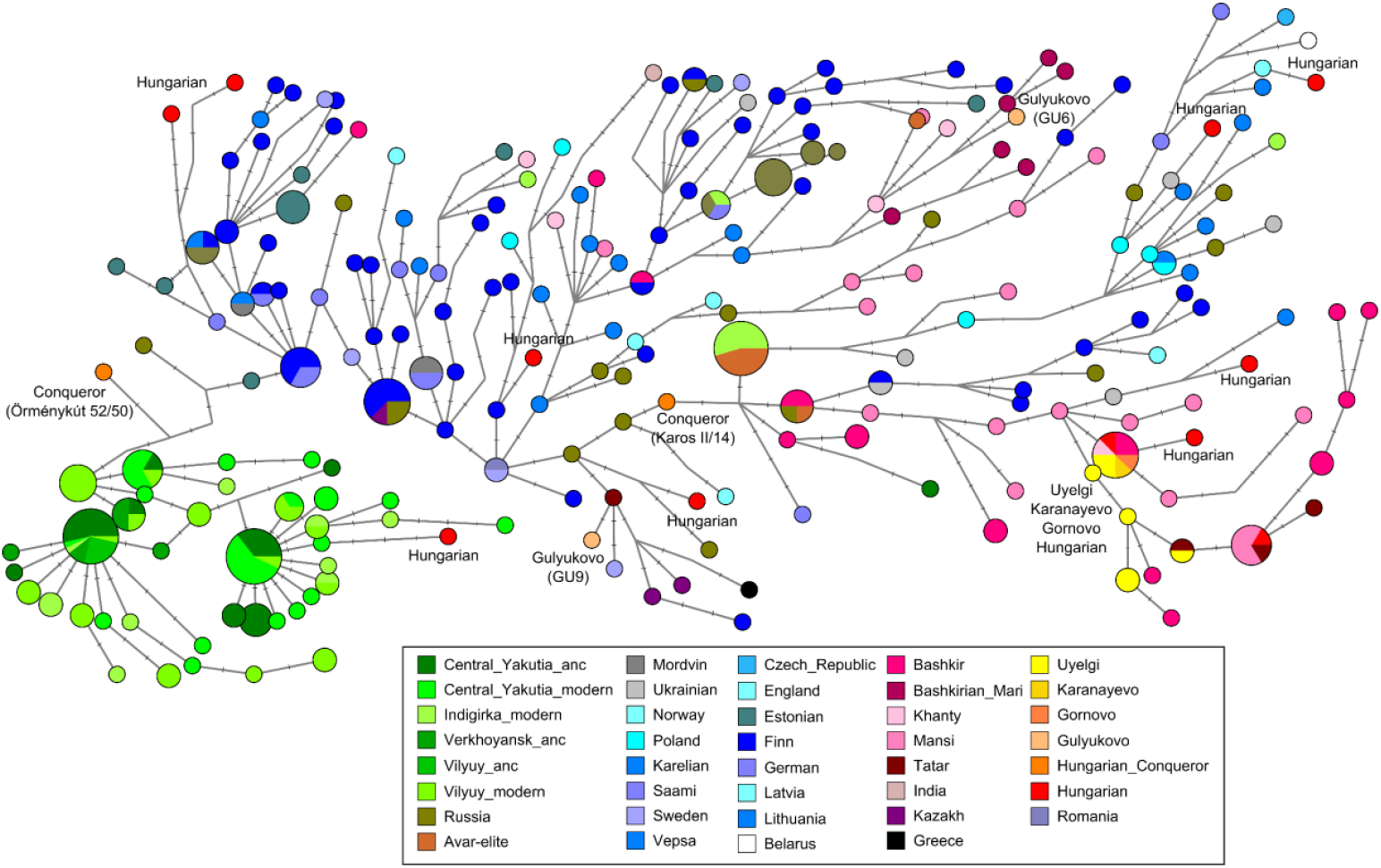
Median-joining network of N1a1 Y-chromosomal haplogroup based on 17 STRs. The studied Y lineages from Uyelgi+Karanayevo (Uyelgi and Karanayevo sites) and Chiyalik (Gornovo) groups are identical or closely related to each other and Bashkirs from Perm, Strelibashevsy, Burzyansky, and Western Orenburg regions, as well as Khantys, Southern and Northern Mansis, modern-day Hungarians, Tatars, and Ukrainians. One sample from the Chiyalik group (Gulyukovo site, GU6) is on a subbranch, mainly composed of Bashkirian Maris. The other sample from this site (GU9) is on another subbranch, which contains Tatars and Russians as well. For information on the groups and STR markers see Supplementary Material, Table S10.

On R1a networks (Supplementary Material, Fig. S40-S41), individuals from the Novinki and Bolshie Tigani groups are related to males from the Middle East, while those from the Chiyalik group are close to Russian and Belorussian males. In addition, these analyses show intra-site connections. Although the predicted haplogroup of a male from Novinki group is the same as King Béla III, based on the network analysis, there can only be a distant relationship between them (Supplementary Material, Fig. S42).

Based on the Y-STR patterns, another intra-site link was detected between two males from Bolshie Tigani sharing haplogroup I1 (Supplementary Material, Table S2).

### Mitochondrial haplogroup frequency based analyses

On the mitochondrial DNA (mtDNA) haplogroup frequency-based Principal Component Analysis (PCA) plot, the examined groups are in an intermediate position between the populations of Eastern and Western Eurasia, indicating that the Volga–Ural region served as a contact zone (Fig. 4, Supplementary Material, Fig. S36, Table S3). Because of their similar macro-haplogroup composition (Fig. 2), the Uyelgi+Karanayevo, Chiyalik, Tankeevka, Bolshie Tigani and Novinki groups are close to one another. Due to the lack of data for the region we sampled, this unit is mapped in a sparsely covered area of the plot closest to modern Turkmens, Uzbeks, Khantys, Mansis, and Iron and Bronze Age groups of Central Asia, pointing to an unexplored genetic cluster between the East and the West. The Ward type clustering supports the haplogroup level relationship among the groups we studied. Most of them fall in the same cluster, in the main branch comprising steppe groups (Supplementary Material, Fig. S37). These results demonstrate that the Iron and Bronze Age groups in the area determined the genetic composition of the territory under investigation, but since they are chronologically distant from the examined medieval groups, we cannot infer direct succession. The ancient proto-Ob-Ugric group is plotted next to the North and Central Asian unit, in the vicinity of the representatives of Iron Age and Bronze Age cultures in Russia (Okunevo, Krotovo, Central Asian Late Iron Age cultures), as well as modern North Asian groups (Nganasan, Even, Evenk), which affinities are also confirmed by the Ward analysis (Supplementary Material, Fig. S37).

**Figure 4:**
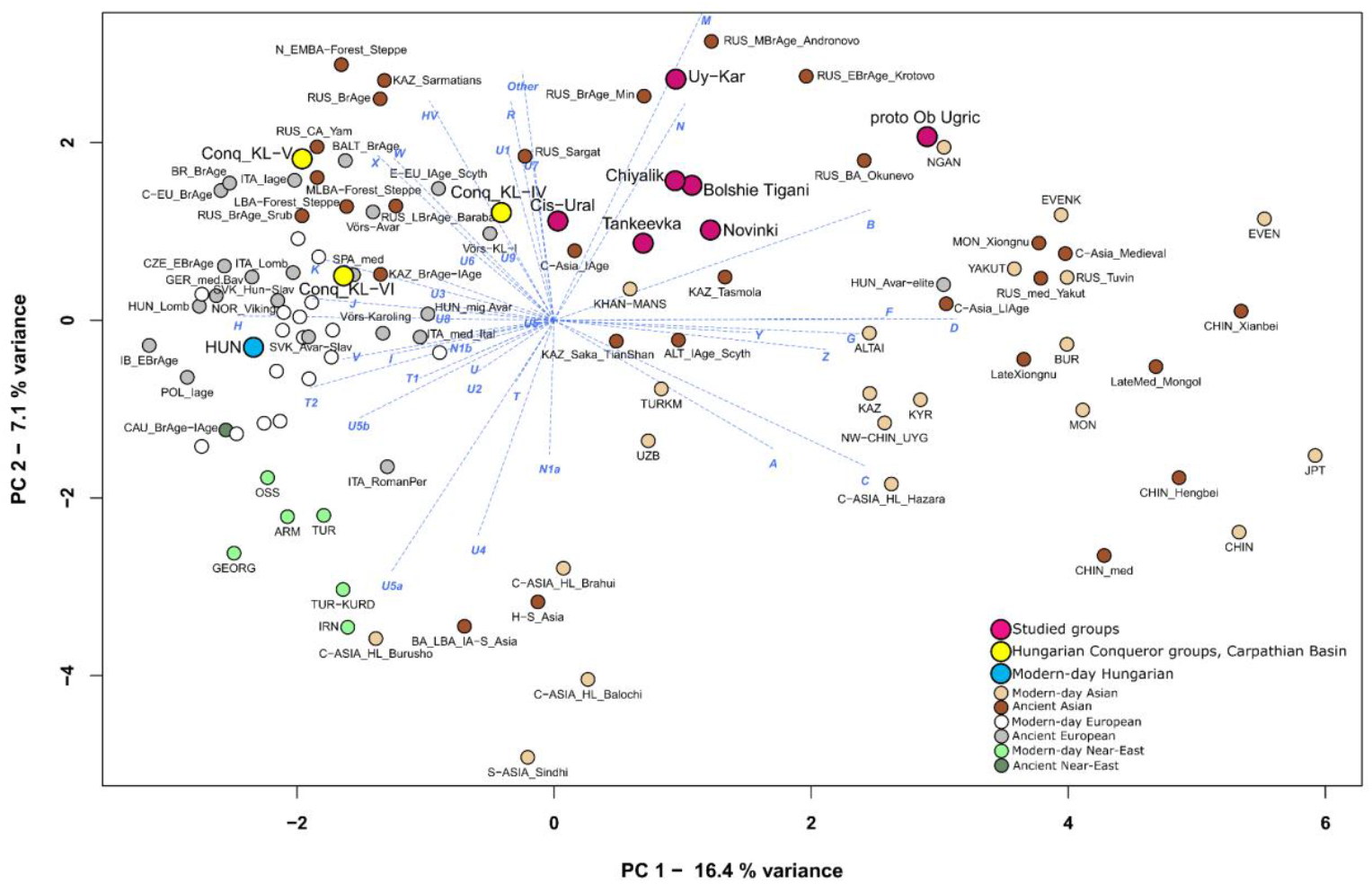
PCA plot based on mitochondrial haplogroup frequencies of ancient and present-day Eurasian and Near-Eastern populations. The variances presented on the first two components: PC1:16.4%; PC2: 7.1%. For abbreviations and information see Supplementary Material, Table S3. PC3 with a variance of 6.3% is presented in Supplementary Material, Figure S36.

The conqueror Hungarians (KL-IV, V and VI) are markedly different from one another. Group KL-IV contains more typical eastern and less characteristic western mitochondrial lines (Fig. 2). Therefore, it is the most closely related to the studied Volga-Ural region’s groups. In contrast, the KL-V-VI are located on PCA plots between the European and Bronze Age steppe groups (Fig. 4, Supplementary Material, Fig. S36).

The role of the representatives of the Iron Age Sargat culture in the development of the early Ugric people has been discussed in genetic publications (18) (45). Though this group from present-day Russia (RUS-Sargat) was close to Group KL-IV in our analyses, its small sampleset (n=18) makes conclusions precarious.

In each analysis, modern Hungarians belong to the European populations, which demonstrates their admixture with Europeans since the Hungarian Conquest (20).

### Mitogenome sequence-based analyses

The genetic distances (F_ST_) are not significant between most of the studied Volga-Ural groups (Fig. 5, Supplementary Material, Table S4) except between the proto-Ob-Ugric and Tankeevka groups. The Uyelgi+Karanayevo group however differs significantly from the others, except for the proto-Ob-Ugric and Chiyalik groups, perhaps due to the numerous intra- and intersite genetic connections, which also indicate a close relationship between the two communities (Supplementary Material, Fig. S43-44).

**Figure 5:**
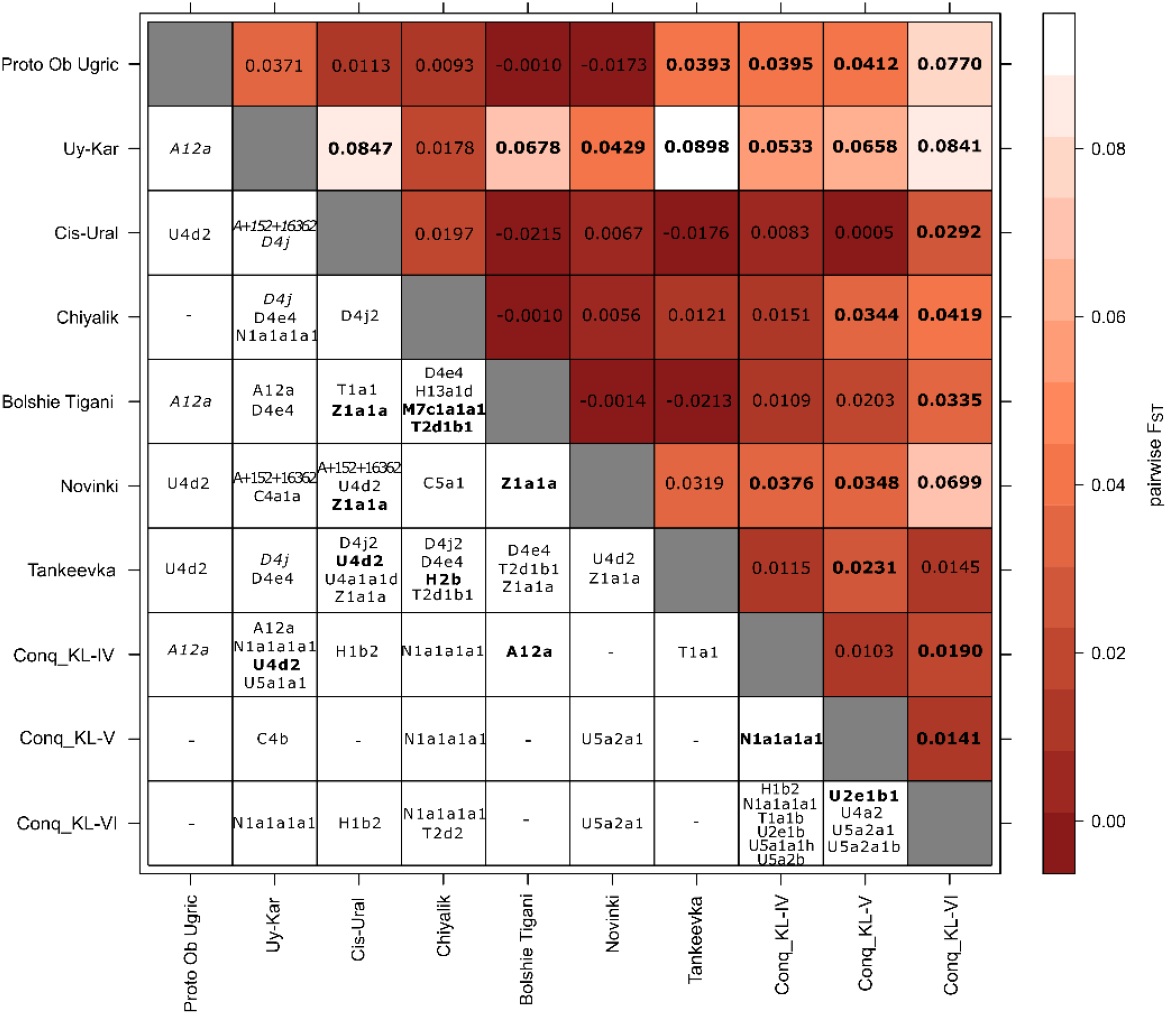
Shared mtDNA subhaplogroups between the studied groups (under the diagonal) and results of F_ST_ analysis (over the diagonal). F_ST_ values in bold indicate a significant genetic distance. In the case of phylogenetic relationships, the more distal but unambiguous relationships are indicated by italics, the normal letters marking close relationships, and letters in bolding showing haplotype identity between detected mitogenomes.

Based on the results of the Mantel Test, the genetical and geographical distances of the examined Volga-Ural region groups are not correlated, although large trends are observable on Eurasia scale. This signalizes that the diversity of the examined groups was not only the cause of spatial separation, but rather mobility and cultural ties that influenced their marital habits and maternal makeup (Supplementary Material, Chapter B, 1.6., Table S6).

The three conqueror groups are related to the Volga-Ural region groups in different ways (Fig. 5), in line with the results of the haplogroup-based analyses. The distance between KL-IV and KL-V is not significant, which is also in agreement with the similarity of the materials found in these cemeteries (39).

We performed clustering on ancient and recent Eurasian groups based on linearised Slatkin F_ST_ (Supplementary Material, Fig. S38), where ancient and recent populations form shared branches. Uyelgi+Karanayevo, Proto-Ob-Ugric and Novinki groups are in one cluster, which contains the Southern and Central Asian Iron Age, Bronze Age and recent period populations as well. Bolshie Tigani and the Chiyalik groups are also in the Southern/Central Asian context but in a cluster different from the earlier groups. Cis-Ural, KL-IV and KL-V are close to each other on a steppe origin subcluster, where the KL-V is the closest to the Sargat population, despite the previously demonstrated initial assimilation signals of the conquerors. Tankeevka and KL-VI are between the modern-day and ancient European groups. For AMOVA analysis (Analaysis of Molecular Variance, Supplementary Material, Table S5), we classified the 10 studied groups into three sets based on the clustering results: 1) Uyelgi+Karanayevo, proto-Ob-Ugric, Novinki; 2) Tankeevka, Cis-Ural, KL-IV, KL-V, KL-VI and 3) Bolshie Tigani, Chiyalik. The source of variance among the sets is 4.06% and their difference is significant (F_CT_= 0.04058, p= 0.00782+-0.00313), while the variance within the sets is 0.83% and they are not significantly different (F_SC_= 0.00869, p= 0.15836+-0.01353). These results are confident with the results of earlier analysis and with Multidimensional Scaling plot as well (Supplementary Material, Fig. S39).

### Mitogenome phylogenetic analyses

Phylogenetic analysis of maternal lineages informs about the diverse connections of the Hungarians in the Carpathian Basin and the studied Volga-Ural region groups. (Supplementary Material, Table S8, Chapter B, 2.2, Extended Figure 4). During the analysis we explored many close intra-site maternal relationships as well.

We made neighbour-joining phylogenetic trees from those haplogroups separately, which were detected in at least two groups associated with the Hungarians (including the Uyelgi, Cis-Ural, and KL-IV–VI groups, see Supplementary Material, Table S8., Chapter B, 2.2). Inter-site haplotype (sequence) identities testify to close relationships between the studied groups suggested also previously by archaeological finds (Supplementary Material, Fig. S43-S44). We found the most numerous examples of this between the Bolshie Tigani and Chiyalik groups (within haplogroups M7c1a1a1, T2d1b1) (Supplementary Material, Fig. S52 and Fig. S56), which is consistent with both haplogroup frequency and F_ST_-based analyses.

The phylogenetic trees of several haplogroups clearly demonstrate the existence of related maternal lineages among the groups under investigation, supporting their assumed connection (Fig. 5 and Supplementary Material, Chapter B, 2.2). Bolshie Tigani, Tankeevka and Chiyalik groups have the largest number of phylogenetic links. Further, Cis-Ural and Tankeevka, as well as Chiyalik and Uyelgi+Karanayevo groups reveal many closely related lineages (Supplementary Material, Fig. S44). From the conquerors, KL-IV is connected with the largest number of maternal lines to the studied groups of the Volga-Ural region (especially to the Uyelgi+Karanayevo) confirming the putative genetic relatedness behind the similarity of the archaeological finds.

The finding of many phylogenetic links between significantly different groups suggests that although the chronological difference and geographical distance, or possible admixtures with different groups transformed the overall genetic picture, common lineages were preserved at the individual level in the descendants. This can be observed, for example, between Uyelgi+Karanayevo and KL-IV, between KL-IV and KL-VI, as well as between KL-V and KL-VI (Fig. 5, Supplementary Material, Fig. S43). Within the haplogroup H1b, the mtDNA of King Béla III showed no connection with the newly analysed mitogenomes; nevertheless, his mitochondrial sequence has the closest relation to an individual from the Carpathian Basin KL-VI group (Supplementary Material, Extended Figure 4).

The majority of maternal lines can be traced back to Central Asia (south of the line connecting the Caucasus and present-day Kazakhstan; e.g., C4b, C4+152, U4a2) and the steppe areas (the forest-steppe and steppe grassland north of the former area through the Altai Mountains to Lake Baikal; e.g., C5c, C4a1a+195) according to phylogenetic analyses (Supplementary Material, Chapter B. 2.2). Several maternal lines of Siberian (e.g., A8a1, D4j4, T2d1b1, Z1a1a) and Middle Eastern (e.g., U3a, U3b) origins have also been detected, while some others point to the Far East (e.g., D4g1b, M7c). This diversity is expected based on migrations going through Central Eurasia (17) (18) and is in line with the results of the population genetic analysis. Several maternal lineages represent a direct link between the groups from the Volga-Ural region and Carpathian Basin (e.g., A12a (Supplementary Material, Fig. S45A), N1a1a1a1a (Fig. 6 and Supplementary Material, Fig. S53), which proves that these groups most probably had a common source in (or near) the study region. The subhaplogroup N1a1a1a1a is particularly common among the Hungarian related groups (with a prevalence of 26% in the Uyelgi+Karanayevo, 12% in the Chiyalik group, 9.57% in KL-IV, and 5.26% in KL-V) and the structure of the mtDNA phylogenetic tree suggests extended and almost exclusive maternal connections within and among these studied groups (and a representative of KL-VI). Based on the N1a phylogeny, the Kushnarenkovo and Chiyalik/Hungarian conqueror split can be dated to 600-750 AD, while the split date of ancestors of the Chiyalik and conquerors (9^th^ century) coincides with the Hungarian conquest.

**Figure 6:**
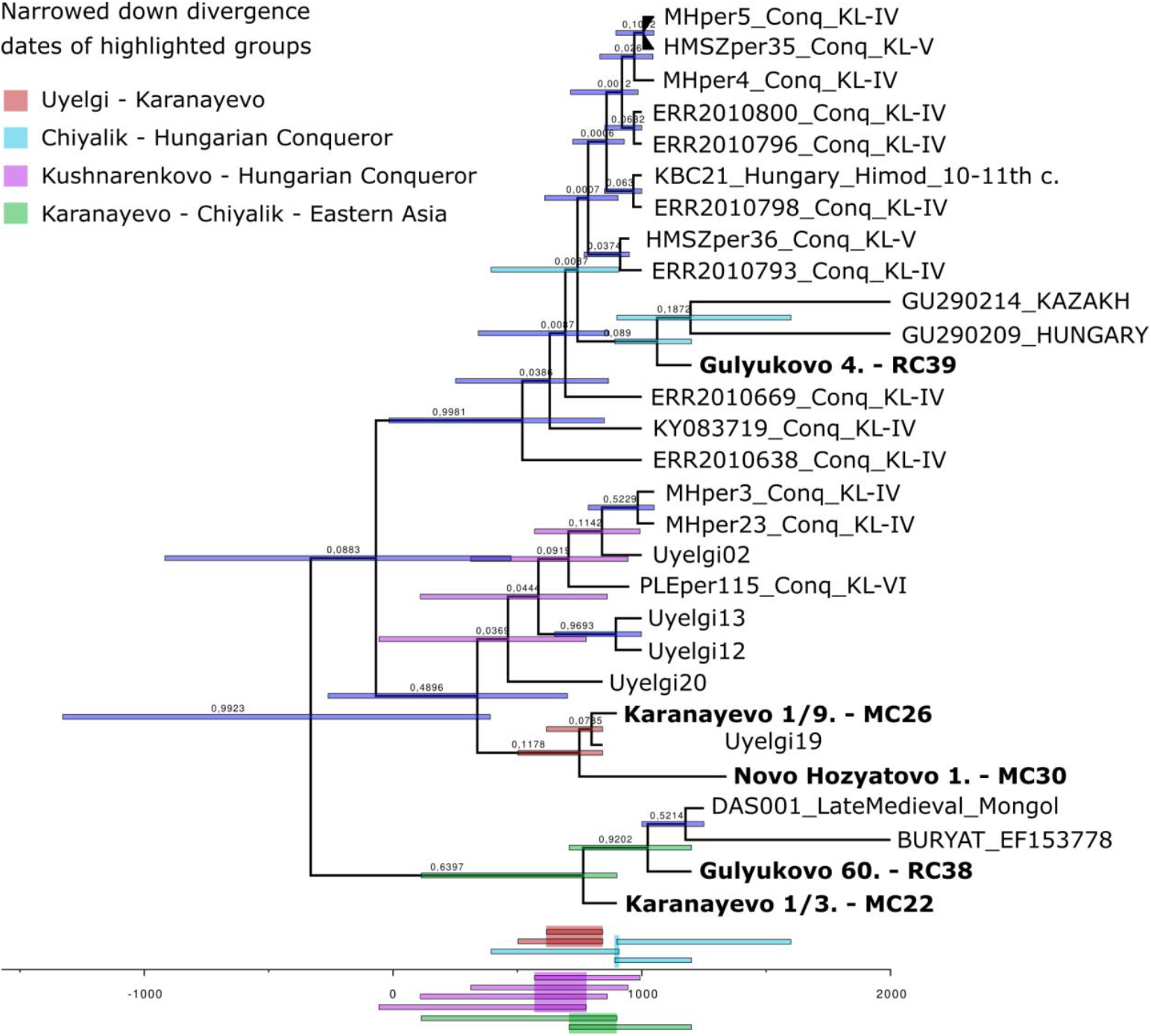
Mitochondrial haplogroup N1a1a1a1a phylogenetic tree with divergence dates, made with BEAST software. This tree mainly consists of samples linked to Hungarian prehistory, including both database and data of this study (in bold). Divergence date estimations correlate with presumptions of group split times based on historical and archaeological data. The majority of the group divergence dates are 600-800 AD, pointing to rapid population movements in this period. For the complete tree, see Supplementary Material, Extended Figure 3.

This latter observation suggests a rapid movement from the Volga-Ural region to the Carpathian Basin that might have taken place within a generation.

### Discussion

This study reveals the first genetic data of the early medieval sites in the Western-Siberian and Volga-Kama regions, in addition to our previous publication from the Ural region (22). These study regions are extremely important in the ethnogenesis of several Uralic and Turkic peoples, including the Hungarians too; however, only a few of these populations have been genetically studied so far.

Our analyses indicate little or no biological connection between the ancestors of Hungarians and proto-Ob-Ugric groups in Western Siberia, despite their close geographical proximity for 1500–2000 years after their split estimated by linguistic models and chronology (9). We identified only a few uniparental links between them, but our results also show that it is necessary to further study the proto-Ob-Ugric peoples as well as nomadic peoples that arrived in Western Siberia from the south and east in several waves, and who provided local ancestral components of later Western Siberian populations.

On the other hand, these new results support the proposed intensive relationship network between the eastern and western Uralic communities in the 9^th^–11^th^ centuries (Uyelgi and Karanayevo sites) in the late Kushnarenkovo culture (46) (47) (48). We detected closely related and identical sequences and haplotypes both in the case of the maternal and paternal lineages. Further, the estimated divergence time of the mitochondrial haplogroup N1a1a1a1a also supports their close connections.

The western sites of the Uralic Kushnarenkovo and Karayakupovo cultures located along the Rivers Volga and Kama (Bolshie Tigani and Tankeevka) show individual genetic links to each other, although population genetic results approximate Tankeevka to the European groups. According to F_ST_ analysis, both groups are similar to Volga-Tatars, who live today in the studied region, suggesting that the base population of the Volga-Ural region still has its imprint on the present-day groups of the region.

Together with historical data, our results clearly connect the population of Bolshie Tigani cemetery (9^th^–10^th^ centuries) to the representatives of the sites of the later Chiyalik culture (11^th^–14^th^ centuries), as we detected numerous (identical) maternal lineages between them. Based on the population genetic analyses, the two groups had elements of common origin, which suggest at least partial continuity of the population, but the chronological gap also explains diverging affinities.

The burials of Novinki-type sites are archaeologically attributed to the representatives of Bulgar and/or Khazar heterogeneous (presumably border guarding military) groups. The genetic links of this group with Central and Inner Asia are in line with the historical and archaeological facts that the Khazars came from the territory of the Western Turkic Khaganate (49). Based on the genetic, historical and archaeological data (50), it is plausible that the members of this community came into indirect or even direct contact with the Hungarians or other Permian (Cis-Uralic) people (e.g., as mitochondrial haplogroup Z1a indicates) living in their geographical neighbourhood.

We re-classified the burial grounds of the Hungarian Conquest period in the Carpathian Basin using archaeological cemetery typology (39) (40). Phylogenetic observations suggest that (at least a part of) the conquerors separated from the representatives of the Kushnarenkovo culture 600-750 AD, while the split between the ancestors of the Chiyalik group and the conquerors dated in the 9^th^ century. In the Carpathian Basin, the new settlers and the local population started admixing only in the second half of the 10^th^ century. The group KL-IV (10^th^ AD), both at the individual and community levels, looks more similar to the population of the Volga-Ural region sites associated with the Hungarians. According to the archaeological chronology, this cemetery group was used by the first and second generations of the conquerors. A significant part of their uniparental lineages can be derived from the Volga and Ural regions and they show affinity to the Iron Age Sargat culture’s population, all of which suggests that they have only limited interaction with the local population of the Carpathian Basin. The cemeteries belonging to groups KL-V and KL-VI reflect increasing genetic absorption and the effect of the local population substrate, but some maternal lineages originating in the east also survived in these groups.

Based on the numerous maternal links we detected among the populations of the Volga-Ural region and the groups of conquerors, we conclude that men and women jointly immigrated to the Carpathian Basin. In the Volga-Ural region, we could identify most of the “eastern” genetic traits of the conquerors (27) (28) (29) (34) (35). Therefore, the historical, linguistic, and archaeological assumption that these population elements (i.e., a part of the conqueror Hungarians) may have originated directly or indirectly from this area has been supported by our results.

This study confirms that the conquerors, and even their predecessors living in the region of the Volga and the Southern Urals, formed a composite, mixed population (6) (9) (22) (51), and their genetic makeup was influenced by the base population of the area.

The highlighted N1a1-M46 Y-chromosomal lineage shows a genetic link between the Kushnarenkovo and Chiyalik cultures, the conquerors and modern-day Hungarians, as well as the Volga-Ural region’s present-day groups (Bashkir, Tatar, Khanty and Mansi). This lineage is another piece of evidence that (at least a part of) the Hungarians came from the Volga-Ural region, from the territory of the Kushnarenkovo and Karayakupovo cultures, and it also shows the shared genetic history of the conquerors and the recent populations of the Volga-Ural region. The shared (and also chronologically and geographically debated) history of the Hungarians and Bashkirs can only be presented here from the aspect of the paternal lines, but it is also plausible on the mitochondrial level because a part of the Chiyalik population was most probably assimilated into the Bashkirs during the Middle Ages.

To gain a full spectrum of the origins of the Hungarians, the early Hungarian cemeteries discovered in the areas west of the River Volga and the earliest Hungarians arriving in the Carpathian Basin (i.e., the actual conquerors) need to be examined in the future. The autosomal DNA analyses with higher information resolution will help understand the processes of population transformation in more detail.

## Materials and Methods

### Presentation of the examined cemeteries and sites

The Bustanaevo cemetery at the western foothills of the Southern Urals (6^th^-7^th^ century, early Kushnarenkovo culture) represents the first generations of the early Kushnarenkovo population (including the early Hungarians, according to most of the researchers) moving there from the Trans-Urals (52). We also studied the Karanayevo site (9^th^–11^th^ centuries, late Kushnarenkovo culture) located at the western foothills of the Urals, where the burial customs and grave-goods are the closest analogues of the Trans-Uralic Uyelgi site.

In the cemeteries of Bolshie Tigani (9^th^–10^th^ centuries) and Tankeevka (10^th^–11^th^ centuries) (Kushnarenkovo/Karayakupovo culture mixed with early Bulgars), a Hungarian ethnic component could be identified based on burial customs and grave-goods. They can be considered as early Hungarians who did not migrate to the areas west of River Volga (2) (53) (54) (55) (56). The Chiyalik culture (Gulyukovo, Novo Hozyatovo, and Gornovo cemeteries, 11^th^–14^th^ centuries) may have been later communities of the Hungarians who remained in the East, in an already Islamised environment. As per the Muslim religion, fewer funeral offerings were placed in the graves than before, but based on their types and shapes, they can be compared with the artefacts characteristic of the Hungarians. The Novinki-type (8^th^–9^th^ centuries) of sites (Novinki, Lebyazinka, Malaya Ryazan, Mullovka, Shilovka and Brusyany) in the Samara Bend of the Volga may be suitable for studying the former neighbours of the Hungarians and early Khazar–Hungarian relations. These sites yielded the archaeological heritage of the population (presumably consisting of artificially organised communities of Bulgar and Khazar origins) settled there to protect the most important river crossing on the eastern border of the Khaganate (49). The examined sites along the left bank of the Volga may provide important information about the migration of the Hungarians due to the archaeological finds discovered near the Urals (presumably belonging to the early Hungarians) and it is possible to compare them with their former neighbours.

The study of the proto-Ob-Ugric samples (6^th^–13^th^ centuries; Ivanov Mis, Panovo, Ust-Tara, Vikulovo, and Barshov Gorodok sites) from Western Siberia was not carried out for linguistic reasons, that is, for a common Uralic origin with the Hungarians as the ancestors of the Hungarians lived in the immediate vicinity of these Ugric-speaking peoples for 1500–2000 years, at least to the mid-sixth century (6) (10) (57), but more likely to the beginning of the 9^th^ century (2) (4). Archaeological evidence of this (e.g., ceramics of Taiga origin) can be found in many cemeteries in the forest-steppe region, even in the Uyelgi cemetery (58).

For more information about the archaeological background and the studied sites, see Supplementary Material, Chapter A.

### Sample collection

We sampled bones and teeth of 112 early medieval individuals from modern-day Russia. These findings came from 18 different burial places (dated to 6^th^-14^th^ centuries AD), which represent different cultures (Fig. 1, Table 1, Supplementary Material, Table S1.). In most cases, we extracted DNA from the petrous bone (*pars petrosa ossis temporalis*). If this part of the skull was not preserved, we used teeth and long bone fragments. We radiocarbon dated 34 samples in Poznan and Debrecen ATOMKI laboratories (Supplementary Material, Table S1), along with d^13^C and d^15^N measurements for 30 samples performed in Poznan Radiocarbon Laboratory. BP dates were calibrated using OxCal 4.4 and calibration curve IntCal20 (59) (60). We found substantial freshwater reservoir effects (61) in the case of the Western Siberian communities (having d^15^N >13 ‰ and d^13^C>-20 ‰ values) along with much older ^14^C dates than the archaeological dating (see Supplementary Material, Chapter A.9). Due to this uncertainty, we followed the original archaeological chronology in all cases and present radiocarbon dates only as supplementary information (Supplementary Material, Chapter A, Table S1, and Table S11).

### Laboratory conditions

We worked in spatially separated pre- and post-PCR laboratories. The pre-PCR laboratory is a dedicated ancient DNA laboratory (Institute of Archaeogenomics, Research Centre for the Humanities, Eötvös Loránd Research Network). In this, every workflow was performed in separate rooms under sterile conditions, following well-established ancient DNA workflow protocols, as described by Csáky et al. (62). We determined the mitochondrial and Y-chromosome haplotype of the laboratory staff and compared these data with the ancient bone sample results.

### Preparation of bone fragments and teeth

We bleached the surfaces of the bones and teeth and took photos of them, thereafter we irradiated them with UV-C light and gained powder from the bone fragments and teeth. We used two different methods for this: the surfaces of long bone fragments and teeth were cleaned with sandblasting and mechanically ground into fine powder in a mixer mill. This method was also used for 53 petrous bones (63). In the case of the other 40 petrous bones, the powder was gained by direct drilling under sterile conditions (64).

### Ancient DNA extraction and DNA library preparation

The DNA extraction was performed according to the protocol of Dabney et al. (65) with minor changes (63) from 80-100 mg of bone powder. The success of the DNA extraction was verified by PCR reaction (63).

During DNA library preparation, we worked according to the protocol of Rohland et al. (66) with small changes. Half-UDG (Uracil-DNA-Glycosylase) treatment was used for most of the samples, except for 20 samples where no-UDG treatment was applied (Supplementary Material, Table S1). We used unique P5 and P7 internal barcoded adapter combinations for each sample. The DNA libraries were amplified (TwistAmp Basic – Twist DX Ltd), purified (AMPure XP beads (Agilent)), and the concentration and fragment size checked (Qubit 2.0, Agilent 4200 TapeStation System (Agilent High Sensitivity D1000 ScreenTape Assay)).

### Hybridization capture and Next Generation Sequencing

We used the hybridisation capture method to selectively enrich the entire mitochondrial genome. Y-chromosome SNP capture (63) (67) (564 SNPs on the Y-chromosome) was also used for 22 individuals, who had been previously genetically defined as male (Supplementary Material, Table S2, and Table S10). The bait production and amplification method were as described by Csáky et al. (62). We provided the captured as well as raw libraries for the shallow shotgun sequencing with unique iP7 and universal iP5 indexes for multiplex sequencing (68). We used the Illumina MiSeq device for the NGS sequencing with Illumina MiSeq Reagent Kit v3 (150-cycles) and Illumina MiSeq Reagent Kit v2 (300-cycles).

### Bioinformatic analysis

We used the same in-house pipeline for read processing as described in Csáky et al. (22), with minor changes (D. Gerber et al. 2022, manuscript in preparation) (for results see Supplementary Material, Table S2). The bam and FastQ files were uploaded to ENA (https://www.ebi.ac.uk/ena/browser/home) under the accession number: PRJEB49842. Haplogroup determination for mitochondrial DNA (mtDNA) was performed by Haplogrep 2 (69) on fasta files that were called with a custom R script suited for archaic DNA.

Contamination levels of the mtDNA were estimated using the ContamMix software (70), for results, see Supplementary Material, Table S2. Certain samples without UDG treatment show higher levels of - false positive - contamination, but our approach on variant call likely eliminated most of the noise for the relatively high coverages.

### Y-chromosome SNP and STR examination

The Y-chromosome SNP analysis was described by Csáky et al (22) and classification was performed according to ISOGG v15.73. Y-chromosome profiles were determined by combining the results of STR and SNP data by using both nevgen.org and yleaf v1 software (71).

We investigated the short tandem repeats (STRs) of the Y-chromosome using the AmpFlSTR® Yfiler® PCR Amplification Kit (Applied Biosystems) and the subsequent data analysis and haplotype determination was carried out with GeneMapper® ID Software v3.2.1 (Applied Biosystems) (62). We performed independently repeated reactions on samples where at least 4 STRs were determined.

The prediction of Y-chromosome haplogroup was made by nevgen.org (https://www.nevgen.org/). We accepted as a valid result a haplogroup probability ofat least 50% and we could determine at least 8 STR loci (for major haplogroups), or where we determined less than 8 loci, but the probability was more than 80% or SNP analysis confirmed the haplogroup (Supplementary Material, Table S2).

Based on the Y-chromosome STR data, we performed a Median-joining network analysis with Network v10.1.0.0 and visualised the results with Network-Publisher v2.1.2.5. (Supplementary Material, Table S9).

### Population genetic analyses

We performed different population genetic analyses, in which we compared the studied populations to several other ancient and modern-day populations.

For the PCAs, we used the mtDNA haplogroup frequencies of 62 ancient and 45 modern-day populations (Supplementary Material, Table S3). The PCAs were made in R v4.0.0. with prcomp and the results visualised in two-dimensional plots.

For Ward hierarchical clustering, we applied the same population dataset as for PCAs. Based on the mtDNA haplogroup frequencies, we used the Euclidean distance measurement method. We displayed the results as a dendrogram in R v4.0.0 with the pvclust library.

We calculated population pairwise F_ST_ and linearized Slatkin F_ST_ (72) values based on whole mitochondrial genome sequences of 3863 modern-day and 1741 ancient individuals, using Arlequin v3.5.2.2. with the following settings: we used the Tamura & Nei substitution model (73) with 10000 permutations and a significance level of 0.05. We applied 0.3 as a gamma value. Modern-day individuals were classified into 40 groups and ancient individuals into 46 distinct groups (Supplementary Material, Table S4).

To assess the correlation between genetic and geographic distances for the studied groups (Table 1), we performed the Mantel test (74) based on pairwise F_ST_ (Supplementary Material, Table S6). In groups that include multiple sites, a geographic centre was given to determine the distance from the other groups or sites. For the Mantel test we used Arlequin v3.5.2.2.

The linearised Slatkin F_ST_ values (Supplementary Material, Table S4) were used for clustering, which we calculated in Python using the seaborn clustermap function with parameters correlation distance metric and complete linkage method for calculating the clusters.

Multidimensional Scaling calculation was made based on linearized Slatkin F_ST_ values and the results were visualised in two-dimensional plots calculated on Euclidean distances implemented in the vegan library of R v4.0.0. (Supplementary Material, Table S4).

The linearised Slatkin F_ST_ values were used for AMOVA analysis in Arlequin v3.5.2.2, in which we created groups from this study, from published Uyelgi, Cis-Ural and Hungarian conqueror groups from Carpathian Basin (KL-IV, KL-V, KL-VI) (Table 1, Supplementary Material, Chapter B, Table S4, Table S5).

### Phylogenetic analyses

We drew and visualised the median-joining network of the mitochondrial genomes of our investigated groups with the PopArt program. The input file of the PopArt was made by DnaSP. For this analysis, we used 424 sequences, which contained 495 variable sites and belonged to 268 haplotypes.

To analyse maternal relationships between the newly investigated sites, we prepared neighbour-joining phylogenetic trees from those haplogroups, that were detected in at least two different sites or groups of sites (including Uyelgi, Cis-Ural, and conqueror groups) (Table 1). We used the method described by Csáky et al. (22) with a highly extended dataset, i.e., by using all the known and available sequences assigned to a certain haplogroup. Phylogenetic trees show only highlighted subbranches of interest (see Supplementary Material, Table S9 and Chapter B.2.2.).

We checked the variants in Haplogrep 2 (69) of those haplogroups that have been described in more than one individual within a site.

BEAST v1.10.4 (75) was used to estimate the divergence dates of the maternal lineage N1a1a1a1a, which has a high prevalence in the most studied groups and also has a relatively high mutation rate, which makes it suitable for such analysis. The following options were used: HKY nucleotide model, base frequencies were estimated according to nucleotide diversity, site heterogeneity model was Gamma + invariant sites where Gamma categories were set to 4. Clock type was set to random local, and we assumed a constant size model tree. We ran four MCMC of 100 million chains, which we merged afterwards with a 10M burnin and median height estimates of Bayesian posterior tree distribution. For divergence dates, we used 95% highest posterior density weights.

## Acknowledgement

We thank László Kovács and Nadin Rohland for their professional support, Dániel Budai for the coordinates of the studied sites and Viktor Szinyei and Zsóka Varga for preparing Fig. 1.

The study was carried out with the financial support of the Russian Foundation for Basic Research in the framework of project No. 18-59-23002 “The origins of the formation of the culture of ancient Hungarians. Archaeological paleoanthropological and paleogenetic aspect of the study of medieval monuments of the Southern Urals and Western Siberia”.

The research was funded by RFBR [Russian Foundation for Basic Research] No. 19-59-23006: “The problem of cultural transformations of Magyars on the way of Hungarian Conquest”.

The research was funded by RFBR [Russian Foundation for Basic Research] No. 20-49-720010. The work was partially performed according to the Basic Research Program RAS No. 121041600045-8.

The research project has been realized within the project framework entitled: Archaeology Research on the Contacts between Hungary and the East (Our Eastern Heritage, PPCU History and Archaeology Interdisciplinary Research Team; TKP2020-NKA-11), with the support of the Thematic Excellence Program, National Research, Development and Innovation Office and also within the project framework entitled: Árpád-dynasty program (IV.2).

## Abbreviations

AMOVA: Analysis of Molecular Variance
KL-IV: Group-IV based on Kovács, 10^th^ century small cemeteries of the camps
KL-V: Group-V based on Kovács, 10^th^ century cemeteries of villages with a large number of burials
KL-VI: Group-VI based on Kovács, The cemeteries of villages opened in the 10^th^ century and used until the 11^th^ and 12^th^ centuries
mtDNA: mitochondrial DNA
PCA: Principal Component Analysis
SNP: Single Nucleotide Polymorphism
STR: Short Tandem Repeat
UDG: Uracil-DNA-Glycosylase

## Author contribution

Designed the study: B.G.M., A.T., A.Sz-N. Performed the experiments: B.Sz.

Analysed the data: B.Sz., D.G., V.Cs.

Collected archaeological and anthropological material: D.A.S., A.A.H., A.G.S., I.R.G., E.V.V., N.P.M., A.S.Z., A.V.S., K.G.K., V.V. I., B.A.K., F.A.S., A.G.K., S.G.B, I.G.V., B.G. M, A.T., O.K.

Wrote the paper:

B.Sz., D.G., P.L., B.E., A.T., A.Sz-N.

