## Supplementary Material for "Tracing genetic connections of ancient Hungarians to the 6-14^th^ century populations of the Volga-Ural region"

Bea Szeifert, Dániel Gerber, Veronika Csáky, Péter Langó, Dmitrii Alekseevich Stashenkov, Aleksandr Aleksandrovich Khokhlov, Ayrat Gabitovich Sitdikov, Ilgizar Ravil'evich Gazimzyanov, Elizaveta Valerievna Volkova, Natalia Petrovna Matveeva, Alexander Sergejevich Zelenkov, Anastasiia Viktorovna Sleptsova, Konstantin Gennadievich Karacharov, Viktoriya Vladimirovna Ilyushina, Boris Aleksandrovich Konikov, Flarit Abdulhaevich Sungatov, Alexander Gennadievich Kolonskikh, Sergei Gennad’evich Botalov, Ivan Valer’evich Grudochko, Oleksii Komar, Balázs Egyed, Balázs Gusztáv Mende, Attila Türk, Anna Szécsényi-Nagy

### Index


Chapter A: Archaeological background - Description of the archaeological sites 4

1. Bolshie Tigani cemetery 4

2. The Chiyalik culture and Eastern Hungarians from an archaeological perspective 8

3. Tankeevka cemetery 14

4. Karanayevo cemetery 17

5. Novinki-type sites 22

5.1. The site at Novinki I 23

5.2. The site at Brusyany 23

5.3. The site at Malaya Ryazan 24

5.4. The site at Shilovka 24

5.5. The site at Lebyazhinka 24

6. Proto-Ob-Ugric group 30

6.1. Nizhneobskaya culture 31

6.1.1. The site of Ust'-Tara VII 31

6.1.2. The cemetery of Barsov Gorodok I 32

6.2. The Potchevash culture 33

6.2.1. Vikulovo cemetery 33

6.3. Ust'-Ishim culture 34

6.3.1. The cemetery of Ivanov Mis I 34

6.3.2. The kurgan cemetery of Panovo I 35

7. Bustanaevo 45

*8.* Classification *of 9-12$^{th}$ centuries cemeteries in the Carpathian Basin* 47

*9.* Discussion of the radiocarbon dates 48

10. References to chapter A 52

Chapter B: Genetic analyses 57

1. Population genetic analyses 57

1.1. PCA analysis 57

1.2. Ward clustering 57

1.3. $F_{ST}$ analyses 58

1.4. AMOVA analyses 60

1.5. MDS analysis based on Slatkin $F_{ST}$ values 60

1.6. Mantel test 61

2. Individual-level analyses 61
2.1. Y-STR network analyses 61
2.2. Phylogenetic analysis of mitochondrial haplogroups 64
2.2.1. Haplogroup A 65
2.2.2. Haplogroup C4 66
2.2.3. Haplogroup C5 67
2.2.4. Haplogroup D4 68
2.2.5. Haplogroup G 69
2.2.6. Haplogroup H1b 69
2.2.7. Haplogroup H2 70
2.2.8. Haplogroup H6a1 70
2.2.9. Haplogroup H13 71
2.2.10. Haplogroup M7c 71
2.2.11. Haplogroup N1a1a1a1 72
2.2.12. Haplogroup T1a1 73
2.2.13. Haplogroup T2d 74
2.2.14. Haplogroup U2e1 75
2.2.15. Haplogroup U3 75
2.2.16. Haplogroup U4 76
2.2.17. Haplogroup U5a 78
2.2.18. Haplogroup Z1a 79
3. References to chapter B 80

### Chapter A:

### Archaeological background - Description of the archaeological sites

#### 1. Bolshie Tigani cemetery

Bolshie Tigani is an emblematic site of the research of Hungarian prehistory. It is located in the Volga-Kama Region, in the district of Alekseevskiy, Tatarstan, Russia. The cemetery, unearthed in 1974, brought a breakthrough in the research of Hungarian prehistory at an international level. The excavations were carried out in 1974 and 1975, and then between 1978 and 1985 under the supervision of E. A. Halikova and A. H. Halikov. From 2017 to 2018, a ground-penetrating radar survey and verifying excavations were conducted at the site, led by Ayrat G. Sitdikov.

A total of 150 burials were discovered, the first 56 of which (i.e. the early part of the cemetery) were published in German in Budapest (CHALIKOVA–CHALIKOV 1981). The same part of the cemetery was published in Russian in 2018 (ХАЛИКОВА–ХАЛИКОВ 2018) with the original images, but it was a revised version completed with several studies. The site had shaft graves directed west–east. The Hungarian analogues of the funerary eye plates, sabres, and especially the head and hooves of horses placed at the feet of the dead were immediately recognised by the researchers. The metal finds reflect Saltovo/Bulgar influence. Additionally, the presence of a Uralic population is suggested by the ceramic finds of the Kushnarenkovo culture, but even more so by those of the Karayakupovo culture. Based on these, researchers associated the site with the latter culture.

E. A. Halikova, the supervisor of the excavations, interpreted the cemetery as one of the sites of Hungarians migrating westwards (HALIKOVA 1976). Her theory was refuted by V. F. Gening, mainly on the grounds that the Kushnarenkovo type of pottery is not present among the Hungarian artefacts of the 10th century Carpathian Basin (GENING 1977). In his monographs, István Fodor also mentioned the cemetery (FODOR 1977, 2015). In his opinion, the population leaving Bolshie Tigani behind was in contact with the Hungarians, but he did not take part in the migration of the Hungarians to the west. Fodor dated the migration itself in the early 8th century but did not discuss in detail what evidence he based his view on. In 1980 and 1981, A. H. Halikov excavated burials belonging to a later period of the cemetery, including grave No. 65, which was dated to the first half of the 10th century by a dirham minted in 900. As this date is later than the Hungarian Conquest of the Carpathian Basin, it prompted Halikov to correct E. A. Halikova's earlier theory, assuming that Hungarians were still present in Bolshie Tigani after the mid-ninth century. In his view, it belonged to the territory of *Magna Hungaria* situated by the River Ethil, which was discovered and reported by Friar Julian in 1236 (HALIKOV 1984).

In 2018, A. V. Komar studied the cemetery in detail. He called attention to the fact that so far it is only here that 'early Hungarian' and Saltovo-type artefacts have been found together (KOMAR 2018). All the male burials in the cemetery had Saltovo-type objects. Parts of weaponry appear in every grave. Artefacts belonging to the Saltovo circle are represented by belts, sabres, and stirrups, which corresponds to the idea of a short-lived military alliance. The finds comprise belt fittings decorated with a 'stick-form' frame, and a triple palmette or a 'mythological' scene, 'crescent' and heart-shaped fittings characteristic of the Ural Region, men's hoop earrings and women's earrings with strings of balls, plain plate bracelets, grip plates without antler bow plates, diamond and lance-shaped flat arrowheads, as well as eye-covers stitched to the burial shrouds (CHALIKOVA–CHALIKOV 1981). The graves contained hand-formed Karayakupovo-type ceramic vessels, hand-formed pottery made locally in the Kama Region, as well as wheel-thrown ceramics typical of the Volga Bulghars. The influence of the Kama Region can also be clearly observed in women's objects of wear. However, the weapons, horse equipment, and belt ornaments comprised mainly Saltovo-type items.

The population of the Bolshie Tigani cemetery was undoubtedly much more closely linked to the Khazar Khaganate and the Saltovo cultural sphere than the population of the Subbotsy-horizon (connected to 9th century Hungarians) occupying the northern foreground of the Black Sea. Provided that there is some truth to the information about *Levedia – as settlement territory of Hungarians in the 8$^{th}$ and early 9$^{th}$ century* – found in Byzantine written records, it is the cemetery of Bolshie Tigani itself that provides a compelling argument for locating *Levedia* east of the Volga and not in the North-Pontic area (KOMAR 2018).

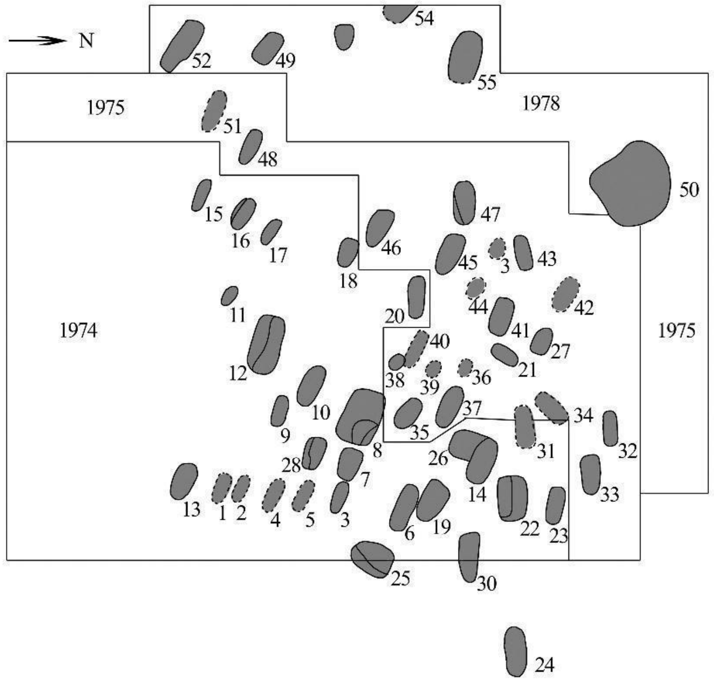


*Fig. S1. The cemetery plan of Bolshie Tigani (after CHALIKOVA-CHALIKOV 1981)*

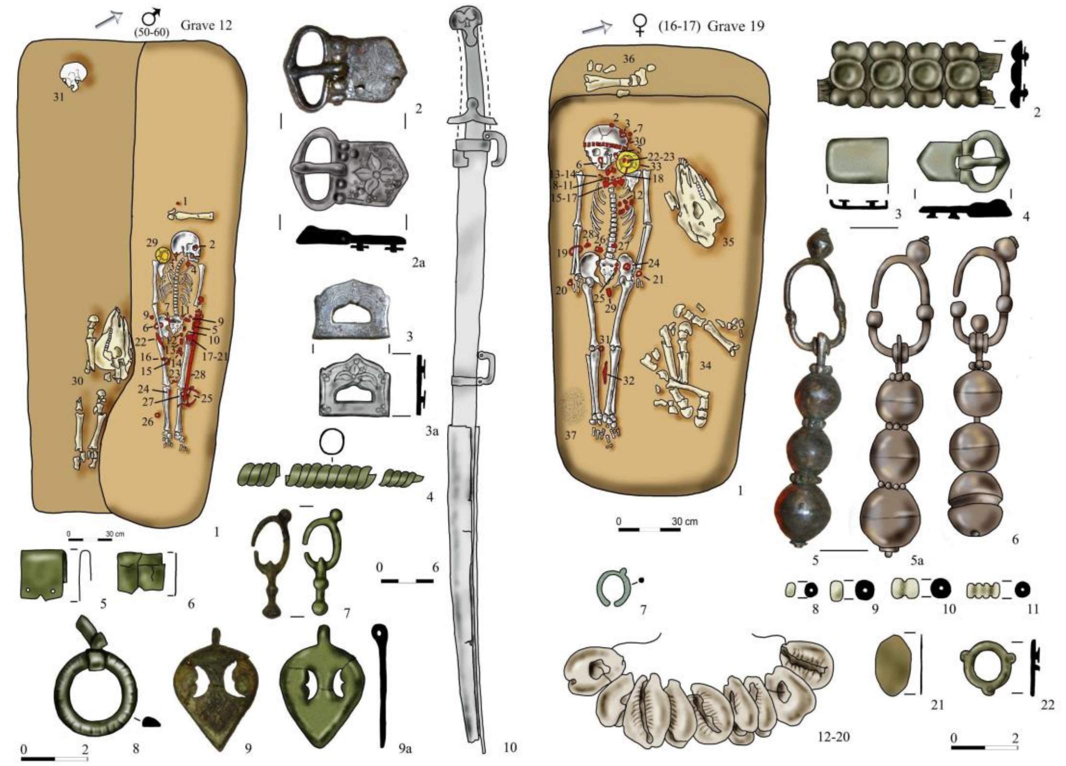


*Fig. S2. 'Hungarian'-type burials of Bolshie Tigani: Graves No. 12 and 19. (after* CHALIKOVA-CHALIKOV *1981. Photos and digital drawing by Attila Türk)*

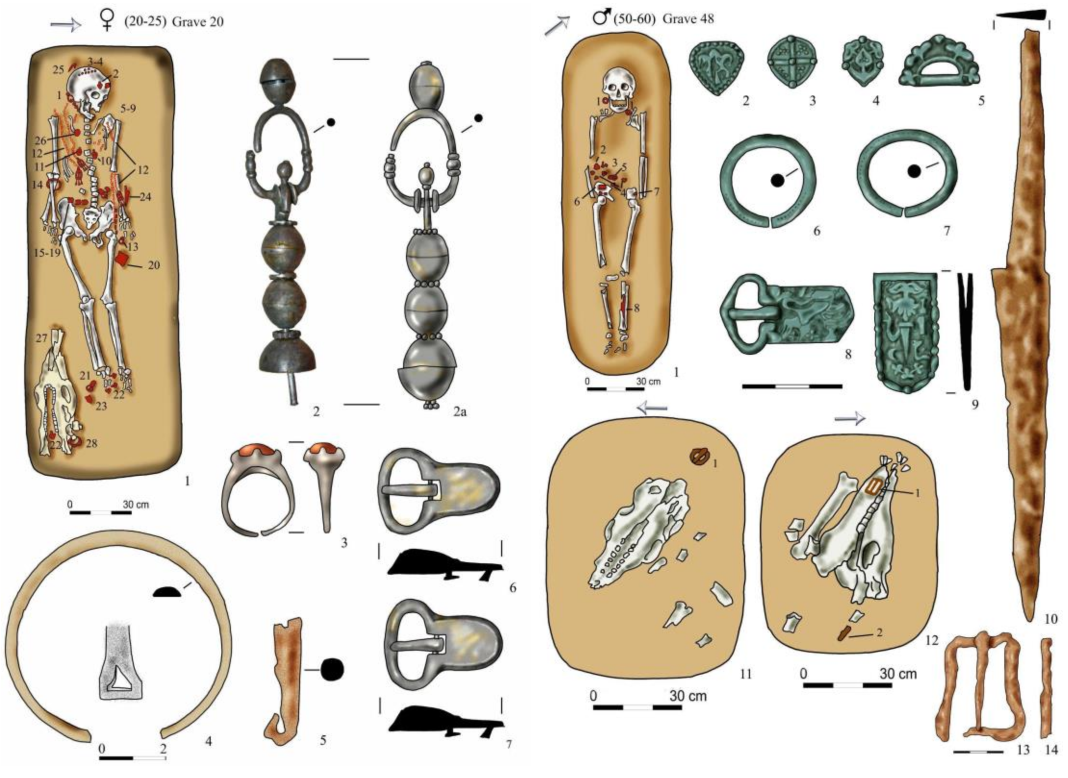


*Fig. S3. 'Bulgar'-type burials of Bolshie Tigani: Graves No. 20 and 48 (after* CHALIKOVA-CHALIKOV *1981. Photos and digital drawing by Attila Türk)*

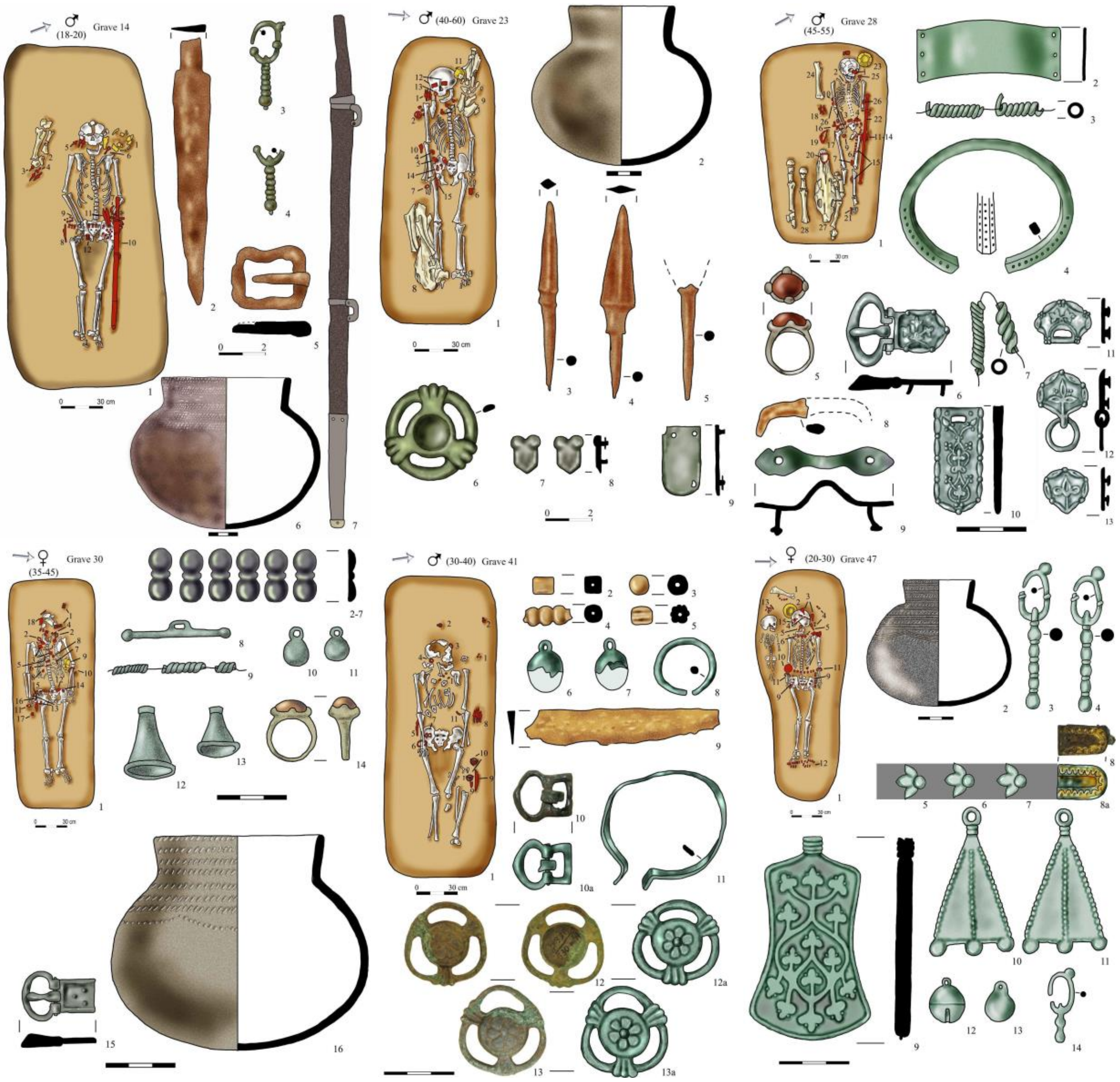


*Fig. S4: 'Uralic'-type burials of Bolshie Tigani: Graves No. 14, 28, 30, 41, 23, 47. (after* CHALIKOVA-CHALIKOV *1981. Photos and digital drawing by Attila Türk)*

#### 2. The Chiyalik culture and Eastern Hungarians from an archaeological perspective

The Chiyalik culture found on the border of Tatarstan and Bashkortostan dates from the late 9th or early 10th century to the early 15th century. It was named after the archaeological site located near the village of Chiyalik (Чиялик), in the Aktanyshsky District of Tatarstan. The excavations started there under the supervision of the Kazan archaeologist E. P. Kazakov in 1969. The Chiyalik culture was spread in the forest-steppe zone of the Southern Urals, both to the east and west of the mountains. It covered the territory of present-day Bashkortostan, Eastern Tatarstan, the south-eastern part of Udmurtia (along the River Kama), the southern part of the Perm border region (the Kungur forest-steppe area), as well as the northern part of the Chelyabinsk Region, the southern part of the Sverdlovsk Region, and the western part of the Kurgan Region. It also emerged in the forest-steppe zone of the Samara Region in the 10th century and the 13th-14th centuries.

The Chiyalik culture was identified by E. P. Kazakov in the early 1970s. In his view, it existed only from the 13th to the 14th century. It was preceded by the Postpetrogrom culture (from the 10th century), from which the Chiyalik culture later evolved. Kazakov was of the opinion that in the first centuries of the second millennium AD, two major groups of people migrated from the Ural Mountains to the foothills of the Urals. The first in the early 10th century and the second in the early 13th century. E. P. Kazakov dated the end of the culture that he called Postpetrogrom in the late 12th century. Afterwards, in his opinion, the Postpetrogrom population merged with a related group of people coming from the Urals, whom he called the Chiyalik (KAZAKOV 1987). Their other groups also merged among the Bulgarians, Udmurts, and other neighbouring peoples as an ethnic component of theirs. According to G. N. Garustovich, however, there is no evidence of any major migration of people in the region at the turn of the 12th and 13th centuries (GARUSTOVICH 1988). All the small number of new cultural elements that emerged in the late Chiyalik culture can be more readily explained by some kind of internal development.

In the 1960s, V. D. Viktorova discovered sites that had pottery with combed and corded decoration, such as the cemeteries along the Rivers Nitse and Makushino, in the central Trans-Urals, in the southern taiga zone. She defined them as Makushino-type sites, thus establishing a new concept (VIKTOROVA 1968). In Western Siberia, the deceased were buried under small kurgans in burial pits with a wooden structure. The graves are shallow, and the corpses are placed in them with the head to the west. The exploration of the funeral rite revealed the cultic role of fire and the horse. The grave goods are mostly implements and work tools, the instruments of daily life, and costume decoration. Although the Makushino-type of ceramic vessels with corded decoration were tempered with sand, researchers from Yekaterinburg later admitted that the Makushino-type sites were closely related to the Chiyalik culture (VIKTOROVA-MOROZOV 1993). In the 1970s, N. A. Mazhitov identified similar sites in Bashkortostan, the cemeteries of Karanayevo and Mryasimovo, which he called Mryasimovo-type sites (MAZHITOV 1977, 1981). The 10th-14th-century sites characterised by pottery with combed and corded decoration are associated with the Chiyalik culture in both the Trans-Ural and Cis-Ural regions, and are now interpreted as the archaeological legacy of the Bashkirs and the Eastern Hungarians living among them (KAZAKOV 2007).

Archaeologist G. N. Garustovich from Ufa agreed with Kazakov's opinion about the Chiyalik culture and differentiated nine regional groups within that. According to him, these regional groups each formed the centre of a major territorial unit, some of which were located at the western foothills of the Urals, while others were found along the Samara section of the Volga (GARUSTOVICH 1988).

The Chiyalik culture is, therefore, generally dated between the late 10th and early 15th centuries. Within this period, researchers have also distinguished between an early Chiyalik (Mryasimovo phase; late tenth century–thirteenth century) and a late Chiyalik (late thirteenth century – early fifteenth century) period.

Around the late 9th and early 10th centuries, a new population migrated from the east to the forest-steppe part of the Cis-Ural region. According to E. P. Kazakov, it was not until the late 10th century that the producers of the Postpetrogrom pottery had arrived in the land of the Bulgars from the area of Petrogromskaya culture, in the Central Urals (i.e. the forested and mountainous area stretching on the eastern side of the Ural Mountains). Before the Mongol era, they occupied a vast area around the Urals and the River Volga (KAZAKOV 1989).

Research has long been concerned with pottery with the combed and corded decoration of Ural origin, which emerged in the land of the Volga Bulghars. This is referred to as Postpetrogrom, Mryasimovo, or early Chiyalik type of pottery. T. A. Hlebnikova classified Postpetrogrom ceramics into group VII of the pottery of the Volga Bulghars. In her opinion, this type of ceramics went back to the Nevolino Culture along the River Kama. In her system of pottery typology, she also classified a hybrid piece of pottery into group VIII, which she originated from the Kushnarenkovo culture (HLEBNIKOVA 1984). However, E. P. Kazakov later formulated his thesis that, in terms of its origin, the Postpetrogrom earthenware is related to the pottery with combed and corded decoration found around the Urals. As it is suggested by its name, E. P. Kazakov derived the Postpetrogrom culture from the Petrogrom culture that existed on the eastern side of the Urals in the first millennium (KAZAKOV 1987), although he had also called attention to research gaps. V. A. Mogilnikov, who presented the culture, described only its general characteristics and dated it between the tenth and thirteenth centuries (MOGILNIKOV 1987). Researchers focusing on the region of the Urals (e.g., V. D. Viktorova, V. M. Morozov) later described the Petrogrom culture in detail and linked it to the early Ugric population (MOROZOV 2004, KAZAKOV 2007).

The early graves are characterised by partial horse burials in addition to funerary furnishing. (The horse's skull and lower part of the legs, the phalanges, and metatarsals were placed in the grave, sometimes together with the femur, and infrequently there are also complete horse burials). According to G. N. Garustovich, it is typical of these graves that the equestrian gear (one of the stirrups, the saddle, and the horse bits) were placed under the head or at the feet of the deceased, while his weapons and the pottery were place next to their head (GARUSTOVICH 1988). It needs to be emphasised that the influence of Islam on burial customs can already be observed during the early phase of the Chiyalik culture, which became more dominant over time. By the 12th century, Muslim customs had become particularly strong, as can be seen in the case of the Karanayevo and Mryasimovo kurgans (in Bashkortostan). The same can be noticed in some cemeteries without kurgans, such as Gulyukovo, Kushulevo, and Ust-Kiserty.

The most distinctive cultural feature of the semi-nomadic population in the forest-steppe zone was their earthenware with combed and corded decoration. The typical type of pottery of the early Chiyalik culture (or Postpetrogrom or Mryasimovo phase) has a round bottom and is formed like a beaker. It can be clearly seen where the neck of the vessel was attached to its cylindrical body. Crushed shells or talc were mainly used for tempering. Later, unique and hybrid pottery emerged among Chiyalik ceramics in the territory of Volga Bulgaria at the influence of Bulgarian pottery making. In terms of their shapes and decoration, these are related to Postpetrogrom ceramic vessels, but the material used for the tempering of the pottery is sand instead of shells.

The second phase of the Chiyalik culture is characterised by the spread of the Islamic funeral rite (PASTUSHENKO 2011). The late Chiyalik period can be dated to the 13th and 14th centuries. As E. P. Kazakov pointed out, the Uralic/Ugric peoples of the late Chiyalik culture preserved many pagan cultural elements and their characteristic type of pottery is hand-formed and round-bottomed with combed and corded decoration. However, towards the end of the culture's existence, this population became predominantly Turkic and embraced Islam.

Ceramics play a major role in archaeological research here, as well. The round-bottomed or bowl-shaped vessels are hand-made and unevenly burnt on a fireplace. Their colour ranges from yellow to black, and their main tempering material is already sand. The rim is often rounded and less often decorated than that of the Postpetrogrom vessels. The neck of the ceramics has a corded decoration. Below, where the neck and the body of the vessels meet, they bear a combed pattern or incised and stamped motifs (zigzag, herringbone, fringed line, and horseshoe motifs). In the final phase of the culture, undecorated earthenware became increasingly common. In the 14th century, local, hand-made ceramic vessels ceased to be used and they were replaced by imported pottery, as well as wood and leather storing vessels (Fig, S6-S7).

E. P. Kazakov distinguished two horizons within the 13th and 14th-century burials of the Chiyalik culture. He observed that among the Muslim-type of graves, there was some change in the funerary rite. Based on this, he divided the burials into early and late groups. In the early phase, the graves were oriented to the south-east, and the deceased were laid out in an extended supine position in the burial pits. In the early phase, some deviation from Islamic laws can be observed in the graves. It is common that the bodies were laid in a supine position and not facing Mecca, and grave goods were also placed in the burial pits. In the late phase, Islamic norms prevailed. The dead were placed in the grave with the head to the west, sometimes turning slightly to the north, and their body was turned a little on the right side. The dead were facing Mecca and the burials had no furnishing (PASTUSHENKO 2011, KAZAKOV 1987). The end of the Chiyalik culture can be dated to the end of the 14th century and the beginning of the 15th century. We have very little information after this time. The period between the 15th and 17th centuries is still very little researched (KAZAKOV 2007).

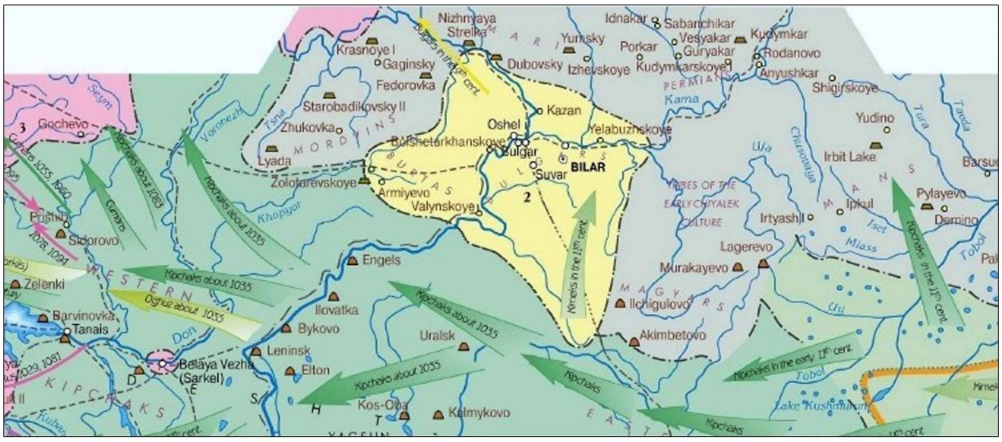


1

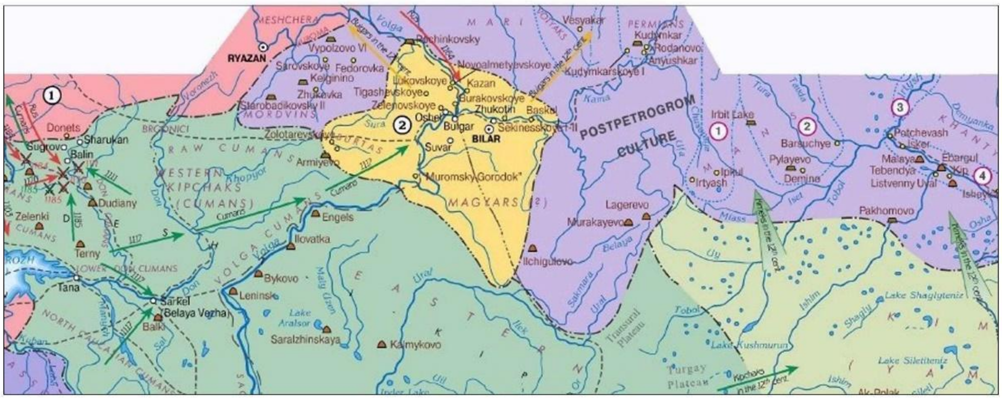


2

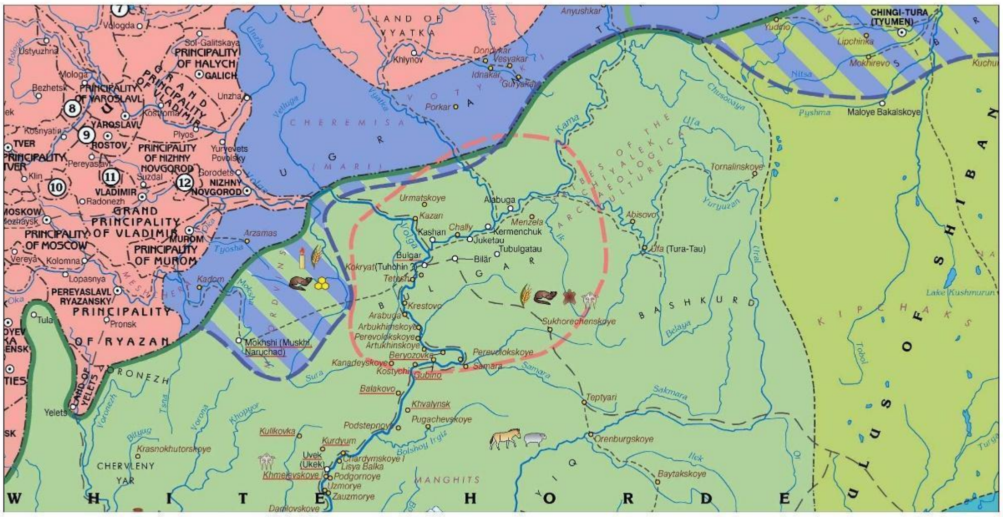


3

*Fig. S5. Changes in the presumed location of the remaining Hungarians in the East (Julianus' Hungarians) in the Russian literature, based on written sources and the Chiyalik archaeological culture (after* RAHMANALIEV *2009. 1: 11th century; 2: 12th century; 3: 13th–14th centuries. Source of the maps: https://historylib.org/historybooks/Rustan-Rakhmanaliev_Imperiya-tyurkov--Velikaya-tsivilizatsiya/).*

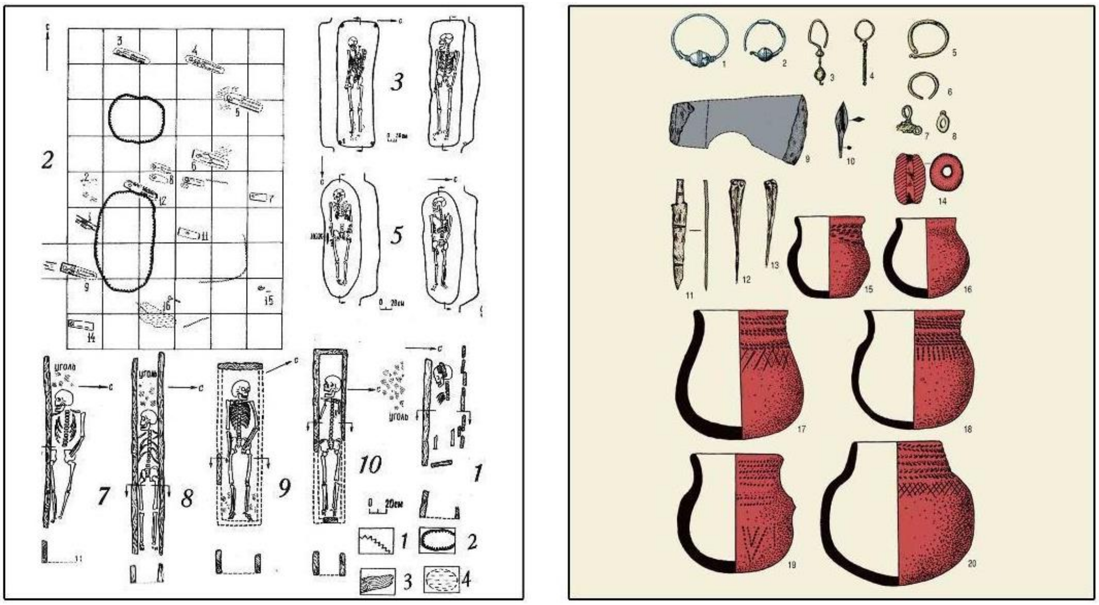

*Fig. S6: Characteristic burials and finds of the Chiyalik culture (after* GARUSTOVICH *1988)*
*Source of the pictures: https://cyberleninka.ru/article/n/chiyalikskaya-arheologicheskaya-kultura-epohi-srednevekovya-na-yuzhnom-urale*

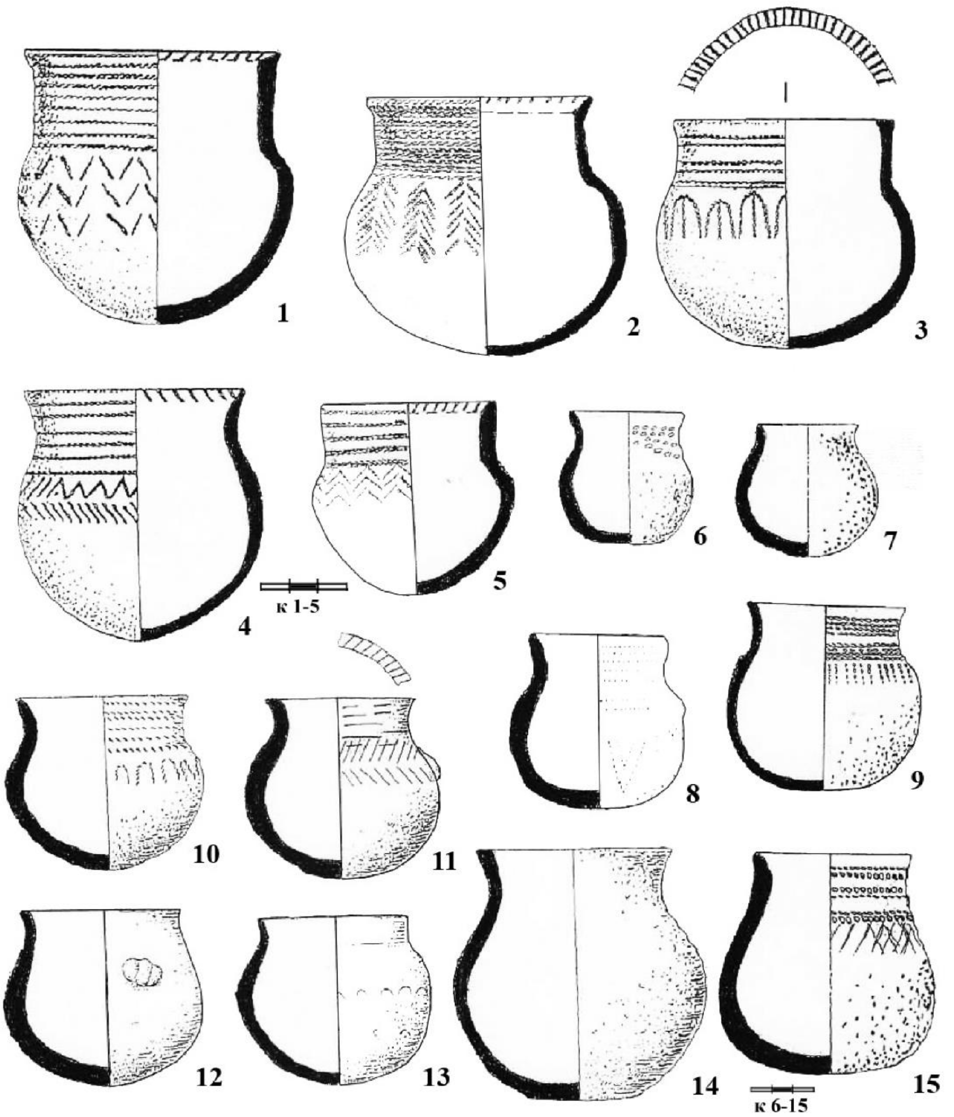


*Fig. S7. Characteristic pottery ware of the Chiyalik culture (after GARUSTOVICH 1998) Source of the pictures: https://cyberleninka.ru/article/n/chiyalikskaya-arheologicheskaya-kultura-epohi-srednevekovya-na-yuzhnom-urale*

#### 3. Tankeevka cemetery

The Tankeevka cemetery lying on the left bank of the River Volga in Tatarstan was discovered in 1904. Before the first major excavations conducted in 1961 and 1962, only stray finds were known from here. Still, researchers almost immediately observed the connection with the 10th-century Hungarian conquerors. The extensive excavations revealed that the cemetery comprising about 1,300 burials belonged to two groups of people, the Volga Bulgarians and the (Ugric) population of the Kama Region (KAZAKOV 1971, 2007, KHALIKOVA-KAZAKOV 1977). The excavation of the cemetery stretching on the eroding embankment of River Staraya Rtivina continued in 2018.

In 1977, E. A. Halikova and E. P. Kazakov published a significant part of the Tankeevka cemetery. In addition to the main Volga Bulgar population, a Hungarian ethnic element was identified (KHALIKOVA-KAZAKOV 1977). Those west–east oriented inhumation burials belonged here where a folded horsehide was placed at the feet and the horse's skull was turned towards the deceased. Another important factor is the covering of the deceased's face (HALIKOVA 1972), which is related to the Subbotsy-type finds (KOMAR 2018), as well as to the eye and mouth plates stitched on the burial shrouds of the Hungarian conquerors (FODOR 1972).

The Tankeevka wheel-thrown pottery vessels are most closely related to 10th- and 11th-century earthenware known from Volga Bulgaria, while the metal finds comprise only a few items with Saltovo-type decoration. All these show that the Tankeevka cemetery has a much later chronology than the Bolshie Tarhani cemetery, and even the Bolshie Tigani cemetery. Among the finds of the Tankeevka cemetery, the Saltovo V horizon is less represented, which may be explained by the fact that the relations between the Volga Bulgarians and the Khazars ceased in the early 10th century. At the same time, the 'post-Khazar' horizon dated after the fall of the Khazar Khaganate can be clearly observed in the last third of the 10th century. As Aleksey Komar pointed out, the cemetery of Tankeevka was established little before the end of Saltovo III horizon, that is, somewhat later than Bolshie Tigani, and was also used until later. Two kinds of funerary rites were employed in the cemetery from the very beginning. The Turkic-type horse equipment of the Ural Region is missing from Tankeevka. Additionally, there is a difference in the production technique of Saltovo- and Subbotsy-type belt mounts (KOMAR 2018). This may be due to the fact that there are no direct imports among the finds discovered in Tankeevka, only replicas of poor quality. In other words, although the groups using the Bolshie Tigani and Tankeevka cemeteries lived near each other, they were not closely related and also had different political ties with the people of the Saltovo cultural sphere. The role of the local population of the Kama Region in the development of the culture that the Tankeevka cemetery belonged to was particularly stressed by R. D. Goldina (GOLDINA 2013).

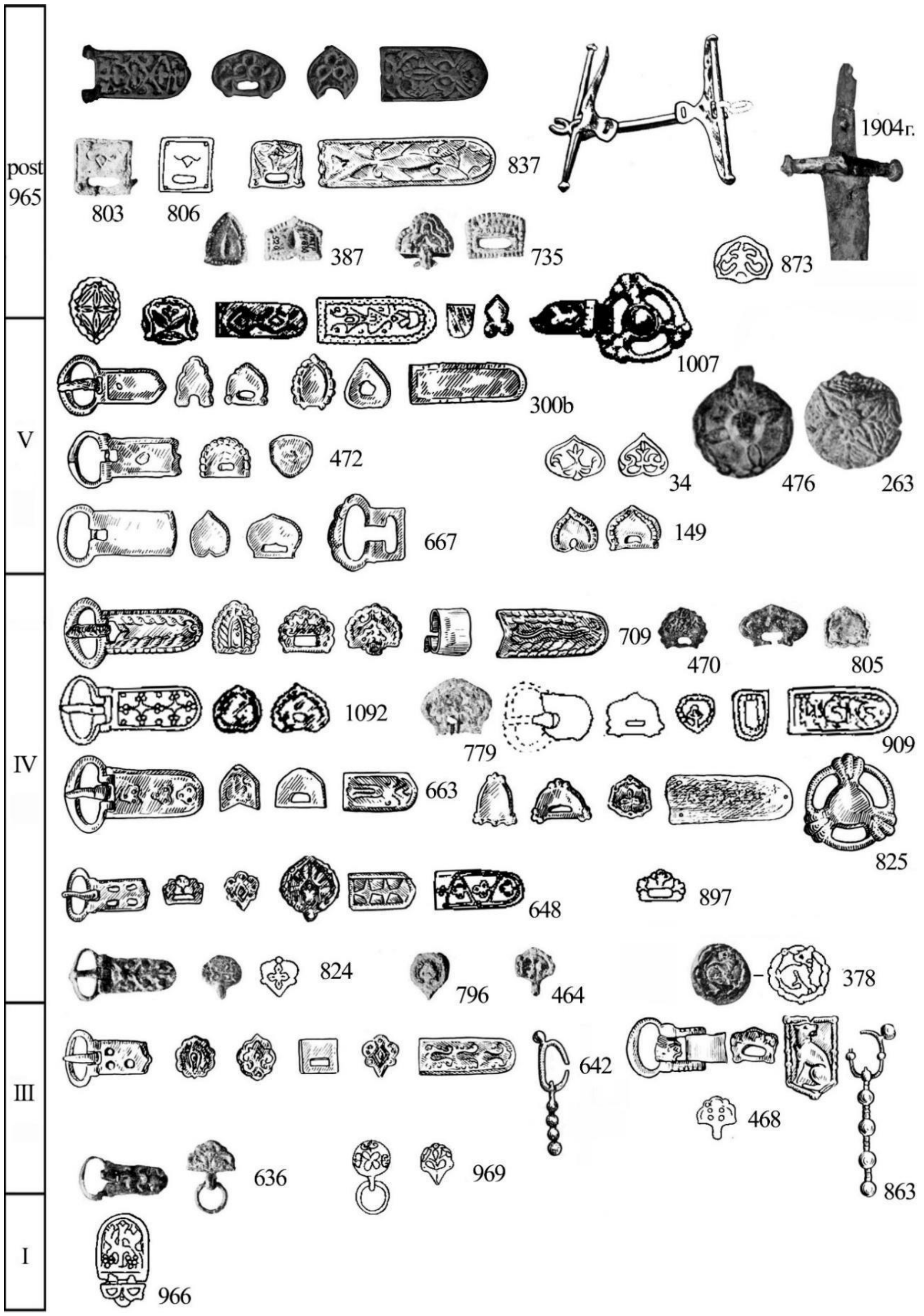


*Fig. S8. Typochronological system of Tankeevka cemetery by Oleksii Komar (after* KOMAR *2018, Fig. 51)*

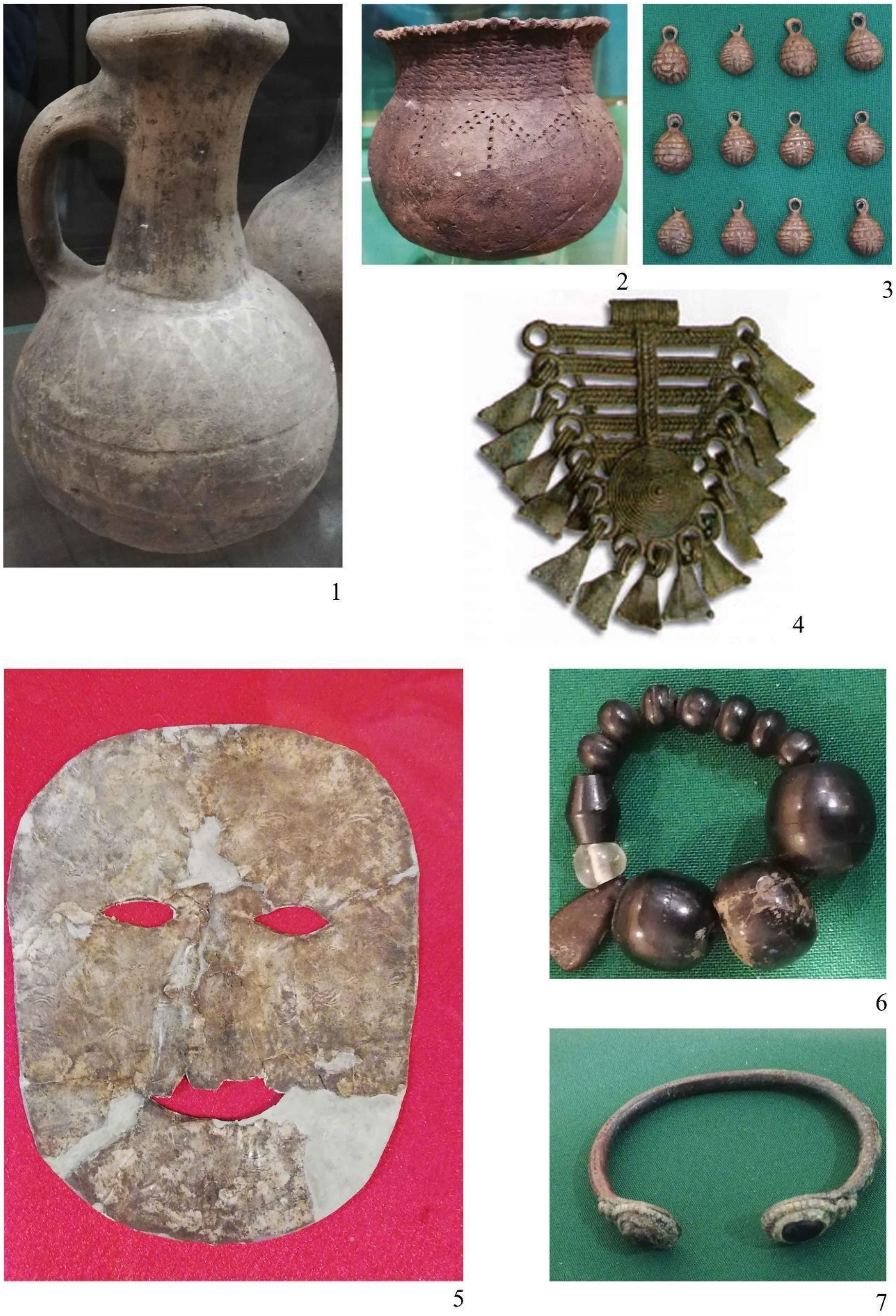


*Fig. S9. Some characteristic and typical finds from the Tankeevka cemetery (photos by Attila Türk in the National Museum of the Republic of Tatarstan)*

#### 4. Karanayevo cemetery

Karanayevo cemetery is located at the western foothills of the Urals, in the northern part of Bashkortostan, in Mechetlinsky District, 1 km south of the village of Karanayevo (SUNGATOV 2016). The first investigation of the site was led by Niyaz A. Mazhitov between 1964 and 1966, during which eighteen kurgans were excavated in the large cemetery covering about 10.000 $m^2$. He published them in his monograph in 1981 (MAZHITOV 1981). The excavations of the kurgan tombs of the Karanayevo cemetery continued in 2001. An area of about 1000 $m^2$ was investigated under the supervision of Flarit A. Sungatov. The twelve uncovered tombs yielded extraordinary finds (SUNGATOV 2016).

The site, located in the South Urals, belonged to a population leading a typical nomadic lifestyle between the $10^{th}$-$12^{th}$ centuries. Based on the burial rites, horse equipment, and especially belt and horse harness mounts made of non-ferrous metals, the cemetery is most closely related to the Uyelgi site located in the Trans-Urals. Additionally, the artefacts of the Hungarian Conquest period are also analogous to them in many respects. Pieces of Srostki-type riding gear are common in the cemetery. However, it should be emphasised that pieces of horse equipment discovered in the early medieval cemeteries of the Ural region reflect a strong nomadic influence even earlier, during the second half of the $6^{th}$ century AD in the so-called 'Ural-Turkic' horizon, as well as during the subsequent transitional Bekeshevo phase (Saltovo III period). Later, the sites of Karanayevo and Sineglazovo are the best examples of the influence of Srostki culture in the Southern Ural (KOMAR 2018). Archaeological finds discovered at the Karanayevo burial ground are contemporaneous with the Saltovo IV and V phases (dated after AD 861). Furthermore, in the area of Bashkortostan (particularly on its northern periphery, at the great pass of the Urals), they belong to those few sites (e.g. Isimbay) that are also contemporaneous with the Subbotsy horizon. In other words, they bear Hungarian archaeological relevance.

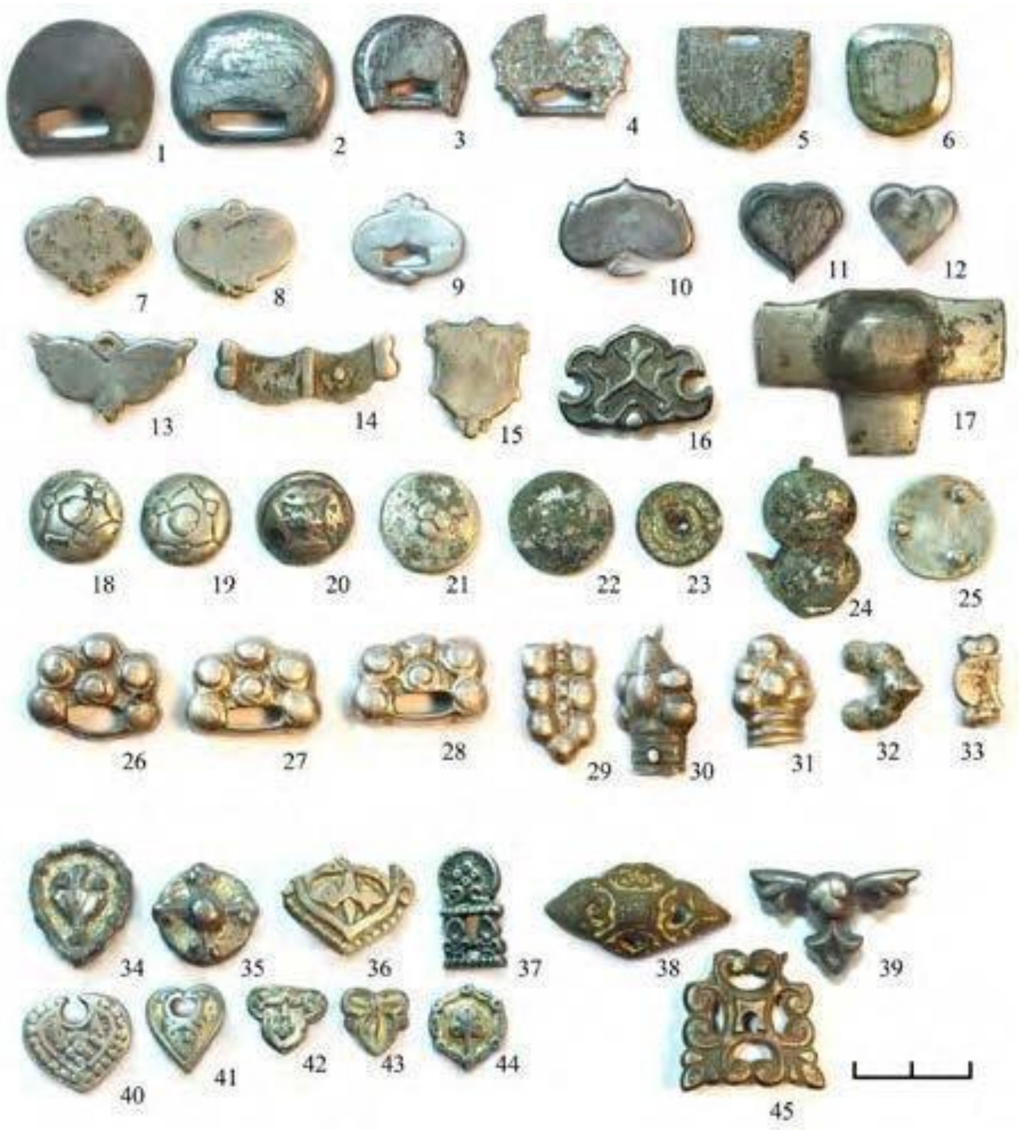


*Fig. S10. Some characteristic and typical belt mounds of the Karanayevo cemetery (photo by Sergei Botalov)*

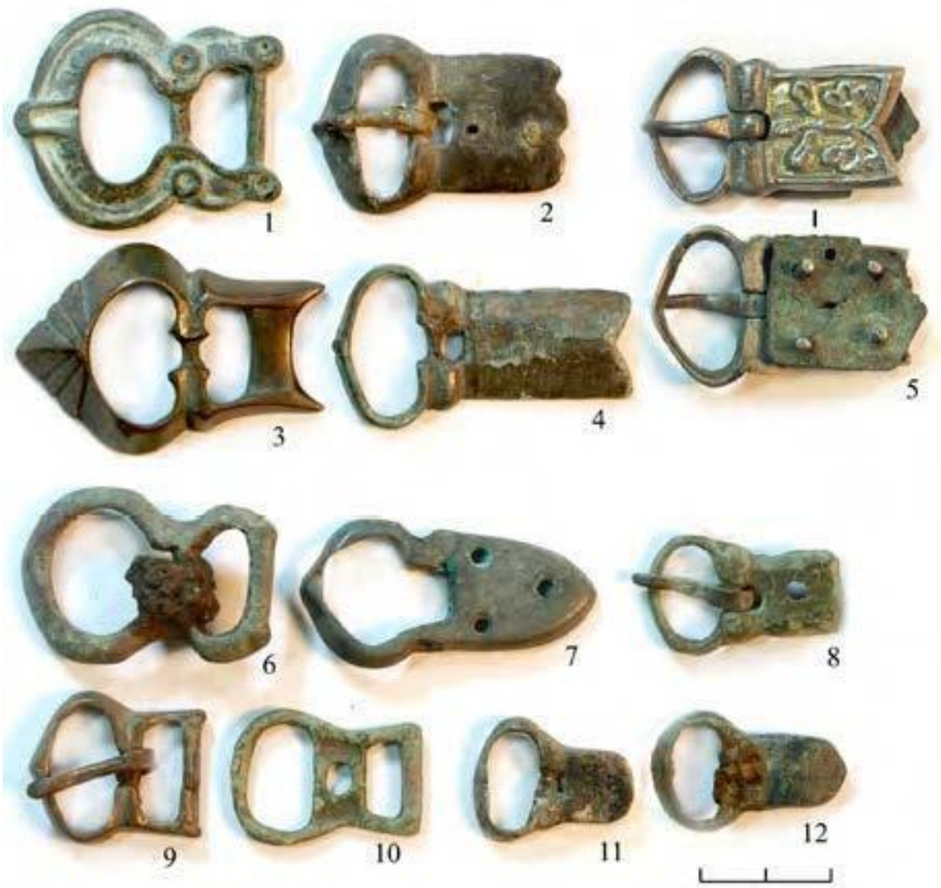


*Fig. S11. Some characteristic and typical belt buckles of the Karanayevo cemetery (photo by Sergei Botalov)*

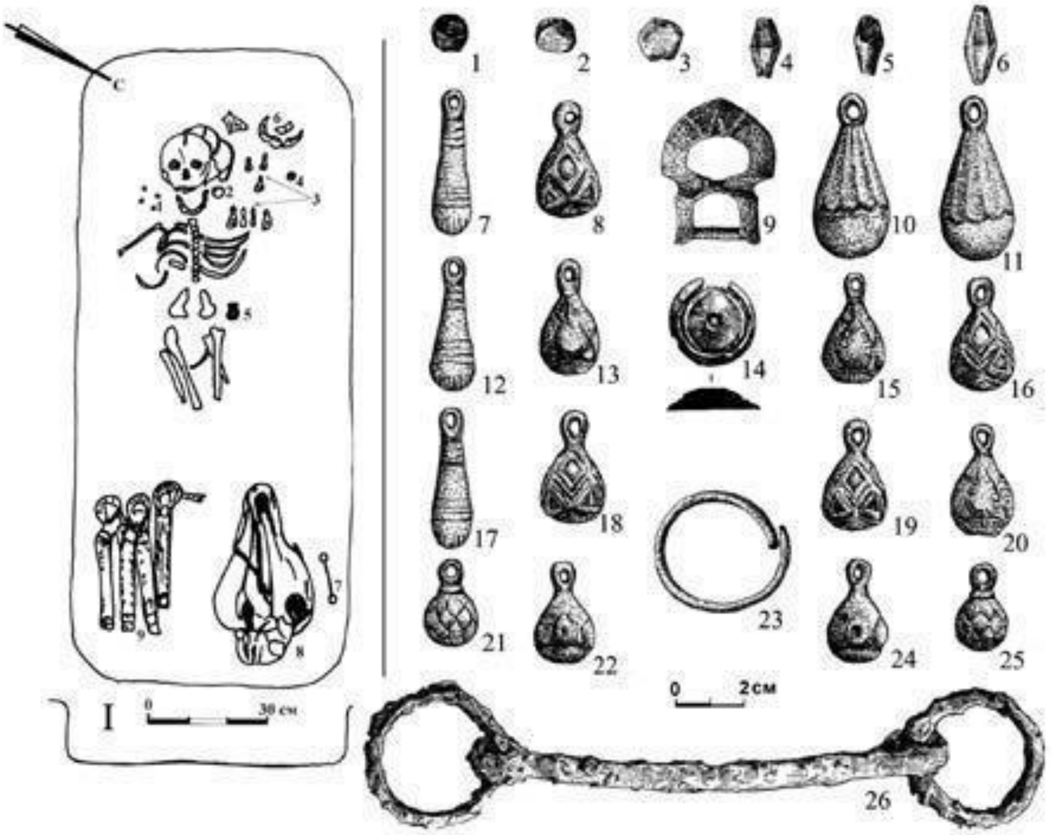

*Fig. S12. Karanayevo cemetery Grave No. 12 (2001) (after SUNGATOV 2016, Fig. 10)*

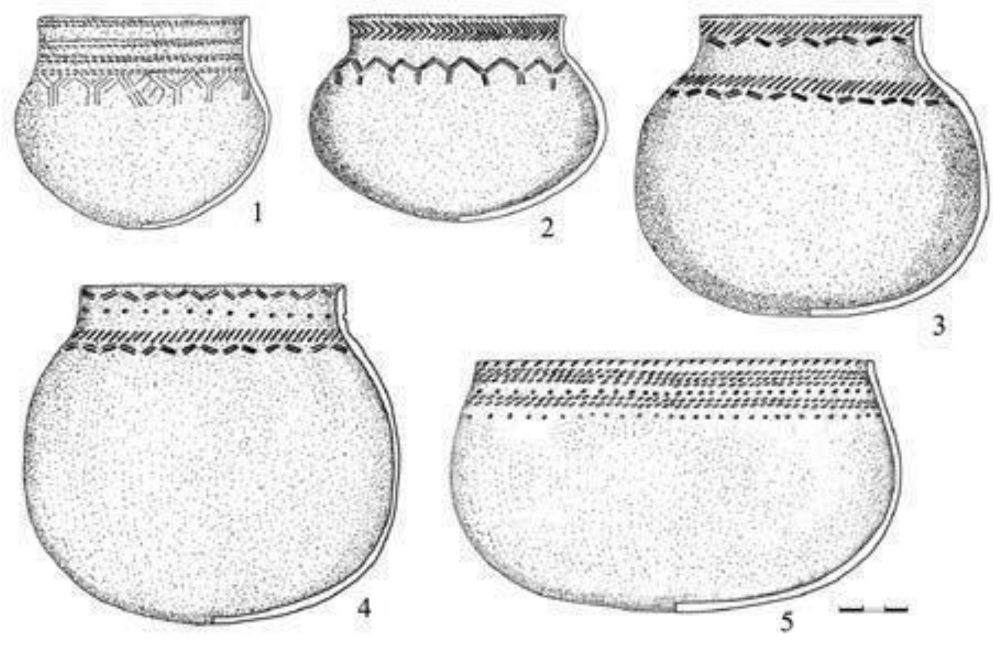

*Fig. S13. Typical ceramic finds from the Karanayevo cemetery (2001) (after SUNGATOV 2016, Fig. 11)*

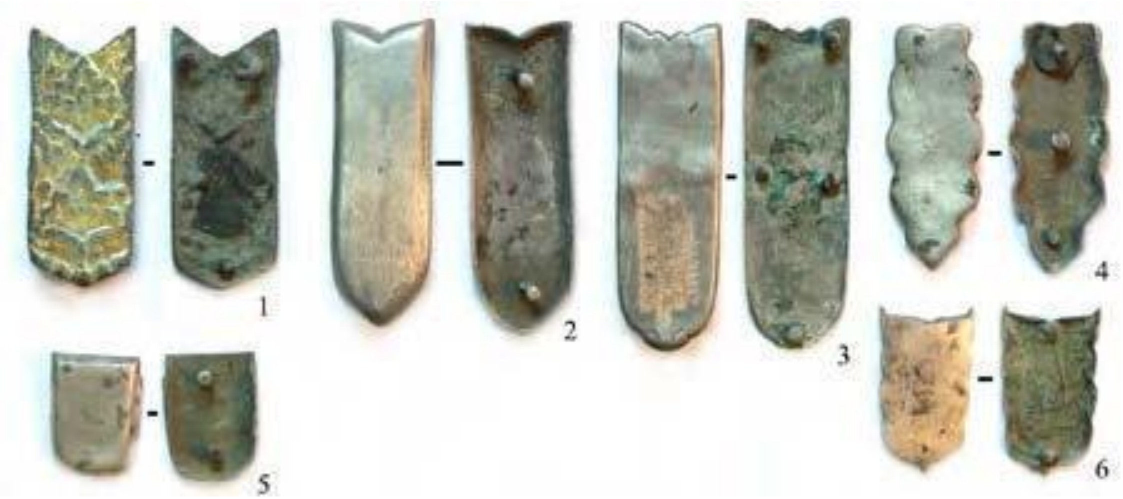


*Fig. S14. Typical belt end finds from the Karanayevo cemetery (2001) (after* SUNGATOV *2016, Fig. 12)*

#### 5. Novinki-type sites

The Middle Volga Region had a great importance in early medieval history, particularly due to the river crossing place located near present-day Samara and the strategic geographical position of the Samara Bend of the River Volga, on the opposite bank of the river. The archaeological finds discovered in the region offered an excellent opportunity for comparing and studying the early medieval history of the eastern and western sides of the Volga. According to historical sources, the relations of the early Hungarians with the peoples of the Khazar Khaganate must have started there; consequently, it was important to take many samples and subject them to archaeological and archaeogenetic analyses.

In the framework of the project, we mainly examined the Novinki-type of sites in the area of the Samara Bend of the River Volga. They belonged to the former neighbours of the Hungarians, a population of Khazar descent, with strong military character, who presumably settled there to defend the border. The Khazar character of the finds and their association with the Bulgars or Khazars is still a matter of debate in the archaeological literature, therefore it was imperative to carry out genetic analyses.

The Novinki archaeological horizon was identified by G. I. Matveyeva in the early 1980s (MATVEEVA 1997). It was named after the first fully excavated kurgan cemetery located in Novinki. To date, more than thirty Novinki-type sites have been excavated in the Samara Bend of the Volga (VASILEV–MATVEEVA 1986). G. I. Matveyeva, A. V. Bogachov, R. S. Bagautdinov, S. E. Zubov, N. A. Lifanov, and D. A. Stashenkov supervised the excavations of the most famous cemeteries located in the vicinity of the settlements of Novinki, Brusyany, Malaya Ryazan, Rozhdestveno, Osinki, and Vipolzovo. The typical burial form of the Novinki-type of sites is the kurgan cemetery, where the mounds contain massive limestone rocks, and there is a cairn above the tombs (BAGAUTDINOV–BOGACHEV–ZUBOV 1998). There are usually one to twelve burials below the barrows. The graves are usually simple pits, but there are also graves with steps, benches, and sidewall niches. Such types of graves are also known without a kurgan (STASHEKOV 1995).

The dead were buried in an extended supine position, with the head to the east, north, or west. There were several kinds of grave goods: pottery vessels, weapons, jewellery, work tools, and horse equipment. The pottery vessels comprised pitchers, jugs, and bowls. Ceramics were also often discovered outside the tombs in the earth of the kurgan testifying funeral feasts held in commemoration of the dead. Jewellery, clothing items, and tools of toiletry were found in men's, women's, and children's graves alike. In men's graves, these are usually represented by the metal parts of a belt set (buckles, fittings, strap ends), earrings, and signet rings. In women's graves, they comprise all kinds of pearls and other items to be strung, pendants, earrings, decorative pins, needle holders, ear spoons, bracelets, and mirrors. Children's graves had amulets and beads.

Weapons and parts of the armour are known from men's graves. These consist of sabres, broadswords, swords with sabre-handle, arrowheads, spearheads, hatchets, war maces, quiver hooks, and bow bone plates. Men's graves also contained horse equipment: stirrups, bits, bridle fittings, and bone saddle fittings adorning the front of the saddle. Based on the analysis of the funerary rites and grave goods, many researchers are of the opinion that the Novinki-type of sites belonged to a part of the early Bulgar tribes that moved to the Middle Volga Region from the North Caucasus and the Sea of Azov Region in the second half of the 7th century, after Kuvrat's *Magna Bulgaria* had fallen apart. However, the population that arrived in the region of the Samara Bend of the River Volga was neither anthropologically nor ethnically homogeneous (GAZIMZYANOV 1995). The descendants of the population that lived in the Novinki-type of sites in the 8th century may have come under the control of Volga Bulgaria in the 9th century.

##### 5.1. The site at Novinki I

At Novinki I, a cemetery where burials with and without kurgans were discovered, like at many of the Novinki-type sites. It was located in the Samara Bend of the River Volga, 2 km east of the village of Novinki (Volga District). The site was first excavated by V. V. Golmsteyn in 1922. The excavations were continued between 1992 and 1993, and in 1999 by the P. V. Alabin Samara Regional Museum of History and Local Lore under the supervision of D. A. Stashenkov. Altogether, ten kurgans were excavated in the cemetery (STASHENKOV 1995). Their diameter ranged from 10 to 18 metres and their height from 0.3 to 0.7 metres. In 1999, excavations were conducted over the entire area of the site, and the area between the kurgans was also explored, where three flat graves were discovered. In the light of this fact, it was necessary to reconsider our assumption concerning the character of the burials and the historical ideas on the number of people living in the Samara Bend of the Volga in the Khazar period (KOMAR 2001).

##### 5.2. The site at Brusyany

The kurgans located near the village of Brusyany, in the Samara Bend of the Volga, were excavated by A. V. Bogachov in 1982. Between 1988 and 1996, the excavations were continued by A. V. Bogachov, R. S. Bagautdinov, and S. E. Zubov in the cemeteries of Brusyany II, III, and IV and at the solitary kurgan of Brusyany II (BAGAUTDINOV–BOGACHEV–ZUBOV 1998).

The kurgan cemetery of Brusyany II was located 1 km west of the village of Brusyany, on the right high bank of the Volga. It comprised thirty kurgans, most of which were investigated in the years 1988, 1989, 1991, 1994, and 1996. All kurgans were small in size, with a maximum height of 1 metre, and 10 to 15 metres in diameter. The number of graves under the kurgans varied from one to eight and were inhumation burials. The bodies were placed in a simple grave pit, in an extended supine position, with the head to the east, north, or west. The grave goods comprised various pottery vessels, jewellery, tools, and weapons.

The kurgan cemetery of Brusyany III was located 3 km north-northeast of the settlement Brusyany. It comprised six kurgans, two of which were unearthed in 1991. There was no cairn in the soil of the kurgan No. 1, which had a large barrow (30 metres in diameter and 3 metres high). Under the kurgan, there was a rectangular ditch, half of which ran beyond the sides of the kurgan. It only had one large round tomb (5 metres in diameter), which yielded objects (an amphora, a candlestick, a large, gilded, leaf-shaped bridle fitting, and a set of silver bridle fittings) suggesting that the deceased buried there was of high social status. The cemetery dates back to the 8th century.

The kurgan cemetery of Brusyany IV is located 0.25 km north of the settlement Brusyany. The cemetery had three kurgans, two of which were excavated in 1996.

The solitary kurgan of Brusyany II was discovered 1.25 km north of the village of Brusyany. During the excavations conducted in 1996, two burials came to light dating back to the 7th and 8th centuries. The grave goods yielded by the burials discovered in the vicinity of Brusyany comprised various types of ceramics, jewellery, and tools of toiletry made of gold and silver (belt sets, earrings, pendants, mirrors), weapons and work tools (arrowheads, remains of bows, spears, blacksmith tongs), and horse equipment (stirrups, bits, and bone plates of the front part of saddles).

According to the archaeologists' view, the kurgan cemeteries of Brusyany belonged to groups of Bulgars and Alans who settled there in the late seventh and eighth centuries (KOMAR 2001).

##### 5.3. The site at Malaya Ryazan

The cemetery of Malaya Ryazan I comprising burials with and without kurgans is situated in the southern part of the Samara Bend of the River Volga, 1.2 km east of the village of Malaya Ryazan. The site had ten kurgans. In 1990, A. V. Bogachov and S. E. Zubov excavated the site. In 1990, 1995–1996, 2009–2010, 2017, and 2019, further excavations were conducted here under the supervision of A. V. Bogachov, S. E. Zubov, N. A. Lifanov, and O. V. Bukina (BUKINA–LIFANOV–ZUBOV 2018). Forty-three graves were unearthed, which were dated to the 8th century. Above most of the burials at Malaya Ryazan I, limestone rocks could be observed. The bodies were laid in the grave in an extended supine position, with the head to the east or north. The men's graves contained wheel-thrown jugs, bridle elements, and belt sets. The women's tombs yielded hand-made jugs, coloured glass beads, bronze bracelets, earrings, and mirrors (BUKINA–LIFANOV–ZUBOV–BAGAUTDINOV 2018).

##### 5.4. The site at Shilovka

In 1992, two kurgans were unearthed during an expedition led by R. S. Bagautdinov from the University of Samara near the village of Shilovka (Sengileyevsky District, Ulyanovsk Oblast). Three graves were discovered under the two kurgans. Even though they had been disturbed, there were the remains of rich furnishing in the tombs: a Byzantine gold *solidus* and *bracteata*, earrings with amethyst pendants, a signet ring, bronze and gilded silver elements of a belt set, bone plates decorated with battle and mythological scenes, and a wheel-thrown vessel. The burials were made in the late 7th and early 8th centuries. The ethnicity of the people buried in the kurgan cemetery of Shilovka is still uncertain. The theory of Bulgar and Khazar origins seems currently the most likely (KOMAR 2001).

##### 5.5. The site at Lebyazhinka

In 1997, an early medieval tomb (Grave 4) was discovered during the excavation of the Neolithic and Bronze Age settlement conducted by the Samara State Pedagogical University and the Institute of History and Archaeology of the Volga Region on the Lebyazhinka farmstead (Krasnoyarsky District, Samara Oblast). The rectangular grave measured 225×65 cm. Around the head of the deceased, small steps were cut in the wall of the tomb. The deceased was a 30–35-year-old Europid man, lying in an extended supine position, with the head to the southeast. The sacrum of a large animal was placed in the grave right next to the head. The grave also contained the iron hanger of a quiver for arrows, three iron arrowheads corroded together, a thick, hexagonal bronze bracelet, a silver ring with a violet oval glass bezel held in place with four small prongs, an iron horse-bit and an iron hatchet, and bronze earrings. The grave discovered on the Lebyazhinka homestead dates back to the 9th century and is the only Ugric burial in the Samara Region where the weapons comprised a hatchet (STASHENKOV–TURETSKII 1999). The Europid skull from the tomb discovered on the Lebyazhinka homestead is similar to the human remains from the Tankeevka cemetery in many ways (GAZIMZYANONV–KHOKHLOV 1999).

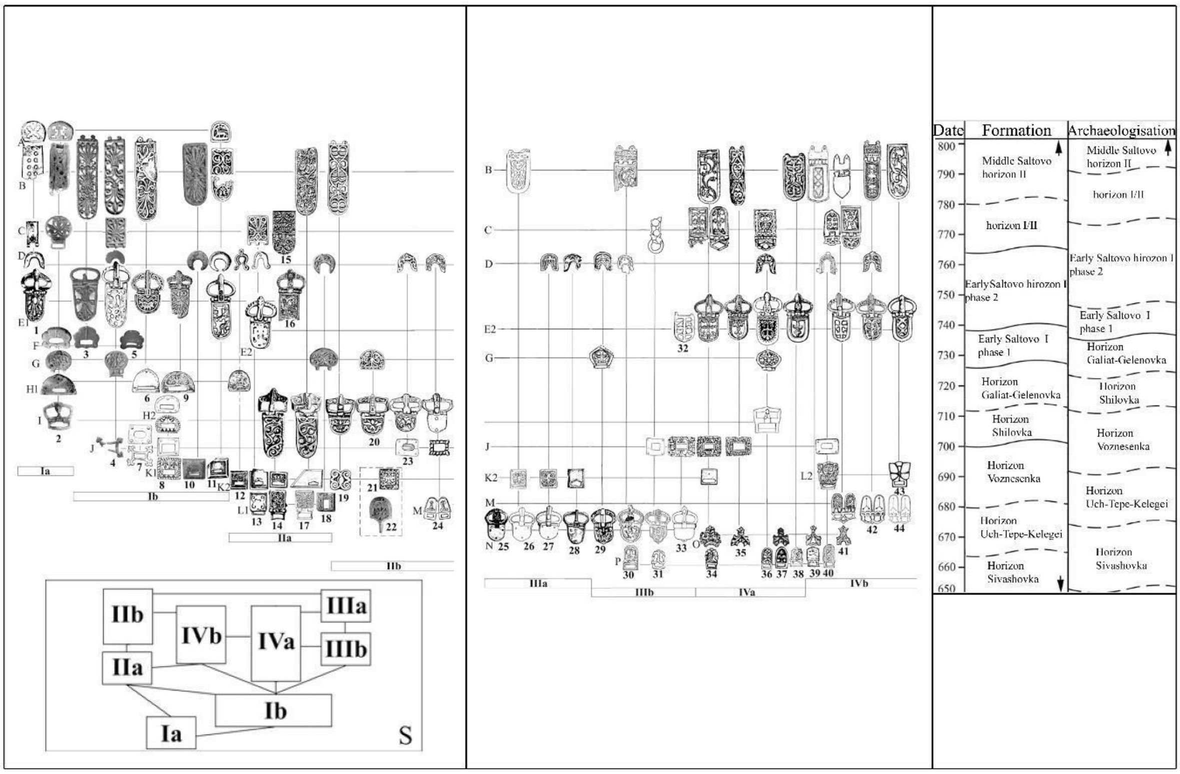


*Fig. S15. Relative typochronological timeline of the Novinki-type sites' belt (1-2) and the absolute timeline (3) following of O. Komar (KOMAR 2001, Fig. 2)*

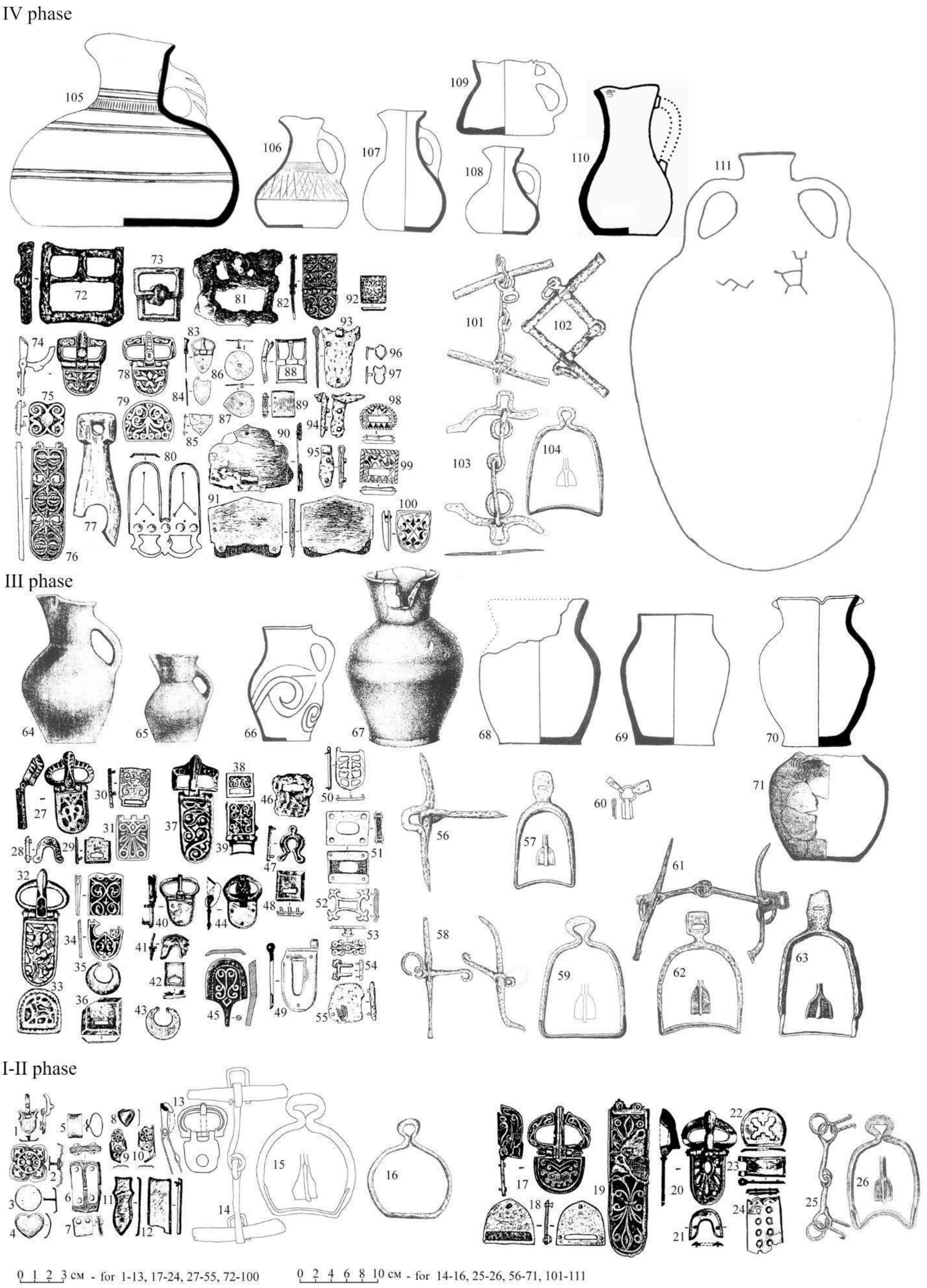


*Fig. S16. Typochronological system of the Novinki-type sites (after Lifanov 2005, Fig. 2)*

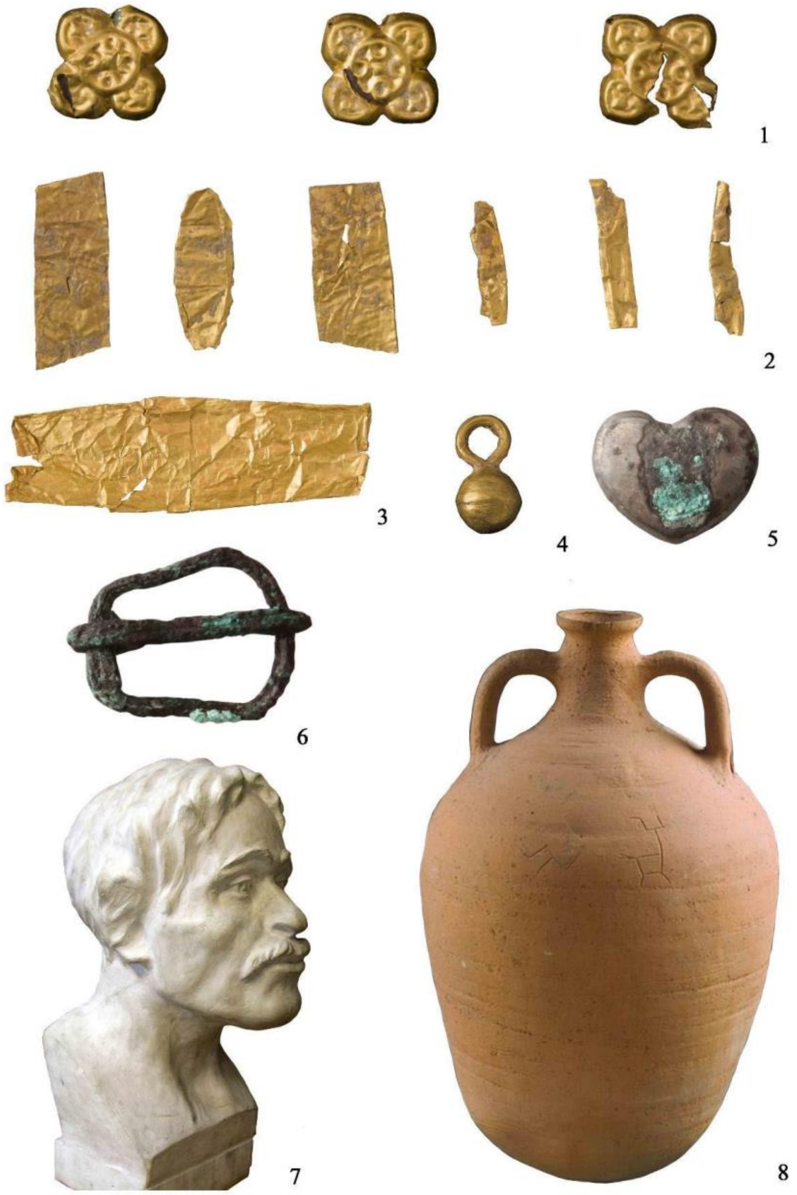


*Fig. S17. Selection from the archaeological finds of the Brusyany IV cemetery (Photo by D.A. Stashenkov)*

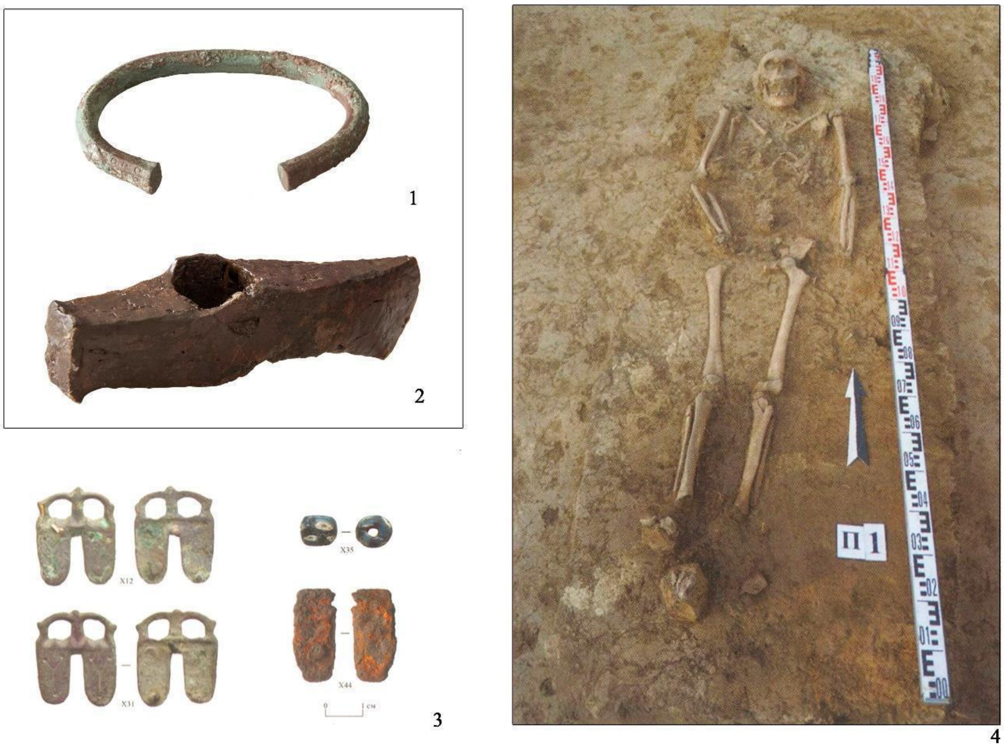

*Fig. S18. Findings from Malaya Ryazan Grave 2019/1. (After LIFANOV 2020)*

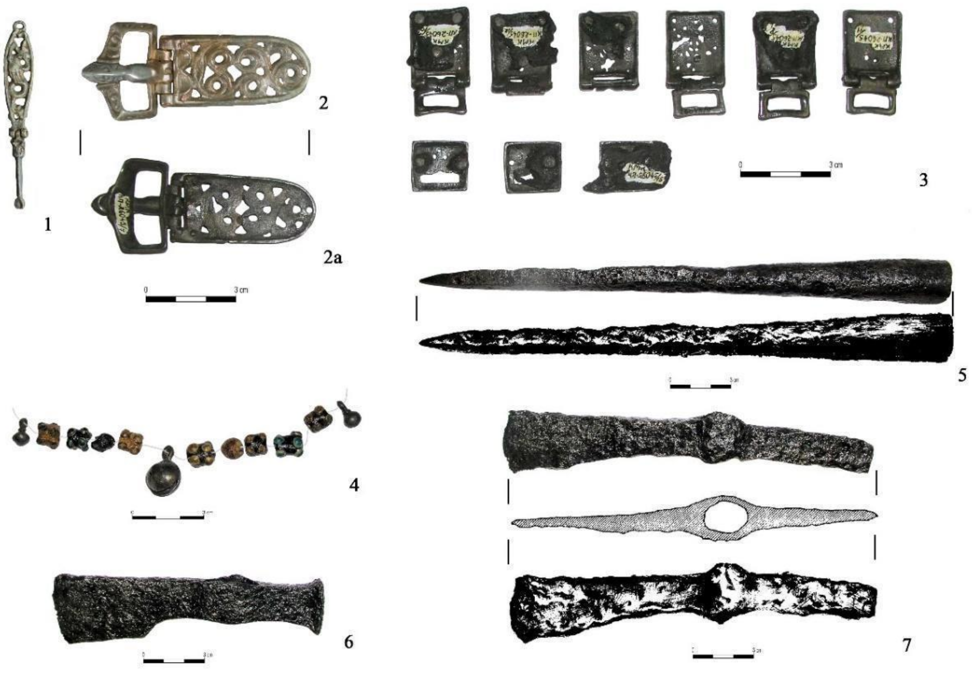

*Fig. S19. Archaeological findings from Lebyazhinka V. Grave 4. (Photos by D. A. Stashenkov)*

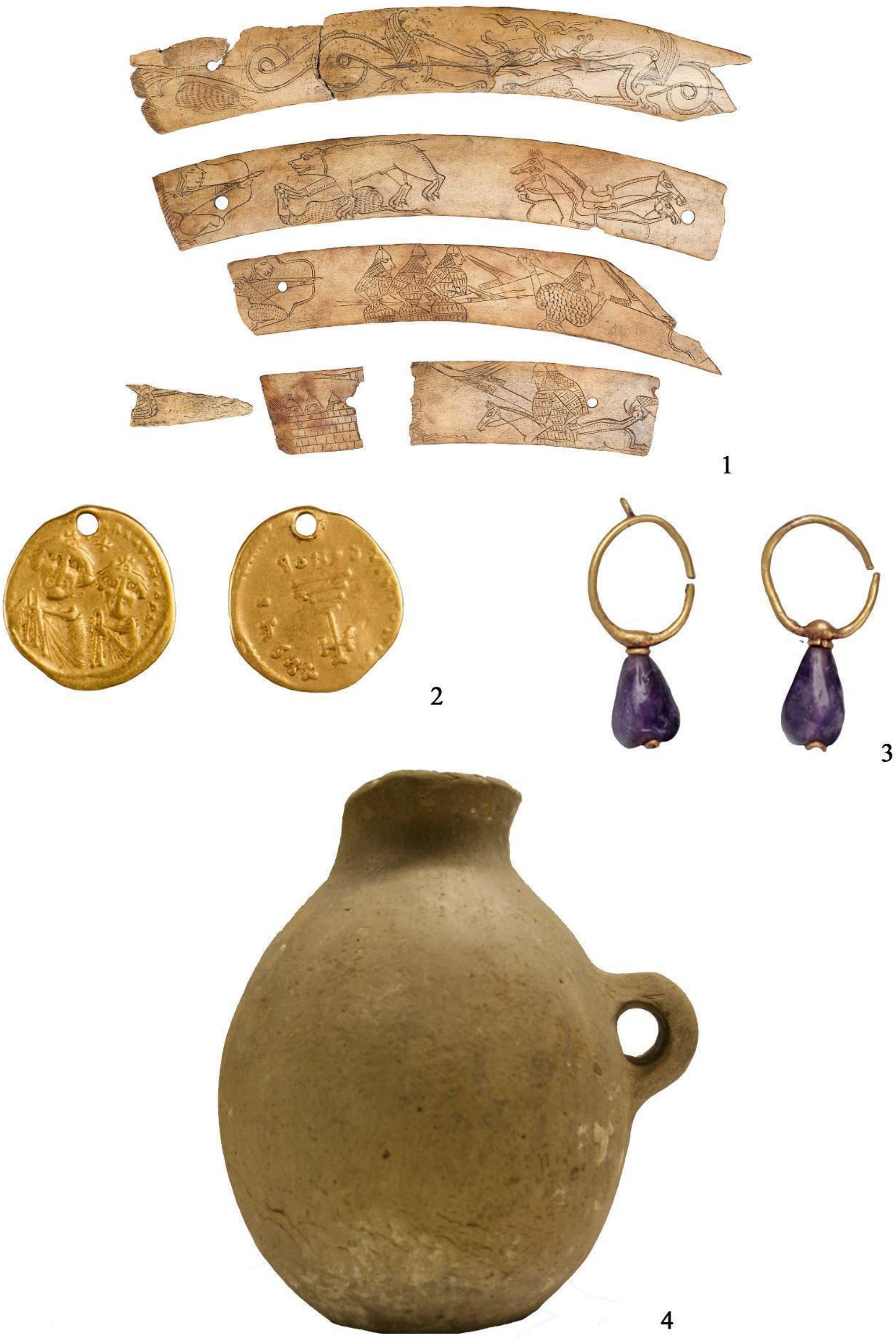


*Fig. S20. Selection from findings of Shilovka kurgan cemetery (Photos by D. A. Stashenkov)*

#### 6. Proto-Ob-Ugric group

The chronology and linguistic identification of the Ob-Ugric peoples and their ancestors have been known for a long time, the Ugric linguistic unit broke up sometime between 1200 and 500 BC. Based on historical and archaeological data, they can hardly be distinguished as a separate people until the 14th-16th centuries AD. Historical and archaeological research dates the separation of the ancestors of Hungarians and Ob-Ugric people to the final phase and disintegration of the Sargatka culture (3rd–5th centuries AD), at the earliest.

In the medieval archaeology of Siberia, the local variants of these people can be identified. The Oronturski-type of sites can be associated with the northern group of Mansi people living along the middle course of the River Ob, while the Potchevash archaeological culture along the lower and middle parts of the River Irtysh can be associated with the southern Mansi. It is important to point out that from the southeast, nomadic peoples with Turkic language and material culture as well as steppe origins arrived in their territory in several stages beginning with the middle of the first millennium BC.

The population of Novochekinsk and Bogochanovka cultures living on the southern edge of the taiga, mixed with the people of the Kulay culture (CHINDINA 2001) living along the Irtysh, to the north, established the following archaeological cultures: the Kashino, Iudino, Petrogrom, and Molchanovo cultures in the Trans-Urals, as well as the Potchevash and Ust'-Ishim cultures along the River Irtysh. In the 8th century, the southern part of the taiga along the River Irtysh was already occupied by a population of southern origin and with Turkic culture and pushed the Proto-Ob-Ugric population northwards. From the fifteenth to the seventeenth century, the Khanty people expanded to the east. It was at that time that the first Ob-Ugric political units were established, such as the Principalities of Konda and Obdorsk. Archaeologically, this period can primarily be characterized by the Saygatinski burial grounds (10th–16th centuries AD). To the south of them, in the forest-steppe region, there was the Mongol Empire followed by the Siberian Khaganate, a Tatar state succeeding the Golden Horde (14th–16th centuries AD), until the Russian conquest of Siberia (1582).

In the area studied by us, stretching from the Trans-Urals to the right bank of the River Irtysh, the Iudino archaeological culture existed between the 10th and 13th centuries AD. The representative pottery of this culture is also known from the Uyelgi site. In the cemeteries without barrows, the former cremation rite was replaced by inhumation, although the role of fire remained prominent in their mortuary cult. Their characteristic finds are mask-like small sculptures representing human faces, masks, fantastic creatures, and animals. Northwest of them, Petrogrom-type of sites are known from the same period. Their trade relations with the Cis-Urals are well known. These people are normally referred to as the ancestors of the Mansi. To the south of them, in the forest-steppe zone between the Urals and the River Ishim, the Bakal archaeological culture could be distinguished based on their horse equipment. It developed as the early Kipchak population coming from present-day Kazakh areas mixed with the local population with an Ugric language and origin who took up a nomadic lifestyle. Its broad chronological framework was earlier set to the 9th-15th centuries AD. Today it is dated between the 4th-13th centuries AD. This culture bears some Sargatka traditions, so based on the cord-decorated pottery emerging there, referred to as Pseudo-Kushnarenkovo-type, S. G. Botalov filled the chronological gap between the Sargatka and Kushnarenkovo cultures (4th–6th centuries AD) in the history of the early Hungarians with this culture. (In summary: BOTALOV 2016; CHINDINA 2001; FJODOROVA ET AL. 1991; KONIKOV 2007; MATVEEVA 2018a,b; ZELENKOV 2018)

We also subjected the Proto-Ob-Ugric population living next to the Hungarians, as well as the Khanty and Mansi people under genetic analysis to find out more about the historical and archaeological connections and development described above. We studied samples from relatively recent and carefully recorded excavations together with our colleagues Tyumen. Therefore, our archaeogenetic studies were based on modern archaeological data from southwestern Siberia and the Trans-Urals instead of the Uralistic linguistic model. In the framework of this work, we focused on the chronology of the archaeological sites presented in the followings.

##### 6.1. Nizhneobskaya culture

The Nizhneobskaya culture occupied a vast area of the West Siberian taiga. Its central area was found along the River Ob and its tributaries (Tobol, Ishim, and Irtysh). In the north, it was bordered by the Kara Sea. In the south, it stretched to the geographical latitude of the cities of Tyumen, Tobolsk, and Omsk. In the middle of the 20th century, V. N. Chernetsov identified this culture during his investigations of the Us-Told fortified settlement in the field. The Nizhneobskaya culture spans a long period between the 4th and 15th centuries AD. Ust'-Tara VII burial ground (along the River Irtysh near Omsk) discussed in this article belongs to the early Karim phase (4th–6th centuries AD) of this culture. The Ust'-Tara VII, Kozlov Mis, Krasnoyarsk 4, Alexeyevka 50, and Alexeyevkva 51 cemeteries are large Western Siberian sites belonging to the Migration Period. The finds of the Barsov I cemetery can be dated to the Kuchiminskaya phase of the Nizhneobskaya culture (7th–9th centuries) and are often associated with the Potchevash culture found in the southern half of the taiga along the Irtysh. Living near the people of the Bakal culture on the forest-steppe, the Karim and Kuchiminskaya population maintained trade relations with the steppe nomads, which explains why so many Eastern European imported goods (belt sets, bronze cups, etc.) were discovered at the site. It can be assumed that in terms of ethnicity, the population of the Nizhneobskaya culture was of Ugric origin (southern Khanty) (KONIKOV 2007, ZELENKOV 2018).

###### 6.1.1. The site of Ust'-Tara VII

In the area of the Rivers Ob and Irtysh, the earliest sample comes from the site Ust'-Tara VII belonging to the Karim culture – or rather Karim chronological phase – dated to the late 4th and 5th centuries AD (BORZUNOV–CHEMYAKIN 2014). The site is located in the southern part of the taiga, along the River Irtysh, in the Tara District of the Omsk Oblast. It consisted of eight kurgans, which were up to 0.5-metre high. The sites were explored by I. E. Skandakov between 1990 and 1994. The archaeological heritage can be characterized by deformed skulls and typical Early Migration Period and Hun artefacts (SKANDAKOV–DANCHENKO 1999).

Grave 1 in Kurgan 9 at Ust'-Tara VII was oval, oriented north-south. The grave pit measured 3×1.9 metres. The south end of the tomb was 0.72-metre deep and its bottom was straight. The northern part of the tomb was raised by 0.25 metres like a step. In the northern part of the grave, burnt pieces of wood were discovered during the excavation, the largest pieces of which were 24–36 cm long, 8–12 cm wide, and 2–3 cm thick.

The deceased was an approximately 20-year-old young woman. She was lying in the grave in an extended supine position, with the head to the south. Due to the shape of the bottom of the grave, the lower parts of the legs and the heels were raised at the edge of the grave. At the end of the left tibia, there was a ceramic vessel. Around the feet, the remains of charred, vertical stakes of 4 cm in diameter were also discovered. On the left side of the hip, there was a bronze buckle accompanied by the pieces of a leather strap, which were covered by the remains of charred birch bark. From the other grave goods, special mention should be made of a bronze necklace consisting of four plates and fastened to each other with iron rivets. On the left side of the skull bearing the marks of deformation was a temple pendant made of wire, and another one was under the skull. One end of the necklace, small pieces of charcoal, and fragments of charred birch bark were also discovered beneath the skull.

##### 6.1.2. The cemetery of Barsov Gorodok I

The burial site of Barsov Gorodok I is found in the northern part of Tyumen County, in Surgut District (a few kilometres from Surgut). It lies in the valley of the River Ob, on an elevated strip of land called Barsov, bordered by two depressions and the tributary of Utoplaya. The exploration of the site covering a huge area (approx. 1,600 m$^2$) and numerous historical periods, started in the late 19$^{th}$ century (F. Martin). The site also served as a place of worship for the Khanty until modern times. From the beginning of the 1990s, N. V. Fyodorova, A. P. Zikov, K. G. Karacharov, and Yu. P. Chemiakin investigated the site. Over 240 graves have been brought to light. At the site, the medieval graves belonged to cemetery Barsov I, while the Early Iron Age graves were found in burial ground Barsov III. The earliest tombs in Barsov I cannot be dated earlier than the very end of the 7$^{th}$ century and the 8$^{th}$ century, whereas the latest graves were probably dug in the 13$^{th}$ century. The tombs of the burial ground form groups in terms of both space and time.

Part No. I of the cemetery, from which our samples also come, was excavated by K. G. Karacharov from 1988 to 1989 and belongs to an early period (7$^{th}$–9$^{th}$ centuries AD) of the cemetery (CHEMYAKIN–ZIKOV 2004). The upper chronological boundary of the cemetery dates to the early 13$^{th}$ century and the Kintunovo period (late 9$^{th}$–early 13$^{th}$ centuries AD) of medieval southwestern Siberia, the archaeological finds of which appear from the middle section of the River Ob to the Ural Mountains and form part of the Great Kulaysk cultural sphere. This was not a peaceful period in the life of the area as numerous fortified settlements were established. In addition to the traditional fishing, hunting, and gathering lifestyle, cattle farming must have existed there by then, and this is when horse farming also emerged sporadically. Iron and bronze processing saw a huge development. Based on the artefacts discovered at the cult sites, high-quality castings were produced. A complex melting furnace was also brought to light. In addition to the use of moose, long-distance export became dominant in the field of the fur trade, as indicated by the specialized hunting techniques and rudimentary industrial processing carried out locally. From the Cis-Urals, objects imported from the Kama Region, as well as Russia and Western Europe, also appear among the local artefacts. The northern part of the population of the Kintunovo period was presumably composed of Samoyed-speaking peoples, the ancestors of today’s Nenets.

The graves reviewed in this article were excavated in 1989 as part of an archaeological expedition carried out by the Ural State University (KARACHAROV 2004). These graves belong to the group dated to the 8th and 9th centuries (KARACHAROV 1993). Both tombs were disturbed in the past, but the remaining finds correspond to this chronology. The 8th- and 9th-century sites of the northern wooded area of Western Siberia form part of the Kuchimskskaya culture. This culture is characterized by a high degree of unification in the construction of houses and forts, as well as pottery making. The large number of imported goods discovered, even from remote areas, show that trade was highly developed. In this regard, the widespread use of Iranian bronze vessels in the region is particularly noteworthy (Chemyakin–KARACHAROV 2002). The groups of graves show differences in terms of anthropology (POSHEHONOVA 2010), which testifies to intensified migration processes and a considerable mixing of populations (TEREHOVA–KARACHAROV 1994). The historical-cultural medium that left Kuchimskskaya culture behind abruptly ceased to exist in the last third of the 9th century. The large fortifications were abandoned, the pottery making changed, and the Western objects clearly began to dominate among the imported goods. It is likely that these events were related to the large-scale political and economic changes taking place in the Urals and the Volga Region at that time, especially the establishment of the state of Volga Bulgaria. Therefore, it can be assumed that behind the cultural change, there was an expansion from the region of the Urals.

##### 6.2. The Potchevash culture

The Potashevas culture was found along the Lower and Middle sections of Irtysh and Ishim stretching from the forest-steppe region to the taiga zone in the regions of Tyumen, Omsk, and Novosibirsk. This culture was identified by V. I. Moshinskaya in the mid-twentieth century after the sites of the Chuvasky Mis near the city of Tobolsk. The sites of the Potchevash culture can be dated 6-9th centuries. The culture of the population was heterogeneous, comprising Nizhneob, forest-steppe Bakal, and early Turkic traditions. In terms of ethnicity, it was identified to be of Ugric-Samoyed origin (GENING–ZDANOVICH 1987).

This culture developed due to the expansion of the population of Kulay culture to the forest-steppe, where they assimilated local people. Their finds include jewellery decorated in a special 'animal style', as well as cultic ornaments bearing human figures. The population engaged in complex farming has historically been associated with the ancestors of the Ugric-speaking peoples primarily, mainly with the ancestors of the southern Khanty, but the possibility of their identification with the Samoyeds also emerged earlier. Nevertheless, it is clear that nomadic peoples (Srostki culture) of steppe origin, presumably speaking Turkic languages, arrived there through the valley of the River Irtysh in the eighth century and merged into the population of the culture. Their presence is attested by weapons, belt fittings, and horse equipment. Despite the influence of peoples coming from the south, the culture developed further into the Ust'-Ishim culture (9th–13th centuries), while the southeastern half of the culture became a steppe culture and formed the basic Baraba population (ZELENKOV 2018).

###### 6.2.1. Vikulovo cemetery

The Vikulovo (Vikulovskoye Kladbishche) cemetery is located in the administrative area of Tyumen County. It is a burial ground consisting of shaft graves without barrows. It lies on the western part of a west-east ridge, on the terrace of the River Ishim. In 2008, V. V. Ilyushina carried out an excavation at the site, a significant part of which belongs to the Potchevash archaeological culture (6th–9th centuries AD), and within that, the period between the 7th and 9th centuries (ILYUSHINA 2009).

Grave 1. It was located in the eastern part of the sector. The grave pit was approximately rectangular, with rounded corners, and was discovered at a depth of 0.6 metres from today's surface. The skull and collarbone were found in its northern part, while the tibia was in the south. The grave measured 1.75×2 metres, and its depth was 0.04-0.08 metres from the subsoil. It was filled with dark grey and yellow clayey soil. It was oriented north-south. At the top of the backfill of the grave, there were three minor pits, presumably the remains of a structure erected by the Russian population. The remains of approximately four skeletons were discovered in the tomb. Despite the clayey soil, the bones were in poor condition, which was probably caused by the fact that the grave had been disturbed by the structure made by the Russian population.

Skeleton No. 1. It was discovered at the western wall of the grave. The femur, tibia, and two fibulae could be found, as well as the pars petrosa of the skull. Due to the poor condition, the sex and age of the deceased could not be determined. Based on the position of the bones, the body must have been lying with the head to the north.

##### 6.3. Ust'-Ishim culture

In the mid-twentieth century, V. N. Chernetsov, presenting the sites located at the lower part of the River Ob and in the southern zone of the taiga, along the Irtysh, gave a general description of the Kintusovo phase of the region (Nizhneobskaya culture). Later, in connection with the field survey made near the Omsk section of the River Irtysh, V. A. Mogilnikov and B. A. Konikov demonstrated the characteristics of the culture of the region in the 9$^{th}$-13$^{th}$ centuries. Its most important features were the kurgan burials and the great role of animal husbandry, while the lower parts of the River Ob were characterized by remains of fishing and burials without kurgans. Based on these factors, B. A. Konikov considered the Ust'-Ishim culture found in the southern part of the taiga along the Irtysh to be of special interest (KONIKOV 2007). The sites belonging to the Ust'-Ishim culture were predominantly located in the taiga zone of Western Siberia, in an area bordered by the River Vasyugan in the north, the River Tobol in the west, and the River Tara in the east (the latter being a tributary of the Irtysh). The culture can be dated between the end of the ninth century and the thirteenth century on the basis of goods imported from Bulgaria, Russia, and the territory along the Kama. The sites of the Ust'-Ishim culture are assumed to have belonged to the ancestors of the southern Khanty, who lived in close contact with Turkic-speaking peoples (the Srostki culture).

###### 6.3.1. The cemetery of Ivanov Mis I

The cemetery of Ivanov Mis I with kurgans was found in the vicinity of the settlement of Ivanov Mis, in the Tevriz District of the Omsk Oblast, 300 km north of Omsk, on the elevated bank of the River Irtysh. The burial ground dated to the 13$^{th}$ and 14$^{th}$ centuries comprised over fifty kurgans, with a diameter of up to 14 metres and a height of up to 1 metre. In 1991, the Omsk expedition led by B. A. Konikov excavated three kurgans (KONIKOV 2007). The samples of this study come from these kurgans.

**Grave 3 in Kurgan 10** was discovered in the northeastern part of the kurgan. The grave was rectangular, oriented northwest-southeast. The enclosing dimensions of the grave were 2.15×1.25 metres, its walls were vertical and its corners were rounded. The grave contained two bodies lying in an extended supine position. The heads of both were in the northwest. Two ceramic vessels had been placed in the southeastern corner of the grave with their bottoms facing up, one of which fell on the cover of the grave and the other on the bottom of the grave pit. Remains of wood and birch bark were preserved on the skeletons. On the man's skull, there was half of a bronze rattle, and there was a lazurite pendant on each side. On the inner side of the left radius bone lay a single-edged iron knife. Between the two skulls, a bronze rattle was discovered.

**Grave 6 in Kurgan 10** was found in the western part of the kurgan. The grave had been disturbed. The parietal bone of the skull was turned north-northwest, with the cheekbones facing up. To the west of the skull, there was a pottery vessel lying on its side, with its mouth turning to the northwest. Below the vessel, there was a bone arrowhead with pointed tang.

**Grave 1 in Kurgan 12** was found in the middle of the kurgan. It was a robbed, rectangular grave directed northwest-southeast. It had vertical walls, rounded corners, and a straight bottom. The skeleton was also affected by the disturbance. The body was lying in the grave in an extended supine position, with the head to the northwest. Between the skull and the corner of the tomb, there was an iron arrowhead with pointed tang hammered flat. Between the bones of the lower legs, there was an oval wooden object, 7 cm in diameter. Parallel to the tibia lay a fragmentary, decorated plate made of bone. There were wooden remains in the southern part of the grave. In addition to these, a single-edged knife with a wooden sheath, a bronze lunula, and decorated bone plates were placed in the tomb.

##### 6.3.2. The kurgan cemetery of Panovo I

The kurgan cemetery of Panovo I was located 4.5 km southwest of the village of Panovo (Ust'-Ishim district, Omsk region) on the first terrace above the floodplain, on the right bank of the Irtysh. It consisted of seventy-seven kurgans with a diameter of up to 20 metres and a height of up to 1.2 metres.

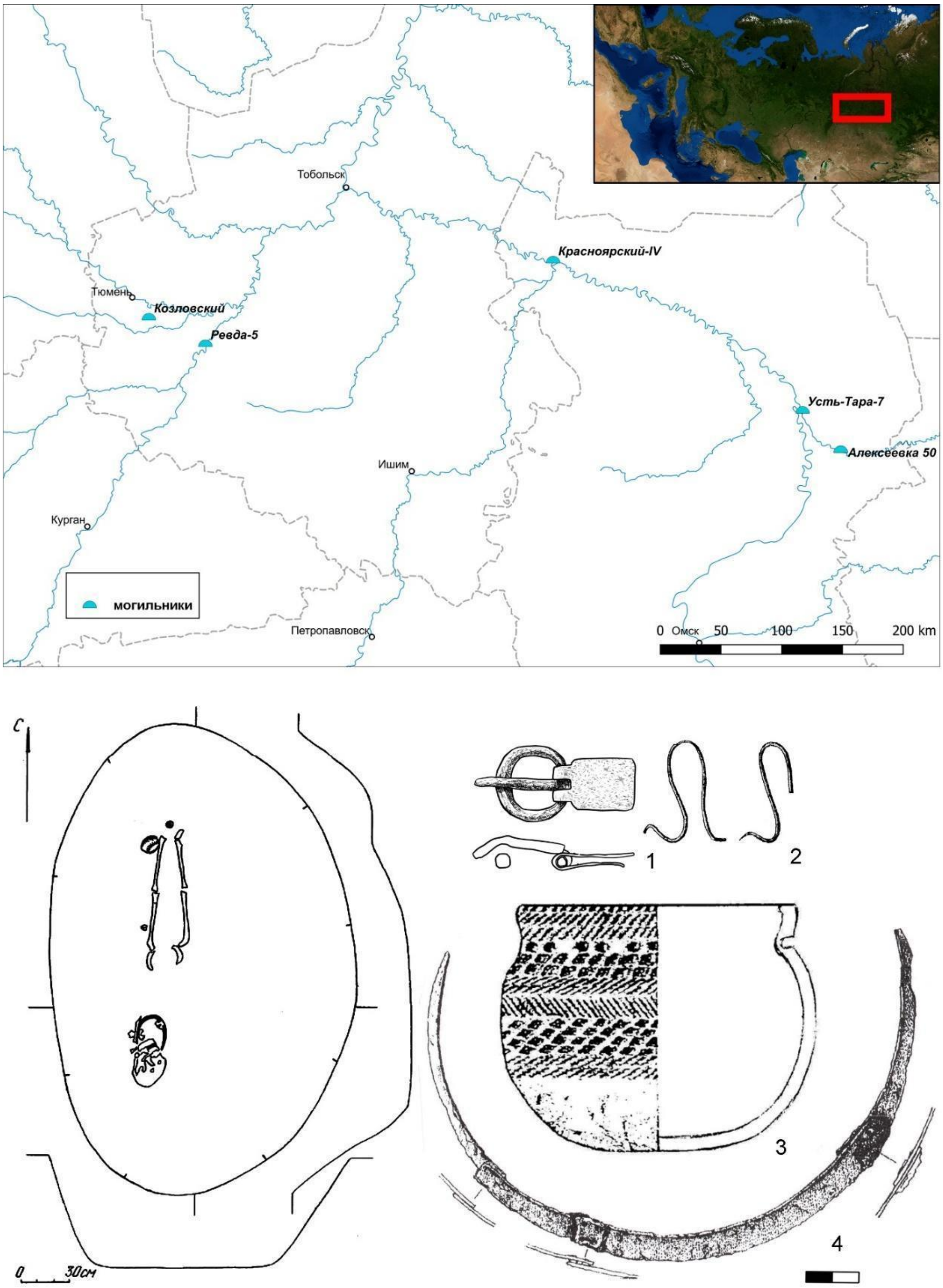


*Fig. S21: 1. The Karim phase of the Nizhneobskaya culture along the Tobol and Irtysh, in the southern taiga zone (Map by A. S. Zelenkov); 1–4: Ust'-Tara, Grave 9 in Kurgan 1 (SKANDAKOV–DANCHENKO 1999)*

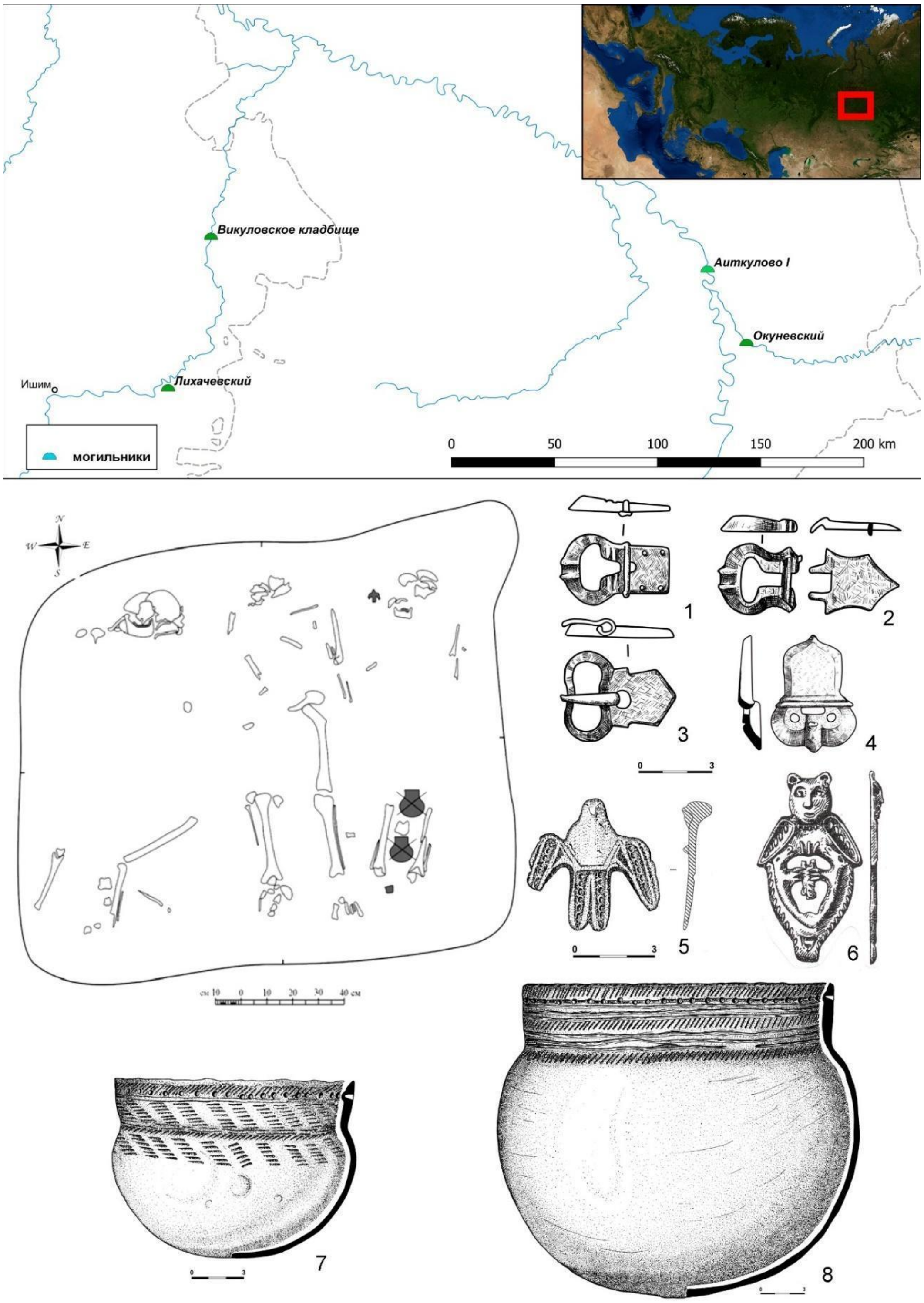


*Fig. S22: The distribution of cemeteries belonging to the Potchevash culture (Map by A. S. Zelenkov). 5,7-8: Vikulskoe Kladbische Grave 1. (after KONIKOV 2019). The representative finds of the Potchevash culture (1–4, 6 from the Okunevsky cemetery) (after MOGILNIKOV 1987)*

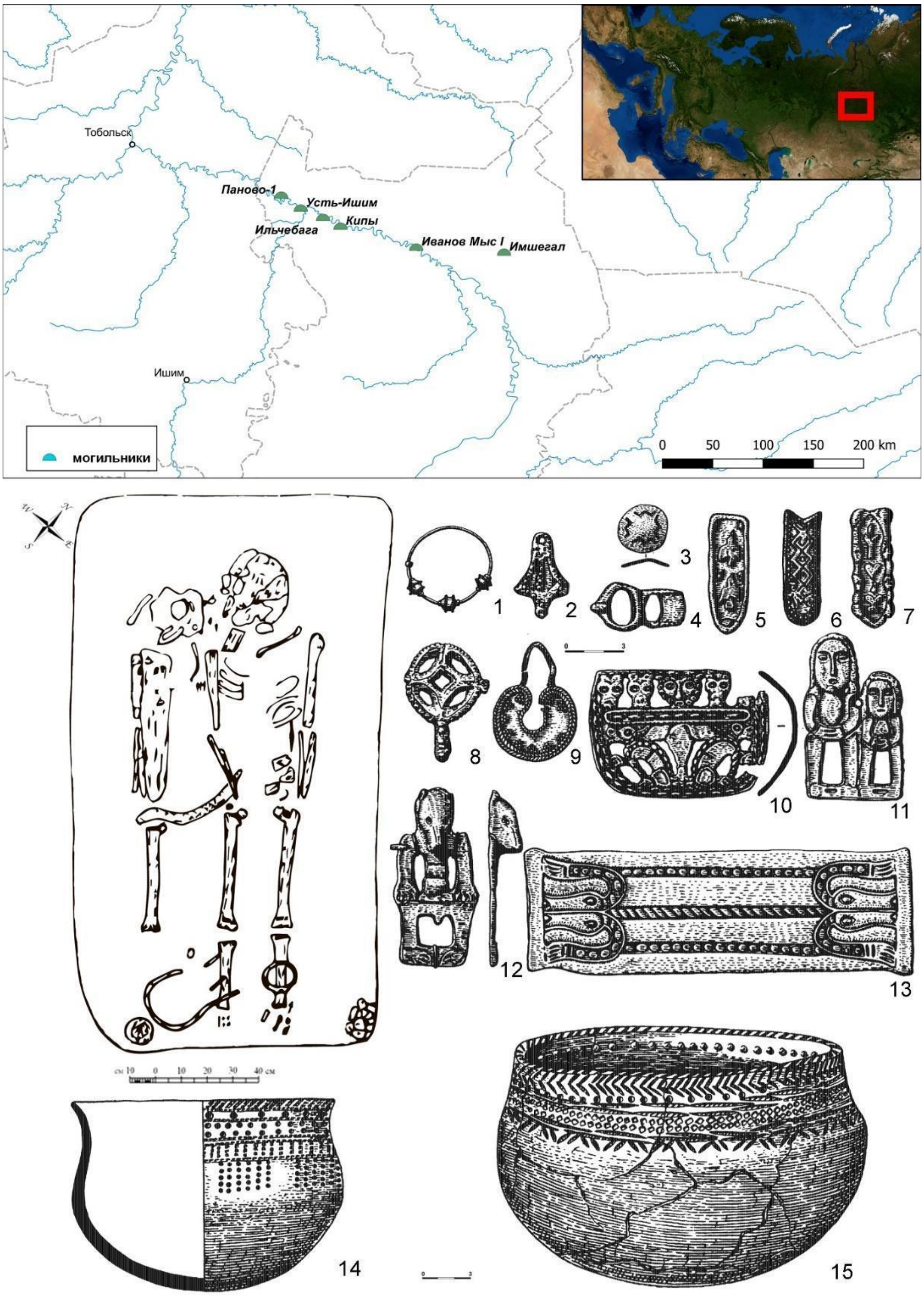


*Fig. S23: The distribution of sites belonging to the Ust'-Ishim culture (Map by A. S. Zelenkov). 1–15: Representative finds from cemetery Ust'-Ishim Cemetery (Ivanov Mis I Kurgan 3 Grave 10) (after MOGILNIKOV 1987)*

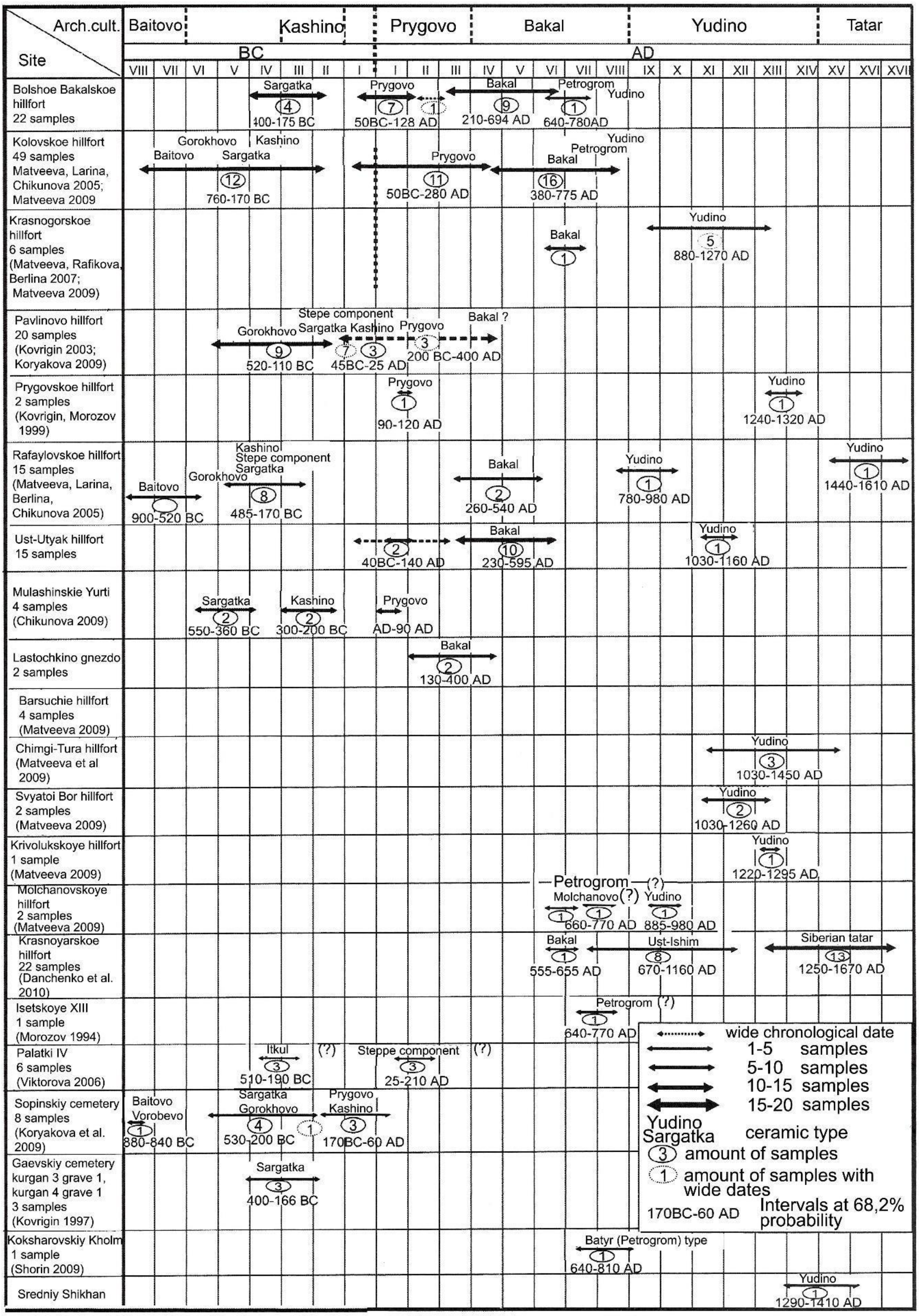


*Fig. S24: The chronology of medieval cultures and sites in Western Siberia and the Trans-Urals based on $^{14}C$ dating – (after* BOTALOV *2016, Table 1.)*

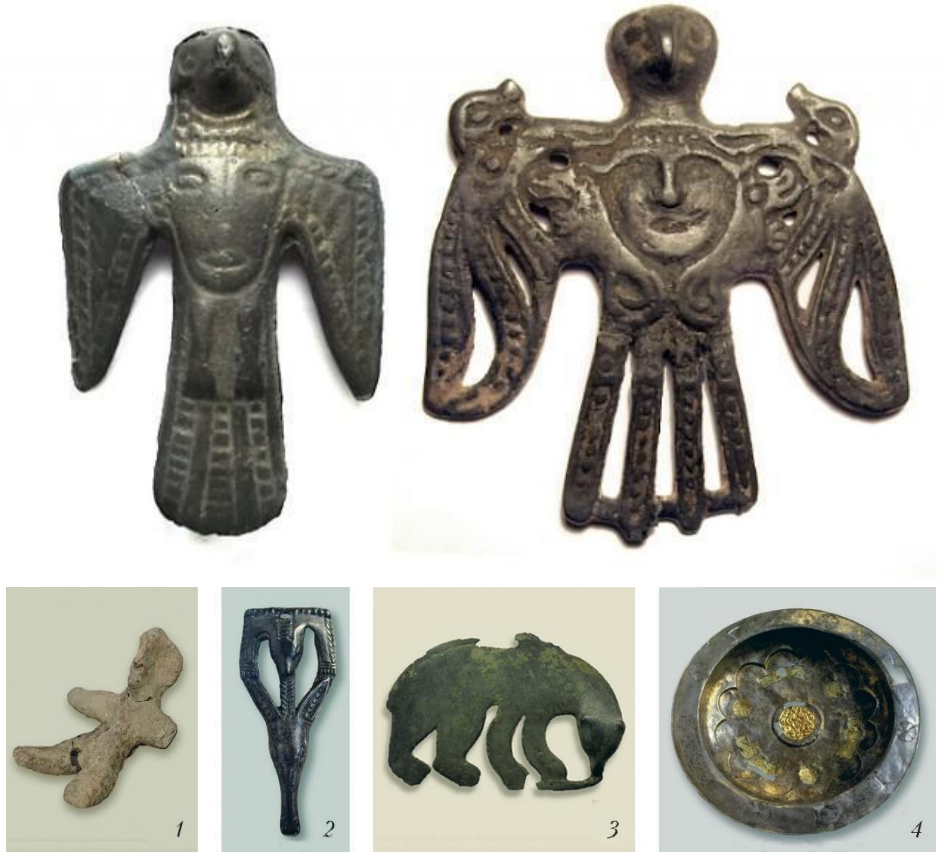


*Fig. S25-26: Representative find from cemetery of Barsov Gorodok (photos by K. G. Karacharov)*

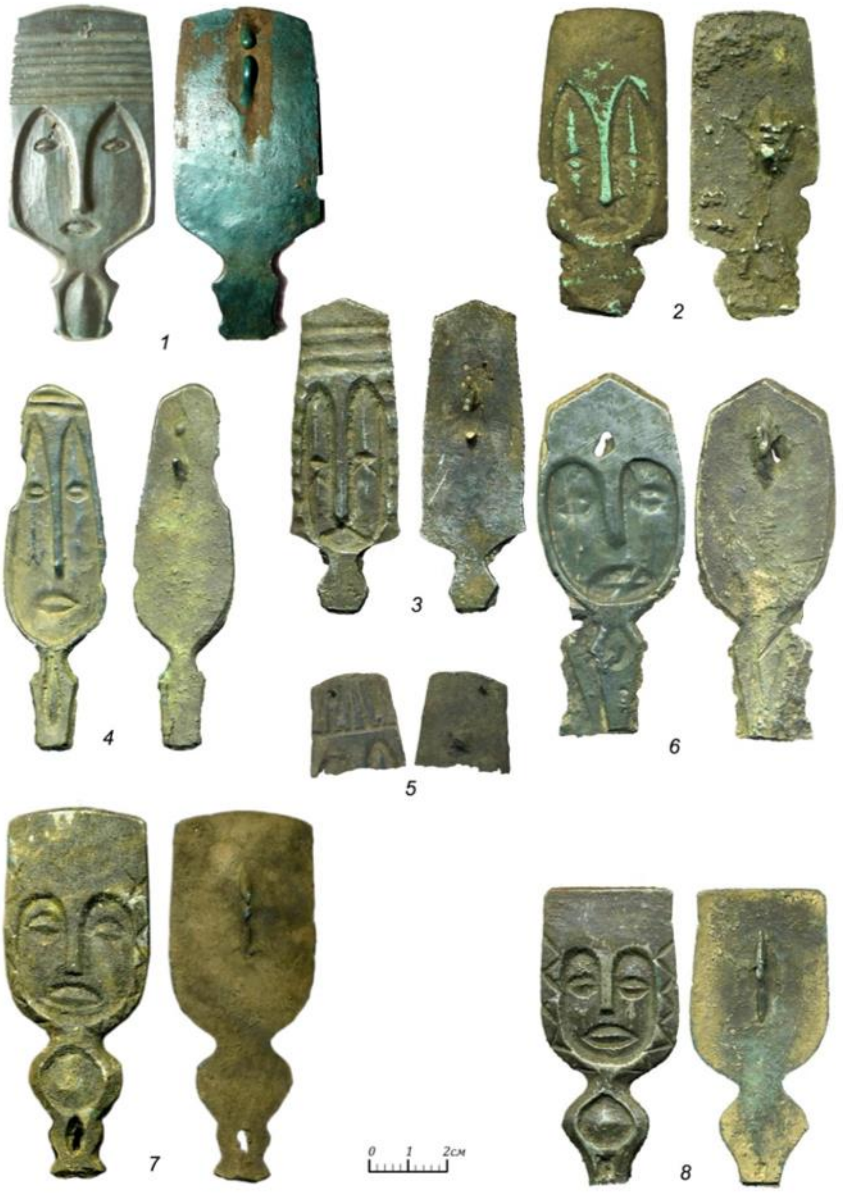


Рис. 1. Нечунаевское святилище. Металлопластика: 1-8 - бронза.

*Fig. S27.: Judino (proto-Ob-Ugric) cultic objects (10th–12th centuries), (after* B*OTALOV*–L*UKININ 2016. Fig. 6.)*

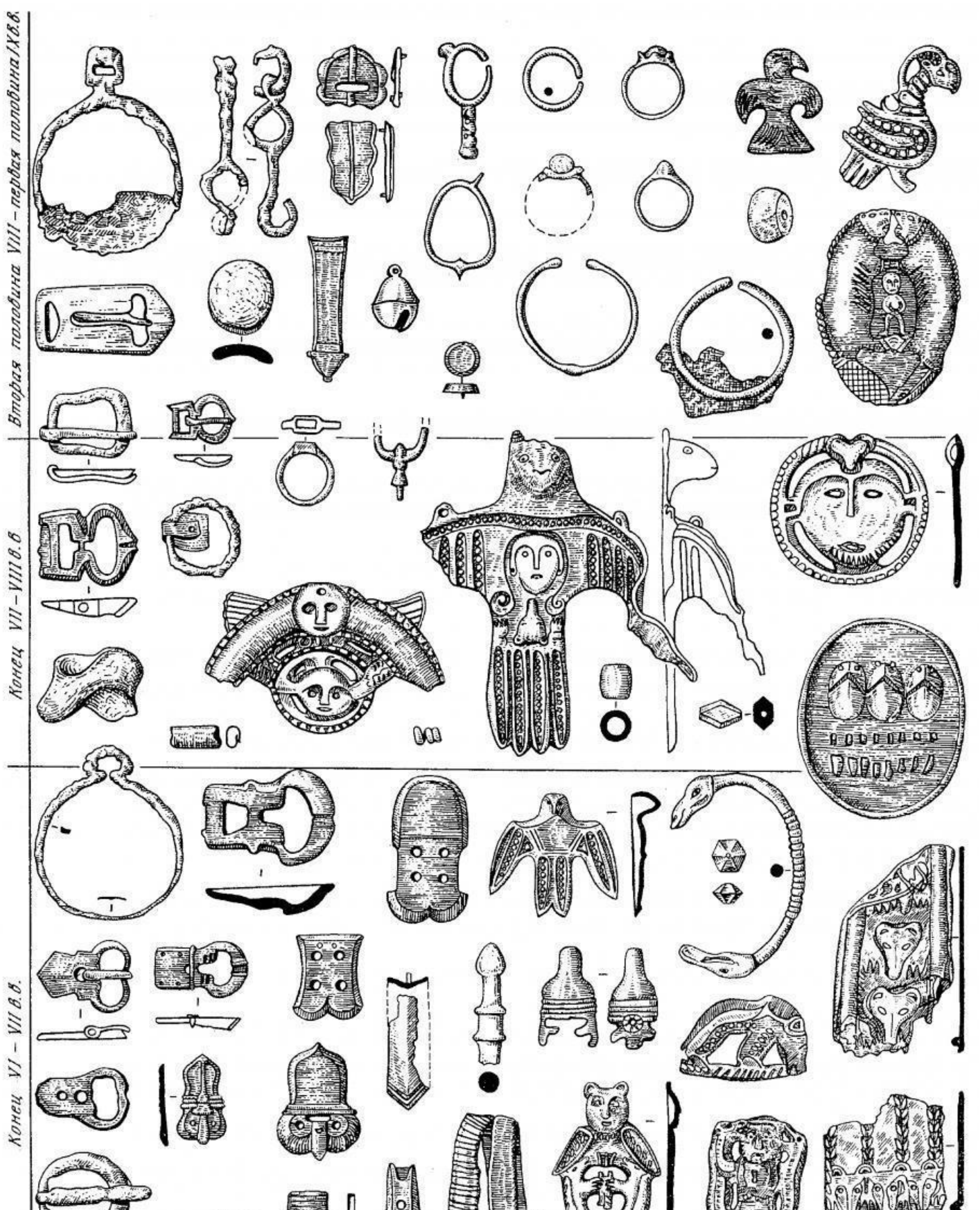


*Fig. S28: Typochronology and typical finds of the Potchevash culture (after* MOGILNIKOV *1987, Table LXXVIII)*

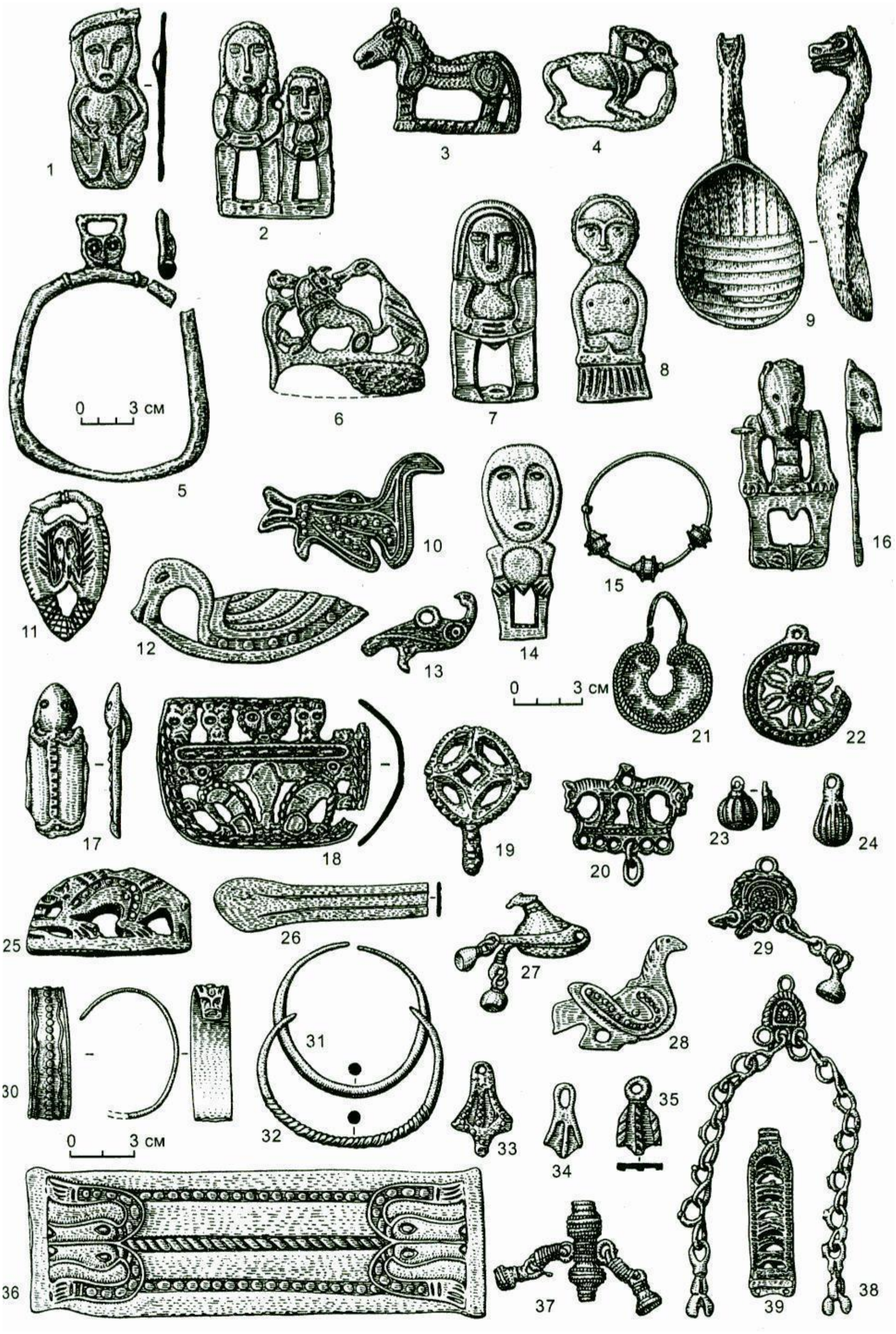


*Fig. S29: Typical finds of the Ust'-Ishim culture (after* MOGILNIKOV *1987, Table LXXXII)*

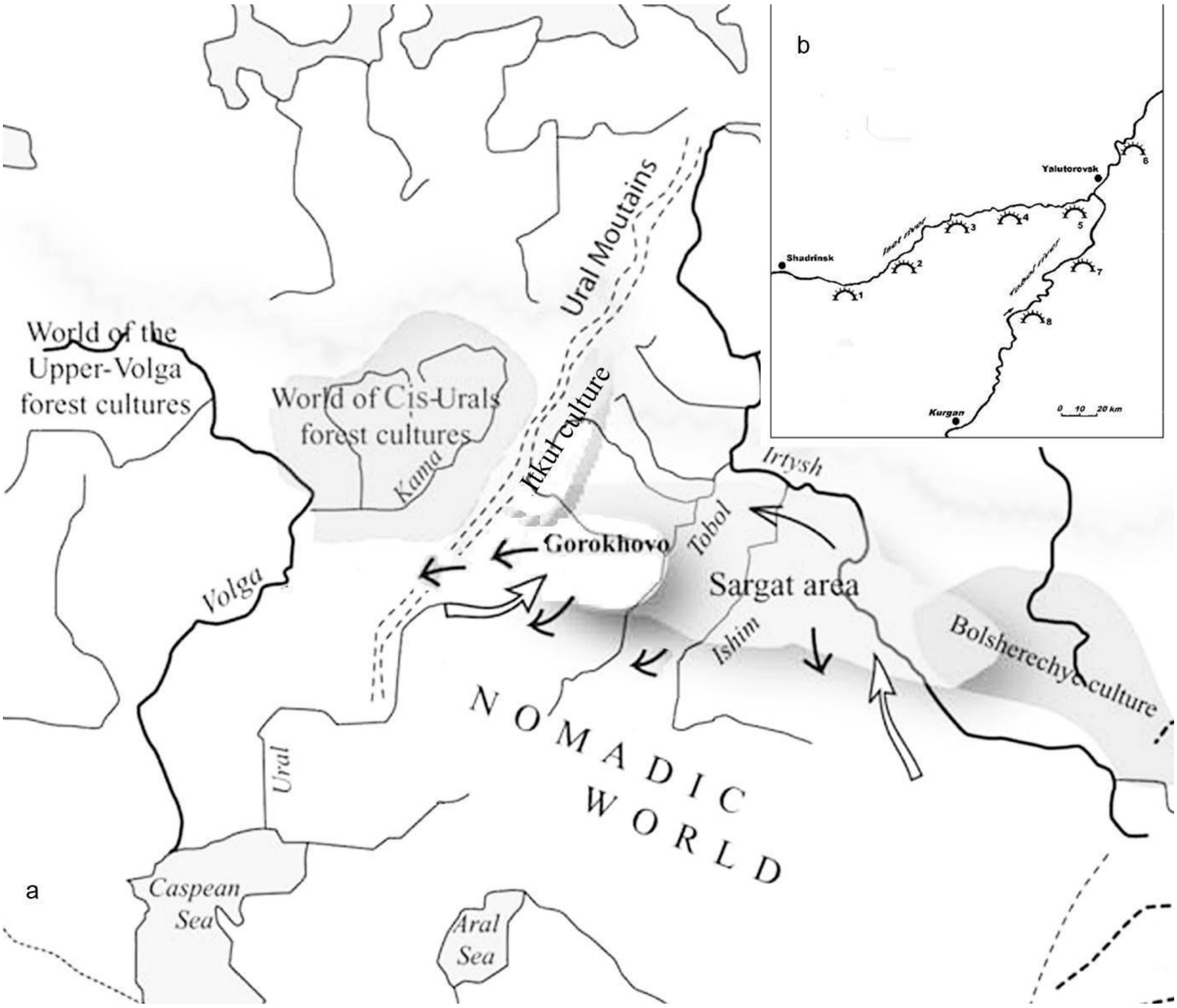


*Fig. S30: Archaeological cultures in the Late Iron Age Ural region (after* MATVEEVA *2018a, Fig. 1)*

#### 7. Bustanaevo

The Bustanaevo burial site is located at the southern part of the western slopes of the Urals, in the Burayevo District, the northern region of Bashkortostan, close to the Perm Krai, in the valley of the River Bistrii Tanip. After its discovery in 2011, the first surveys of the site were undertaken in 2015. It was in 2018 that systematic archaeological excavations started. The kurgan cemetery of Bustanaevo has been one of the greatest archaeological discoveries in the early medieval archaeology of the western slopes of the Urals in recent years. No other early Kushnarenkovo site (dated to the late sixth and early seventh centuries AD) has been found under authentic conditions since the early 1980s. According to the traditional historical view, the artefacts of this culture dated to this period belong to the first generations of the population moving here from the eastern side of the Urals (who are identified by most researchers with the early Hungarians). The finds discovered so far, particularly the so-called heraldic belt mounts (from the turn of the 6$^{th}$ and 7$^{th}$ centuries AD), horse equipment, weapons, and ceramic vessels belong to a rich site of the Kushnarenkovo culture (KOLONSKIKH 2020).

An outstanding assemblage of the site was found in Kurgan 45, which comprised a simple 0.7 m deep rectangular tomb of a woman, oriented NW–SE. The burial yielded mounts in the shape of four-petalled flowers made of white metal, a fragment of a bronze lyre-shaped buckle, a heraldic-style strap end, iron knives, riding gear, as well as two Kushnarenkovo ceramic vessels.

Close analogues of the archaeological finds discovered in the Bustanaevo kurgan cemetery are known in the South Urals from the early burials of the cemeteries of Manyak, Novo-Turbaslino, Birsk, Kushnarenkovo, Bahmuthino, and Novikovsk, as well as from the Novo-Bikkinsk Kurgan (MAZHITOV 1959, 1981, 2012, AKIMOVA–GENING 1959; SMIRNOV 1957). Most of the cemeteries above reflect mixed Kushnarenkovo, Bahmuthino, and Turbaslino traditions. Only the Manyak burial ground and the Novo-Bikkinsk Kurgan have a pure Kushnarenkovo character. Along with the latter two, the Bustanaevo kurgan cemetery represents the earliest Kushnarenkovo horizon. This was a period when the first representatives of this culture appeared at the western slopes of the Southern Urals.

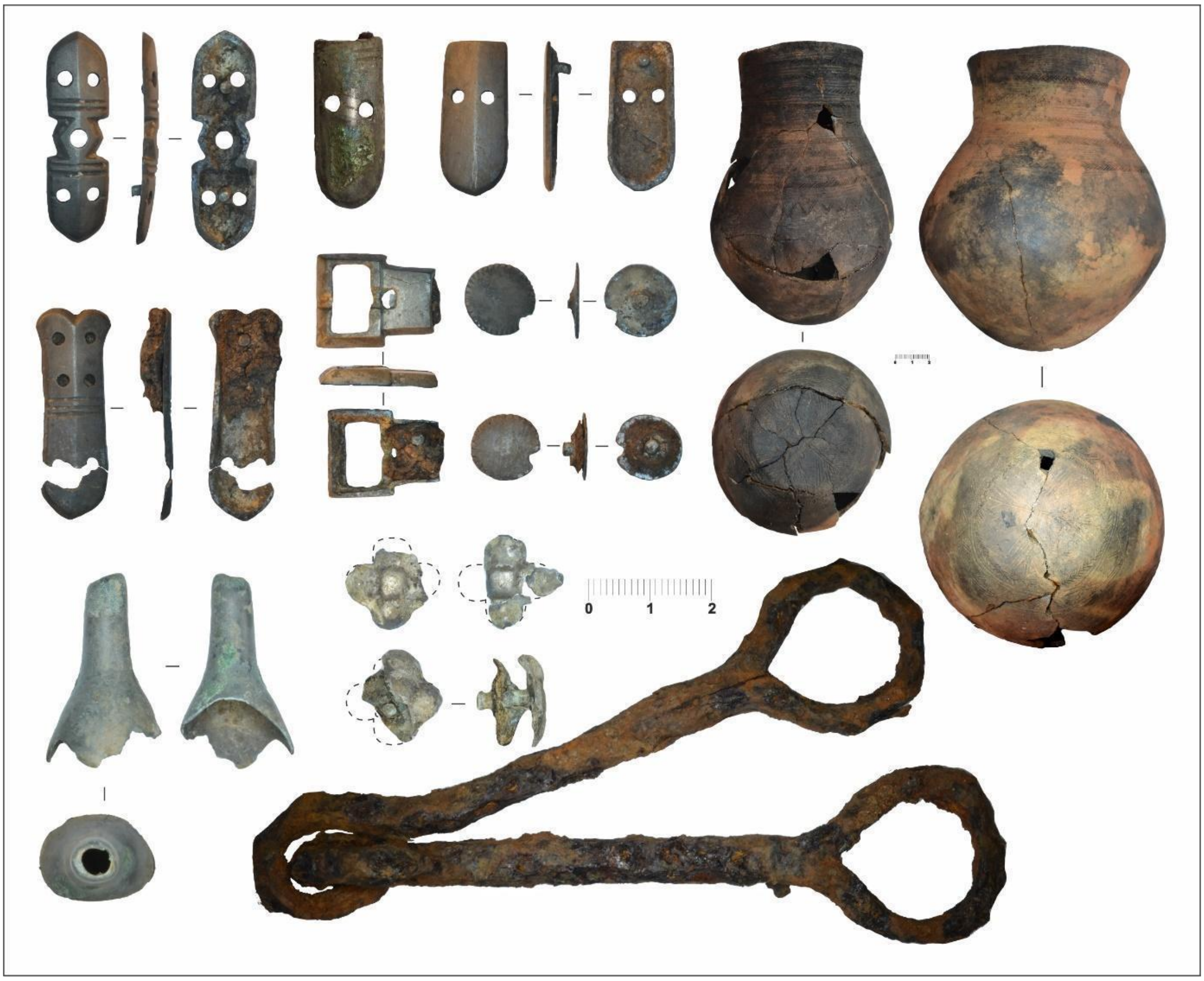


*Fig. S31. Some characteristic and typical finds from the early Kushnarenkovo culture Bustanaevo cemetery (photo by Aleksandr G. Kolonskikh)*

#### *8.* Classification *of 9-12$^{th}$ centuries cemeteries in the Carpathian Basin*

During the analysis of tenth- to twelfth-century burials discovered in the Carpathian Basin, the studies on genetics published so far used a dichotomy dividing the individuals into groups of "elites" and "common people" based on the grave-goods and burial customs. The manifestation of various historical processes was the idea underlying the division (NEPARÁCZKI ET AL. 2017, NEPARÁCZKI ET AL. 2018, MAÁR ET AL. 2021). However, archaeologists exploring this period no longer use this – fundamentally threefold (elites/middle class/common people) – division. Instead, they rely on a much more sophisticated classification using a clearly defined description, which is based on the times periods when the burial grounds were opened and used (KOVÁCS 2013, RÉVÉSZ 2020). The conceptual framework of the new classification has a much more specific definition, therefore the grouping of the graves/cemeteries is not based on the personal judgement of individual professionals. Instead, the decision and classification can be done by anyone with the help of a well-defined system. Comparing the results obtained in this way with the previous analyses (CSÁKY ET AL 2020, NEPARÁCZKI ET AL 2017, NEPARÁCZKI ET AL 2018, MAÁR ET AL. 2021), it can be clearly seen that the burials formerly regarded as "elite" can be largely classified as Kovács Type IV (so-called nomadic campsite cemeteries), and their genetic traits are very similar. Additionally, the majority of the published data mainly belong to groups IV–VI. In our present study, we use this new classification, and we carry out our analyses on this basis.

László Kovács divides the cemeteries of the examined period discovered in the Carpathian Basin into the following categories:

Type I: 9$^{th}$- to 12$^{th}$-century village cemeteries;

Type II: village cemeteries opened in the 9$^{th}$ century and still used in the 10$^{th}$ century;

Type IIIa: village cemeteries opened in the middle of the 9$^{th}$ century and abandoned in the 10$^{th}$ century (after the Hungarian conquest);

Type IIIb: cemeteries opened in the 9$^{th}$ century yielding Avar grave-goods, in the area of which graves dated to the 10$^{th}$ and 11$^{th}$ centuries were also discovered;

Type IV: 10$^{th}$-century (so-called) nomadic campsite cemeteries comprising a few graves;

Type V: 10$^{th}$-century village cemeteries;

Type VI: village cemeteries opened in the 10$^{th}$ century and used to the 11$^{th}$–12$^{th}$ centuries;

Type VII: 11th-century village cemeteries;

Type VIII: cemeteries opened in the 11$^{th}$ century around churches.

In our study, for reasons of simplification, we refer to the typological classification established by László Kovács, omitting the detailed description of the individual types.

#### *9*. Discussion of the radiocarbon dates

We found notable differences between the radiocarbon dates and the archaeological chronology in the case of the Western Siberian proto-Ob-Ugric sites. The radiocarbon results of bone samples from the burials of the archaeological cultures of Nizhny Novgorod (Barsov Gorodok, Ust-Tara-7), Potchevo (Vikulovo cemetery) and Ust-Ishim (Ivanov Cape 1, Panovo-1) are in most cases not synchronous with the relative dates.
We had the $d^{13}C$ and $d^{15}N$ isotopes also measured in the Poznan Radiocarbon Laboratory and in the Debrecen Isotoptech Laboratory (Supplementary Material Table S11) and experienced more negative values for the $d^{13}C$ and more positive ones for $d^{15}N$ on the proto-Ob-Ugric sites than is expected in a terrestrial environment (SCHOENINGER-DENIRO 1984).

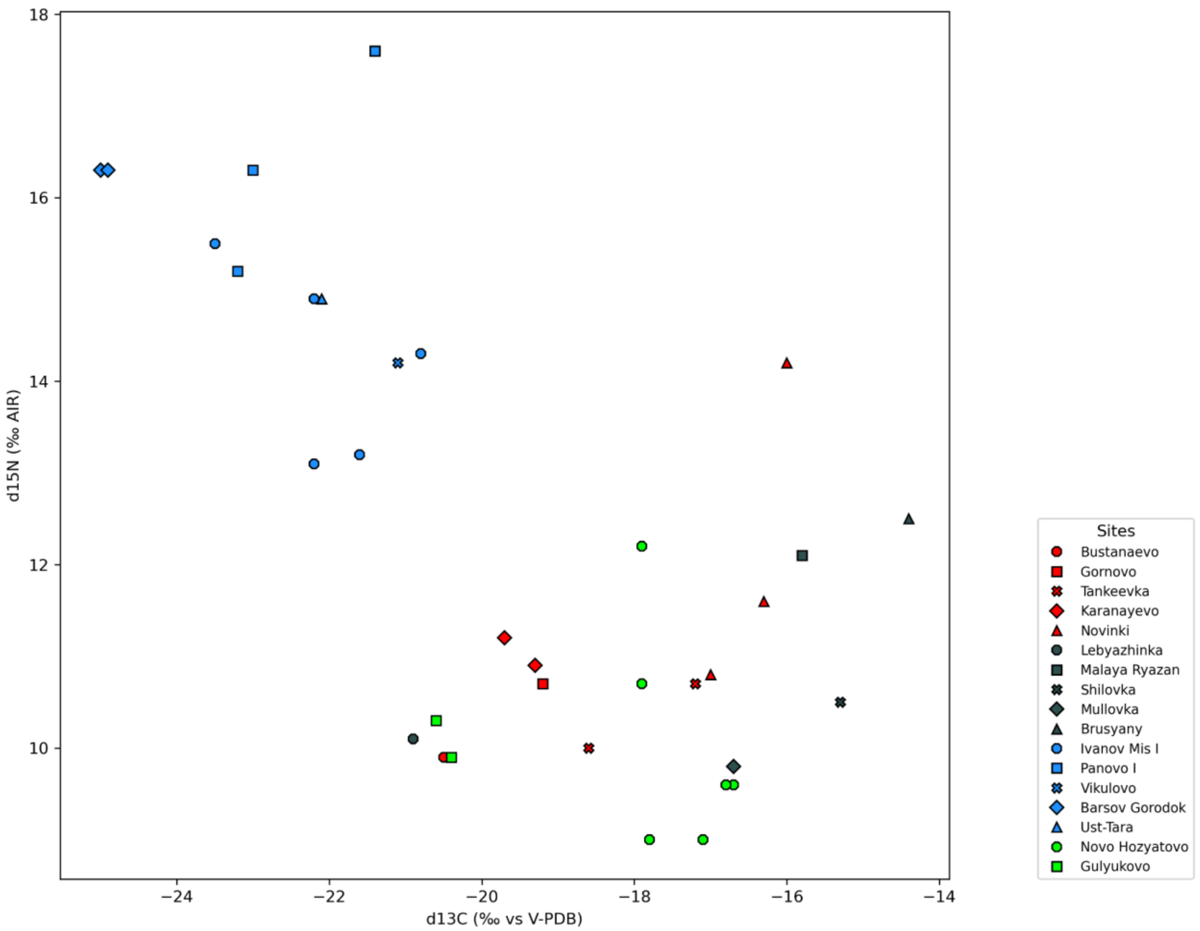


*Fig. S32. Scatterplot of δ13C and δ15N values of human bone collagen investigated in this study, presented by sites.*

The $d^{15}N$ values were elevated in these Western Siberian communities, except for the two samples from Ivanov Mis (kurgan 12 grave 1 and kurgan 10 grave 3) that also yielded radiocarbon dates in the expected range ($11^{th}$-$13^{th}$ centuries).

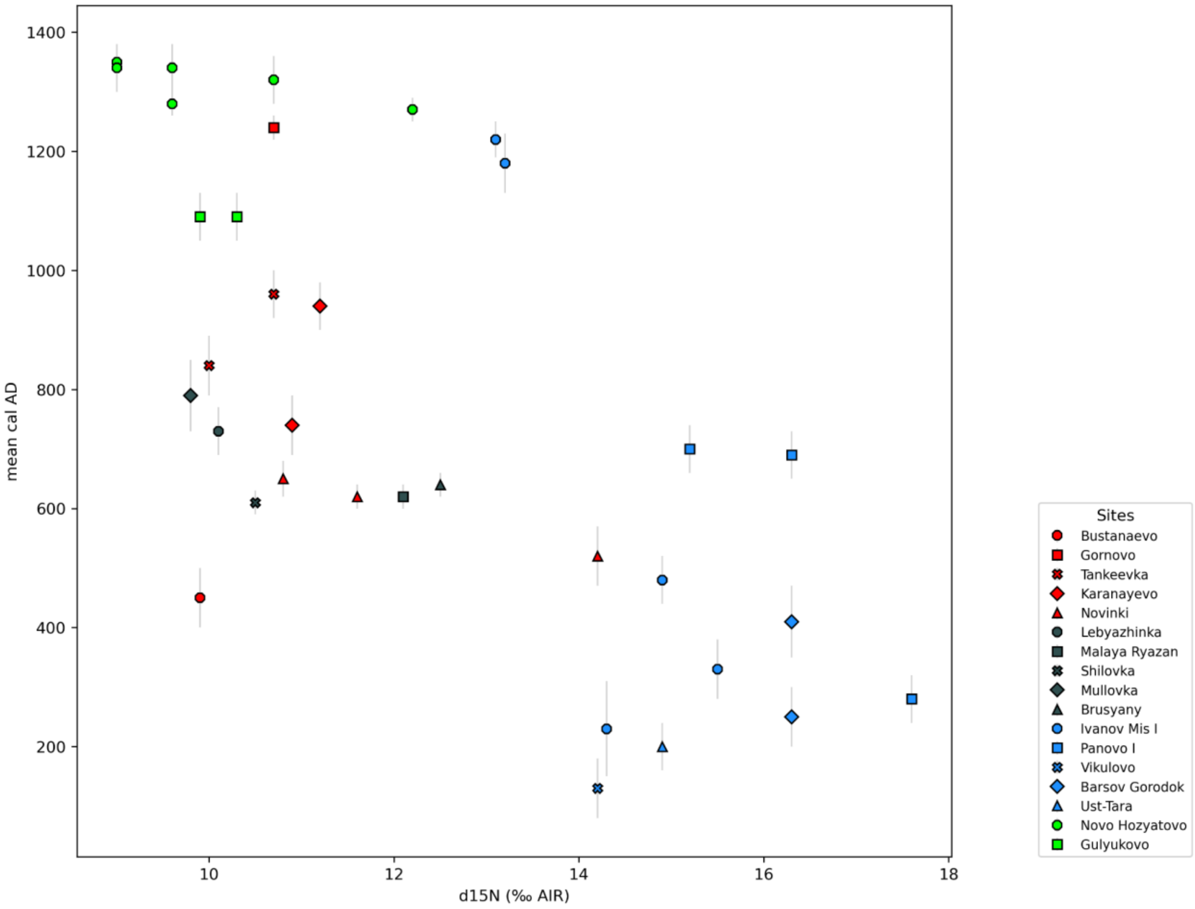


*Fig. S33. Scatterplot of absolute chronology (mean cal AD calculated in OxCal with sigma) and $\delta^{15}N$ values of human bone collagen investigated in this study, presented by sites.*

A similar trend where human samples with more depleted $\delta^{13}C$ values have older radiocarbon ages, e.g., the difference between archaeological age and radiocarbon age increases is not quite consistent; however, most of the Western Siberian samples have depleted $\delta^{13}C$ values. These can also be connected to a freshwater fish-based diet; however, the fishes' isotope signatures vary to such an extent (MARCHENKO ET AL. 2021) that small studies would be needed from the study areas for a detailed evaluation of the stable isotope results.

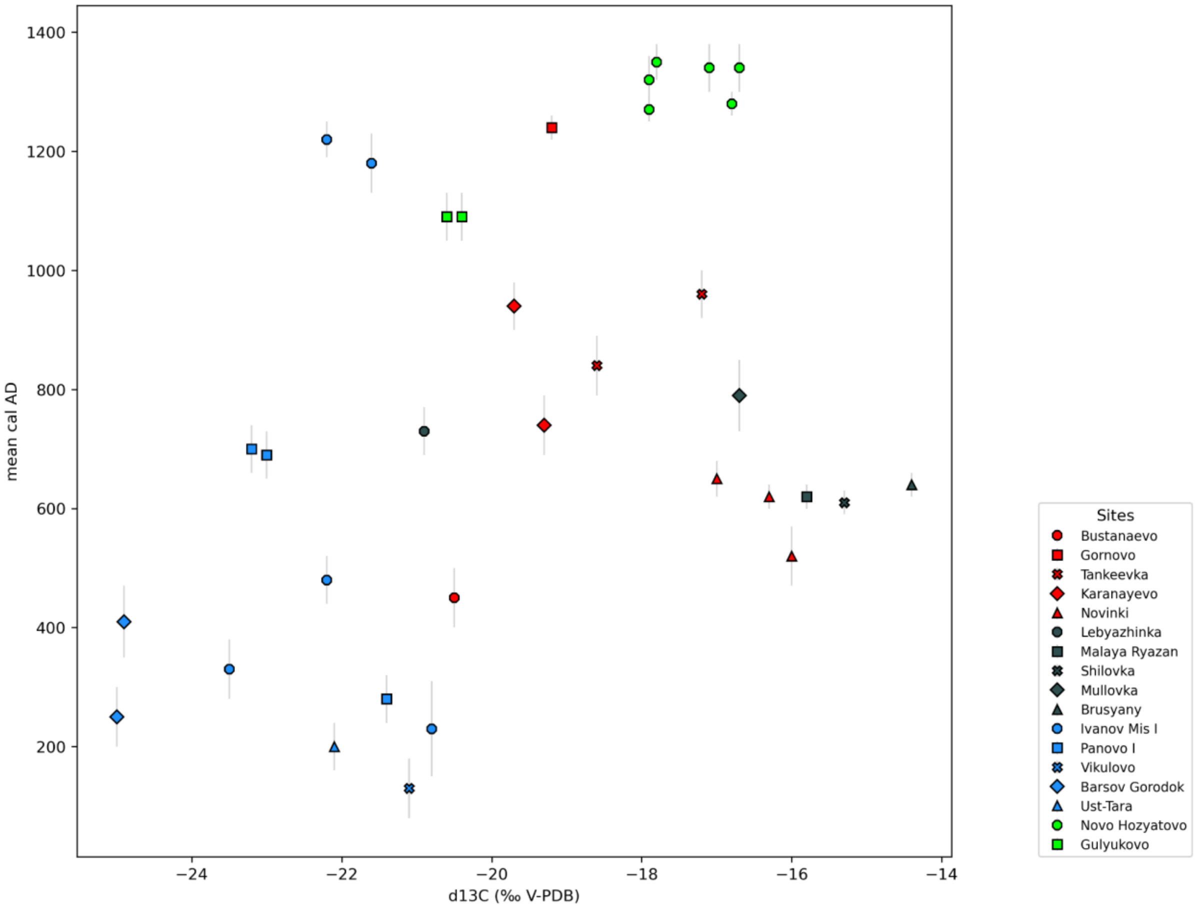


*Fig. S34. Scatterplot of absolute chronology (mean cal AD dates calculated in OxCal with sigma) and $\delta^{13}C$ values of human bone collagen investigated in this study, presented by sites.*

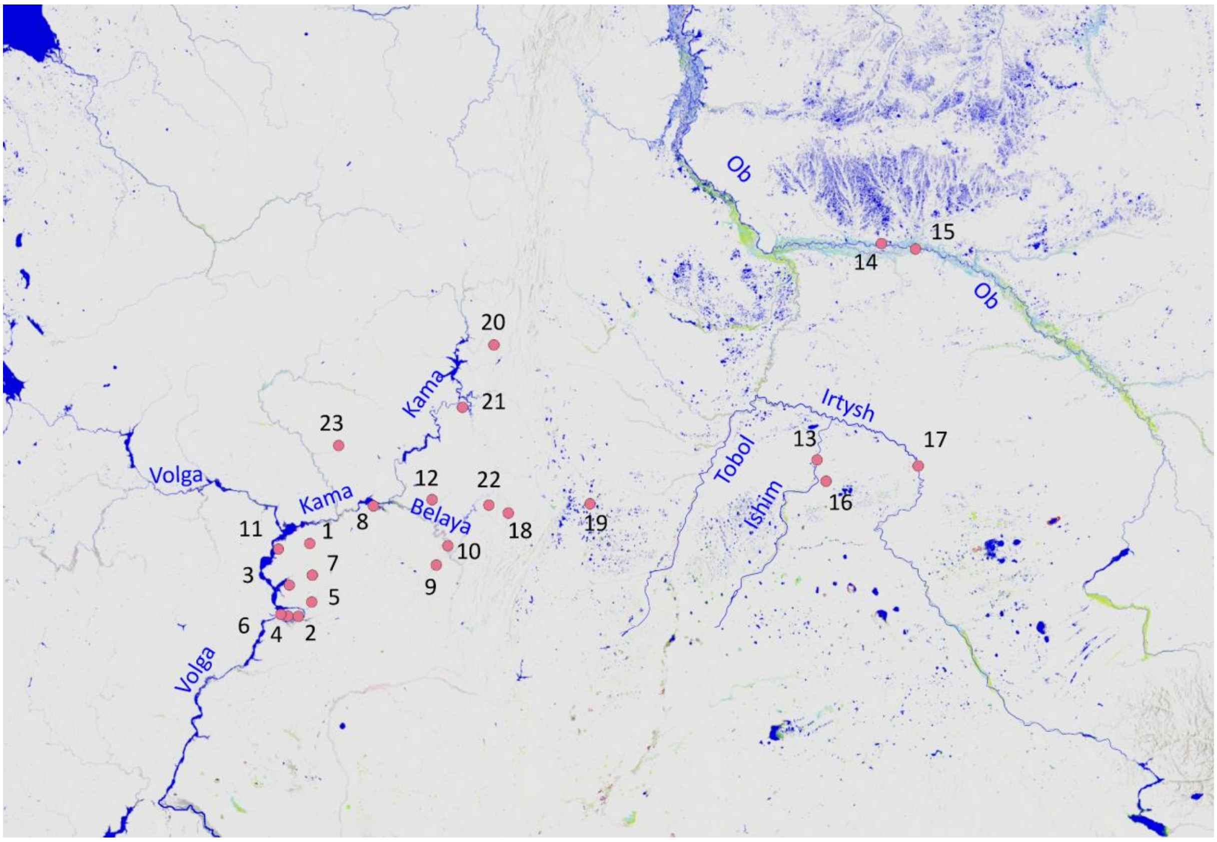


*Fig. S35. Map with the sites, zooming-in to the southern Ural region. Bolshie Tigani (1); Novinki group: Novinki (2), Mulovka (3), Brusyany (4), Lebyazhinka (5), Malaya Ryazan (6), Shilovka (7); Chiyalik group: Gulyukovo (8), Novo Hozyatovo (9), Gornovo (10); Tankeevka (11); Bustanaevo (12); Proto Ob Ugric group: Vikulovo (13), Barshov Gorodok (14), Ivanov Mis (15), Panovo (16), Ust-Tara (17); Uyelgi+Karanayevo group: Karanayevo (18), Uyelgi (19); Cis-Ural group: Bayanovo (20), Brody (21), Bartim (22), Sukhoy Log (23) (source of 19-23.: CSÁKY ET AL. 2020).*

In our opinion, these radiocarbon dates are a clear example of the distorting effect often observed while radiocarbon dating the bones of domestic animals (such as dogs and pigs) in the taiga and tundra parts of Western Siberia (KUZMIN ET AL. 2020). This phenomenon is probably due to the dietary characteristics of the population groups living close to river systems in Western Siberia, based on the consumption of large quantities of freshwater fish, which had a major contribution to the so-called freshwater reservoir effect (FRE) (LOSEY ET AL. 2018). FRE depend not just on the type of animal or plant consumed, but also on the age and type of carbonates or organic material in the watershed. Rivers like Ob (along which sites Ivanov Mis and Barsov Gorodoc are located) have limestone bedrock that can also affect the isotope values ("hard water effect") of the freshwater food consumed by the studied communities (PHILIPPSEN 2013). Lacking other local comparative data (wood, animal bones, and others), FRE cannot be reliably suggested, but just hypothesised in this paper.

### 10. References to chapter A


AKIMOVA–GENING 1959: Акимова, М. С. – Генинг, В. Ф.: Отчёт об исследованиях археологических памятников у с. Кушнаренково БАССР в 1959 г. Научный архив ИЭИ УФИЦ РАН. Ф. 1. Оп. 6. Д. 14.

BAGAUTINOV–BOGACHEV–ZUBOV 1998: Багаутдинов, Р. С. – Богачев, А. В. – Зубов, С. Э.: *Праболгары на Средней Волге (у истоков истории татар Волго-Камья).* Самара 1998.

BORZUNOV–CHEMYAKIN 2014: Борзунов, В. А. – Чемякин, Ю. П.: Карымские могильники таежного Приобья. *Вестник археологии, антропологии и этнографии* 25:2 (2014) 64–70.

BOTALOV 2016: Боталов С. Г.: Историко-культурные горизонты в эпоху раннего железного века и средневековья лесо-степного Зауралья (Historical-cultural horizons of the early Iron and Middle Ages in the Trans-Urals forest-steppe). In: Археология Южного Урала. Лес, лесостепь (проблемы культурогенеза). Ред.: Боталов, С. Г. Челябинск 2016, 468–553.

BOTALOV–LUKININ 2016: Боталов, С. Г. –Лукиных А. А.: Святилищный комплекс у села Нечунаево. In: Археология Южного Урала. Лес, лесостепь (проблемы культурогенеза). Ред.: Боталов, С. Г. Челябинск 2016, 427–442.

BUKINA-LIFANOV-ZUBOV 2018: Букина О.В., Лифанов Н.А., Зубов С.Э. Раскопки могильника Малая Рязань I в 2017 г. // Вопросы археологии Поволжья. Вып. 7. Отв.ред. М.А. Турецкий. Самара: СГСПУ, 2018. 209–214;

BUKINA-LIFANOV-ZUBOV-BAGAUTDINOV 2020: Букина О.В., Лифанов Н.А., Зубов С.Э., Багаутдинов Р.С. Исследования могильника Малая Рязань I в 2019 г. // Археологические открытия в Самарской области 2019 года. Отв.ред. Д.А. Сташенков. Самара, 2020, 27–28;

CHALIKOVA–CHALIKOV 1981 Chalikova, E. A. – Chalikov, A. H.: Altungarn an der Kama und im Ural (Das Gräberfeld von Bolschie Tigani). Régészeti Füzetek Ser. II. 21. Budapest 1981.

CHEMYAKIN–KARACHAROV 2002: Чемякин, Ю. П. – Карачаров, К. Г.: Древняя история Сургутского Приобья. In: *Очерки традиционного землепользования хантов (материалы к атласу). 2-е изд., испр. и доп*. Екатеринбург 2002, 5–74.

CHEMYAKIN–ZIKOV 2004: Чемякин, Ю. П. – Зыков, А. П.: *Барсова Гора: Археологическая карта. Сургут–Омск*. Омск 2004.

CHINDINA 2001: Чиндина, Л. А.: Кулайская культура. In: *Народы и культуры Томско-Нарымского Приобья: Материалы к энциклопедии Томской области*. Томск 2001, 78–81.

CSÁKY, V. ET AL 2020*:* Csáky, V., Gerber, D., Szeifert, B., Egyed, B., Stégmár, B., Botalov, S.G., Grudochko, I.V., Matveeva, N.P., Zelenkov, A.S., Sleptsova, A.V., *et al.* Early medieval genetic data from Ural region evaluated in the light of archaeological evidence of ancient Hungarians. *Sci. Rep.* **10**, (2020).

FJODOROVA ET AL. 1991: Федорова, Н. В. – Зыков, А. П. – Морозов, В. М. – Терехова, Л. М.: Сургутское Приобье в эпоху средневековья. Вопросы археологии Урала. 20. Екатеринбург 1991.

FODOR 1972: Фодор, И.: К вопросу о погребальном обряде древних венгров. In: *Проблемы археологии и древней истории угров*. Ред.: Смирнов, А. П. и др. Москва 1972, 176–188.

FODOR 1977: Fodor I.: Bolgár-török jövevényszavaink és a régészet [Bulgarian Turkic loan words in the Hungarian language and archaeology]. In: Magyar őstörténeti tanulmányok [Studies on Hungarian prehistory]. Ed.: Bartha A. – Czeglédy K. – Róna-Tas A. Budapest 1977, 79–114.

FODOR 2015: Фодор, И.: Венгры: древняя история и обретение Родины. Пермь 2015.

GARUSTOVICH 1988: ГАРУСТОВИЧ, Г. Н.: ОБ ЭТНИЧЕСКОЙ ПРИНАДЛЕЖНОСТИ РАННЕМУСУЛЬМАНСКИХ ПАМЯТНИКОВ ЗАПАДНОЙ И ЦЕНТРАЛЬНОЙ БАШКИРИИ. IN: ПРОБЛЕМЫ ДРЕВНИХ УГРОВ НА ЮЖНОМ УРАЛЕ. УФА 1988, 130–139.

GAZIMZYANOV 1995: Газимзянов И.Р. Новые данные по антропологии населения Самарского Поволжья в эпоху раннего средневековья // Средневековые памятники Поволжья. Отв.ред. Г.И. Матвеева. Самара: Самарский университет, 1995, 95–109.

GAZIMZYANOV-KHOKHLOV 1999: Газимзянов И.Р., Хохлов А.А. Черепа из средневековых захоронений на территории древнего поселения Лебяжинка V // Вопросы археологии Поволжья. Вып.1. Самара, 1999.

GENING 1977: Генинг, В. Ф.: Проблема происхождения венгров. Советская археология 1977:1, 317–321.

GENING–ZDANOVICH 1987: Генинг, В. Ф. – Зданович, С. Я.: Лихачевский могильник на р. Ишим-памятник потчевашской культуры VI–VIII вв. н.э. In: *Ранний железный век и средневековье.* Челябинск 1987, 119–133.

GOLDINA 2013: Голдина, Р. Д.: Некоторые замечания относительно формирования теории угорского присутствия в Предуралье в эпоху средневековья. В: *II-й Международный Мадьярский симпозиум.* Отв. ред.: Боталов, С. Г. – Иванова, Н. О. Челябинск 2013, 89–109.

HALIKOVA 1972: Халикова, Е. А.: Погребальный обряд Танкеевского могильника и его венгерские параллели. In: *Проблемы археологии и древней истории угров*. Ред.: Смирнов, А. П. и др. Москва 1972, 145–160.

HALIKOVA 1976: Халикова, Е. А.: Ранневенгерские памятники Нижнего Прикамья и Приуралья. Советская археология 1976:3, 141–156.

HALIKOV 1984: Халиков, А. Х.: Новые исследования Больше-Тиганского могильника (о судьбе венгров, оставшихся на древней Родине). In: Проблемы археологии степей Евразии. Ред.: Мартынов, А. И. Кемерово 1984, 122–133.

HLEBNIKOVA 1984: ХЛЕБНИКОВА, Т. А.: *КЕРАМИКА ПАМЯТНИКОВ ВОЛЖ-СКОЙ БОЛГАРИИ. К ВОПРОСУ ОБ ЭТНОКУЛЬТУРНОМ СО-СТАВЕ НАСЕЛЕНИЯ.* МОСКВА 1984.

ILYUSHINA 2009: Илюшина, В. В.: Подчевашские погребения из грунтового могильника «Викуловское кладбище». *Вестник археологии, антропологии и этнографии* 10 (2009) 74–82.

KARACHAROV 2004: Карачаров, К. Г.: *Раскопки могильника Барсов Городок (Барсовский I) в 1989 году: отчет о НИР / УрГУ.* Архив АСА. Ф. 1. Д. 143. Екатеринбург 2004.

KARACHAROV 1993: Карачаров, К. Г.: Хронология раннесредневековых могильников Сургутского Приобья. In: *Хронология памятников Южного Урала: Сборник статей.* Уфа 1993, 110–118.

KAZAKOV 1971: Казаков, Е. П.: Погребальный инвентарь Танкеевского могильника. In: *Вопросы этногенеза тюркоязычных народов Среднего Поволжья.* Ред.: Халиков, А. Х. Казань 1971, 94–155.

KAZAKOV 1987: Казаков, Е. П.: О происхождении и культурной принадлежности памятников с гребенчато-шнуровой керамикой. In: *Проблемы средневековой археологии Урала и Поволжья.* Уфа 1987.

KAZAKOV 1989: КАЗАКОВ, Е. П.: О ВЗАИМОДЕЙСТВИИ БОЛГАРО-САЛТОВСКОГО И ПРИУРАЛЬСКОГО НАСЕЛЕНИЯ (ПО МАТЕ-РИАЛАМ КЕРАМИКИ ВОЛЖСКОЙ БОЛГАРИИ). IN: *РАННИЕ БОЛГАРЫ В ВОСТОЧНОЙ ЕВРОПЕ*. ОТВ. РЕД.: ХА-ЛИКОВ, А. Х. КАЗАНЬ 1989, 122–133

KAZAKOV 2007: *Волжские булгары, угры и финны: Проблемы взаимодействия*. Казань 2007.

KHALIKOVA– KAZAKOV 1977: Khalikova, E. A. – Kazakov, E. P.: Le cimetière de Tankeevka. In: *Les anciens hongrois et les ethnies voisines a l'est.* Budapest. Ed. Erdélyi, I. 1977.

KOLONSKIKH 2020: Колонских, А. Г.: Научный отчёт об итогах археологических раскопок Бустанаевского курганного могильника на территории Бураевского района Республики Башкортостан в 2019 году. Уфа 2020 // Научно-отраслевой архив ИА РАН.Ф-1. Р-1. № 64776

KOMAR 2001: *Комар, А. В.:* К вопросу о дате и этнокультурной принадлежности Шиловских курганов: [Самар. обл.]. In: *Степи Европы в эпоху средневековья* 2 (2001) 11–44.

KOMAR 2018: Комар, А.: История и археология древних мадьяр в эпоху миграции – A korai magyarság vándorlásának történeti és régészeti emlékei [Historical and archeological data of the migration of the early Hungarians]. Eds.: Türk, A. – Budai, D. Studia ad Archaeologiam Pazmaniensia 11. – Magyar Őstörténeti Témacsoport Kiadványok 5. Budapest 2018

KONIKOV 2007: Коников, Б. А.: Омское Прииртышье в раннем и развитом Средневековье. Омск 2007.

KONIKOV 2019: Коников, Б. А.: *Иванов-Мыс-I курганный могильник. Тевризский район, Омская область.* Омск 2019.

KOVÁCS 2013: Kovács, L.: *A Kárpát-medence honfoglalás és kora Árpád-kori szállási és falusi temetői.* In Révész, L. (ed.), A honfoglalás kor kutatásának legújabb eredményei, Szeged, pp. 511–604.

RÉVÉSZ 2020: Révész, L. (2020) *A 10–11. századi temetők regionális jellemzői a Keleti-Kárpátoktól a Dunáig*. Martin Opitz Kiadó: Budapest.

KUZMIN ET AL. 2020: Kuzmin, Y. V., Kosintsev, P. A., Boudin, M., Zazovskaya, E. P. The freshwater reservoir effect in northern West Siberia: 14C and stable isotope data for fish from the late medieval town of Mangazeya. *Quat Geochronol* 2020;**60**:101109.

LIFANOV 2005: Лифанов, Н. А.: К вопросам периодизации и хронологии памятников новинковского типа. In: *Степи Европы в эпоху средневековья* 4 (2005) 25–40.

LIFANOV 2020: Лифанов, Н. А.: Малая Рязань I, курганно-грунтовый могильник. In: Культурное наследие Самарской области. Гл. ред., Сташенков, Д. А. Самара 2020, 270.

LOSEY ET AL 2018: Losey, R. J., Fleming, L. S., Nomokonova, T., Gusev, A. V., Fedorova, N. V., Gervie-Lok, S.,Bachura, O. P.,Kosintsev, P. A.,Sablin, M. V. Human and Dog Consumption of Fish on the Lower Ob River of Siberia: Evidence for a Major Freshwater Reservoir Effect at the Ust'-Polui Site. *Radiocarbon* 2018;**60**:239–60.

MAÁR ET AL. 2021: Maár, K., Varga, G.I.B., Kovács, B., Schütz, O., Maróti, Z., Kalmár, T., Nyerki, E., Nagy, I., Latinovics, D., Tihanyi, B., *et al.* Maternal lineages from 10-11th century commoner cemeteries of the Carpathian Basin. *Genes (Basel).* 12, (2021).

MARCHENKO ET AL. 2021: Marchenko, Z. V., Svyatko, S. V., Grishin A. E. δ13C and δ15N isotope analysis of modern freshwater fish in the south of Western Siberia and its potential for palaeoreconstructions. *Quat Int* 2021;**598**:97–109.

MATVEEVA 1997: Матвеева, Г. И.: *Могильники ранних болгар на Самарской Луке*. Самара. 1997.

MATVEEVA 2018a: Матвеева, Н. П.: Саргатская и гороховская культуры: исторические судьбы. Ранний железный век западносибирской лесостепи по археологическим данным (A szargatkai és gorohovói kultúra történeti interpretációja. A korai vaskor Nyugat-Szibéria erdős sztyeppi területén a régészeti adatok alapján). In: *III-й Международный мадьярский симпозиум по археологии. Будапешт, 6–10 июня 2016 г. – 3. Nemzetközi Korai Magyar Történeti és Régészeti Konferencia. Budapest, 2016. június 6–10.* Ред.: Türk, A. – Зеленков, A. C. Studia ad Archaeologiam Pazmaniensia 12. – Magyar Őstörténeti Témacsoport Kiadványok 6. Budapest 2018, 15–34.

MATVEEVA 2018b: Матвеева, Н.П.: О миграциях из Западной Сибири в Европу в раннем железном веке и в эпоху Великого переселения народов. In: Археология евразийских степей. 2018. №6. 150–154.

MAZHITOV 1959: Мажитов, Н. А.: Курганный могильник в деревне Ново-Турбаслы. In: Башкирский археологический сборник. Под ред. А. П. Смирнова и Р. Г. Кузеева. Уфа 1959, 114–142.

MAZHITOV 1977: МАЖИТОВ, Н. А.: *ЮЖНЫЙ УРАЛ В VII–XIV ВВ.* МОСКВА 1977

MAZHITOV 1981: Мажитов, Н. А.: Курганы Южного Урала VIII–XII вв. н.э. Москва 1981, 105–109.

MAZHITOV 2012: Мажитов, Н. А.: Башкортостан в IV–VIII вв. In: История башкирского народа: в 7 т. 2. Гл. ред.: Кульшарипов, М. М. Уфа 2012.

MOGILNIKOV 1987: Могильников, В. А.: Угры и самодийцы Урала и Западной Сибири. In: *Финно-угры и балты в эпоху средневековья.* Ред.: Седов, В. В. Археология СССР. Москва 1987, 163-235 https://www.archaeolog.ru/media/

MOROZOV 2004: МОРОЗОВ, В. М.: ИСТОРИЯ ИЗУЧЕНИЯ ПЕТРОГРОМСКОЙ КУЛЬТУРЫ. IN: *ЧЕТВЕРТЫЕ БЕРСОВСКИЕ ЧТЕНИЯ*. ОТВ. РЕД.: КОВАЛЕВА, В.Т. ЕКАТЕРИНБУРГ 2004, 110–114.

NEPARÁCZKI ET AL. 2017: Neparáczki, E., Kocsy, K., Tóth, G.E., Maróti, Z., Kalmár, T., Bihari, P., Nagy, I., Pálfi, G., Molnár, E., Raskó, I., *et al.* Revising mtDNA haplotypes of the ancient Hungarian conquerors with next generation sequencing. PLoS One 12, 1–11 (2017).

NEPARÁCZKI ET AL. 2018: Neparáczki, E., Maróti, Z., Kalmár, T., Kocsy, K., Maár, K., Bihari, P., Nagy, I., Fóthi, E., Pap, I., Kustár, Á., *et al.* Mitogenomic data indicate admixture components of Central-Inner Asian and Srubnaya origin in the conquering Hungarians. PLoS One 208 (2018).

PASTUSHENKO 2011: ПАСТУШЕНКО, И. Ю.: ВОЗМОЖНО ЛИ ГОВОРИТЬ ОБ «УГОРСКОЙ ЭПОХЕ В ПРИКАМЬЕ? ВЕСТНИК УДМУРТСКОГО УНИВЕРСИТЕТА. СЕРИЯ ИСТОРИЯ И ФИЛОЛОГИЯ 2001:1, 144–150.

PHILIPPSEN 2013: Philippsen B. The freshwater reservoir effect in radiocarbon dating. *Herit Sci* 2013;**1**:1–19

POSHEHONOVA 2010: Пошехонова, О. Е.: Краниологические особенности средневековых популяций Сургутского Приобья (по материалам могильников с Барсовой Горы). *Вестник археологии, антропологии и этнографии.* 13 (2010:2) 91–97.

RAHMANALIJEV 2009: Рахманалиев, Р: Империя тюрков. Великая цивилизация. Москва 2009.

SCHOENINGER-DENIRO 1984: Schoeninger M.J., DeNiro M.J. Nitrogen and carbon isotopic composition of bone collagen from marine and terrestrial animals. *Geochim Cosmochim Acta* 1984;**48**:625–39.

SKANDAKOV–DANCHENKO 1999: Скандаков, И. Е. – Данченко, Е. М.: Курганный могильник Усть-Тара-VII в южнотаежном Прииртышье. In: *Гуманитарное знание. Серия «Преемственность». Ежегодник. Сборник научных трудов*. Омск 1999:3, 160–186.

SMIRNOV 1957: Смирнов, А. П.: Железный век Башкирии. In: Культура древних племен Приуралья и Западной Сибири. Отв. ред.: Мерперт, Н. Я. Москва 1957, 5–113.

STASHENKOV 1995: Сташенков, Д. А.: Особенности погребального обряда хазарского времени в Среднем Поволжье. In: *Культуры степей Евразии второй половины I тысячелетия н.э. Тезисы докладов Международной научной археологической конференции. 14–17 ноября 1995 г.* Самара 1995, 83–85.

STASHENKOV–TURETSKII 1999: Сташенков, Д. А. – Турецкий, М. А.: Погребение эпохи раннего средневековья у хутора Лебяжинка (к вопросу об этнокультурной ситуации в Самарском Поволжье в IX в.). In: *Вопросы археологии Поволжья.* Вып. 1. Отв. ред.: Выборнов, А. А. Самара, 1999, 289–302.

SUNGATOV 2016: Сунгатов, Ф. А.: Каранаевский курганный могильник. In: Археология Южного Урала. Лес, лесостепь (проблемы культурогенеза). Ред.: Боталов, С. Г. Челябинск 2016, 409–426.

TEREHOVA–KARACHAROV 1994: Терехова, Л. М. – Карачаров, К. Г.: Среднеобская низменность, п.5.6. In: *Очерки культурогенеза народов Западной Сибири 2. Мир реальный и потусторонний*. Томск 1994, 277–289.

VASILEV–MATVEEVA 1986: Васильев, И. Б. – Матвеева, Г. И.: *У истоков истории Самарского Поволжья*. Куйбышев 1986.

VIKTOROVA 1968: ПАСТУШЕНКО, И. Ю.: ВОЗМОЖНО ЛИ ГОВОРИТЬ ОБ «УГОРСКОЙ ЭПОХЕ В ПРИКАМЬЕ? ВЕСТНИК УДМУРТСКОГО УНИВЕРСИТЕТА. СЕРИЯ ИСТОРИЯ И ФИЛОЛОГИЯ 2001:1, 144–150.

VIKTOROVA–MOROZOV 1993: ПАСТУШЕНКО, И. Ю.: ВОЗМОЖНО ЛИ ГОВОРИТЬ ОБ «УГОРСКОЙ ЭПОХЕ В ПРИКАМЬЕ? ВЕСТНИК УДМУРТСКОГО УНИВЕРСИТЕТА. СЕРИЯ ИСТОРИЯ И ФИЛОЛОГИЯ 2001:1, 144–150.

ХАЛИКОВА–ХАЛИКОВ 2018: Халикова, Е. А. – Халиков, А. Х.: Раниие венгры на Каме и Урале (Больше–Тиганский могильник). Археология евразийских степей 25. Казань 2018.

ZELENKOV 2018: Зеленков, А. С.: Историко-культурная модель эпохи средневековья Тоболо-Иртышской провинции (A Tobol–Irtis régió középkorának történeti és régészeti-kulturális modellje). In: *III-й Международный мадьярский симпозиум по археологии. Будапешт, 6–10 июня 2016 г. – 3. Nemzetközi Korai Magyar Történeti és Régészeti Konferencia. Budapest, 2016. június 6–10.* Ред.: Türk, A. – Зеленков, А. С. Studia ad Archaeologiam Pazmaniensia 12. – Magyar Őstörténeti Témacsoport Kiadványok 6. Budapest 2018, 35–46.

### Chapter B:
### Genetic analyses

#### 1. Population genetic analyses

##### 1.1. PCA analysis

We performed PCA analyses based on mitochondrial haplogroup frequencies of archaic and modern-day populations (Supplementary Material, Table S3). We visualised the results in two-dimensional plots. Plots based on PC1-PC2 (main text Fig. 4) and PC1-PC3 values (Fig. S36) show consistent results. Uyelgi+Karanayevo, Chiyalik, Tankeevka, Bolshie Tigani, and Novinki groups are close to each other, but the PC3 axis separates them.

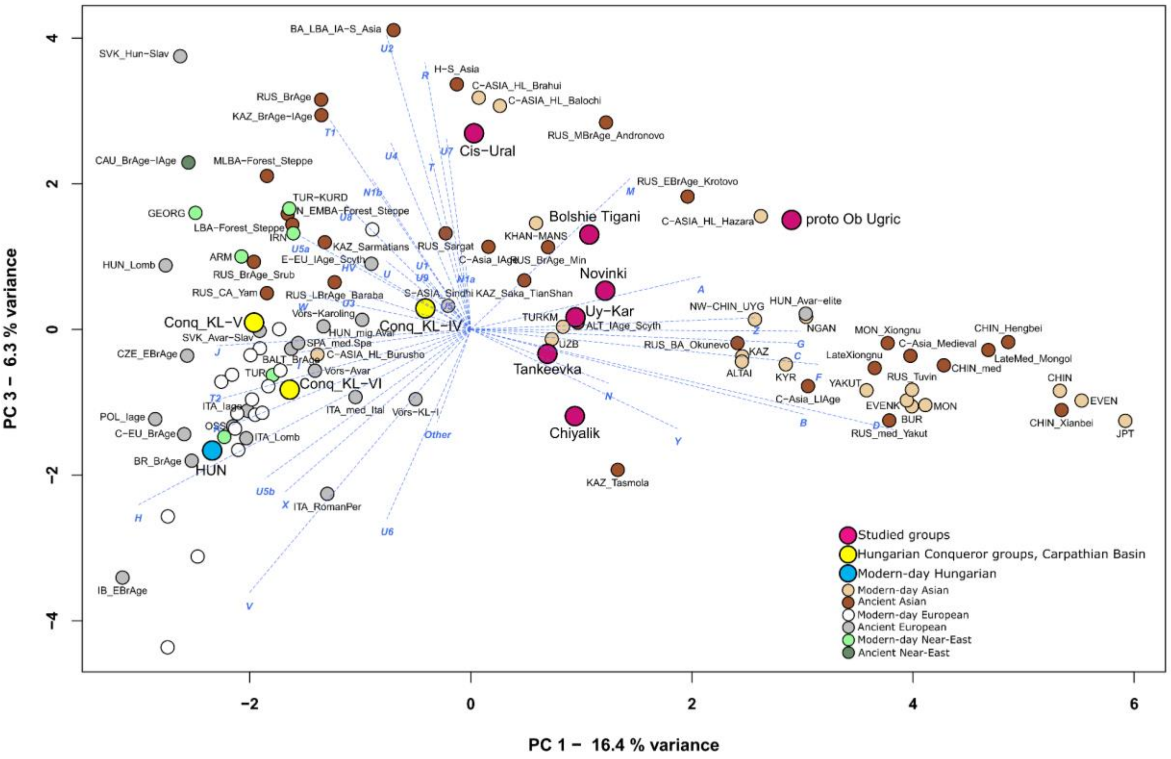


*Figure S36: PCA plot representing PC1 and PC3. We compared the studied groups (magenta) with ancient and modern-day populations from Eurasia and Near East. The variance of the components is as follows: PC1: 16.1%; PC3: 6%. For more information about the populations see Supplementary Table S3.*

##### 1.2. Ward clustering

Ward cluster analysis (Fig. S37) was performed on the populations of the PCA analysis, based on PC1-6 values. The results show that our studied groups are on a bigger sub-branch, together with steppe groups mainly. The proto-Ob-Ugric group forms a small sub-branch with modern Nganasan and it is a sister-branch of the Uyelgi+Karanayevo. Tankeevka, Chiyalik, Bolshie Tigani, Novinki, Cis-Ural and RUS_Sargat groups form another sub-branch whose sister-branch consists of the KL-IV and KL-V groups. The KL-VI group is on another branch among European populations.

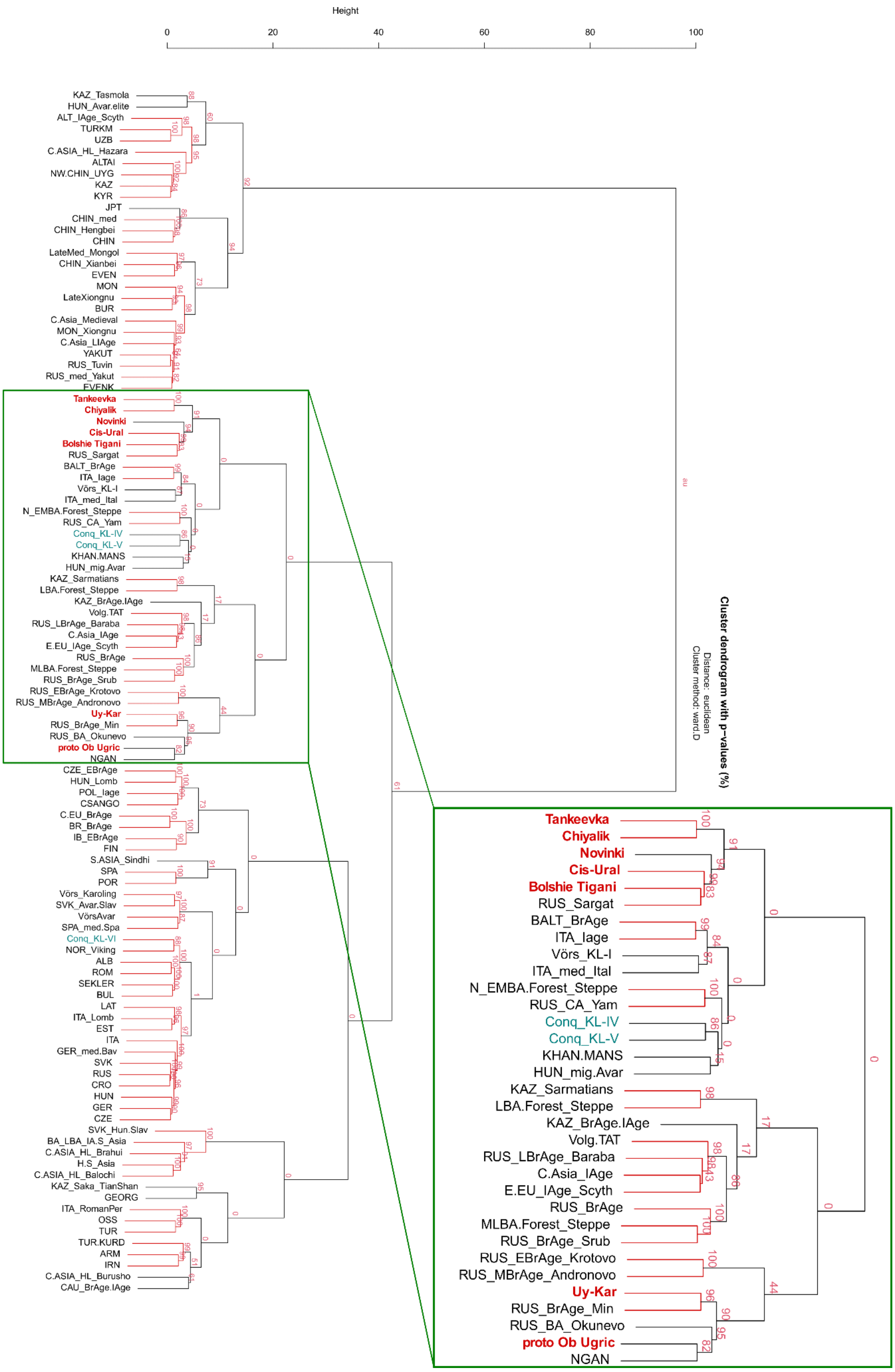


*Figure S37: Results of Ward cluster analysis based on ancient and modern-day population datasets. We highlighted the sub-branch containing our examined groups. For more information about the populations, see Supplementary Table S3. For the larger image in PDF format, see Extended Data Figure 1.*

##### 1.3. $F_{ST}$ analyses

We calculated $F_{ST}$ and linearized Slatkin $F_{ST}$ (SLATKIN 1995) in Arlequin and performed clustering based on these genetic distance values. We compared the 7 studied groups from the

Volga-Ural region and Western-Siberia and the three conqueror groups with other ancient and modern-day Eurasian and Near-Eastern populations. The result of this analysis is shown in the heatmap in Fig. S38, and the exact $F_{ST}$ values are seen in Supplementary Material, Table S4.

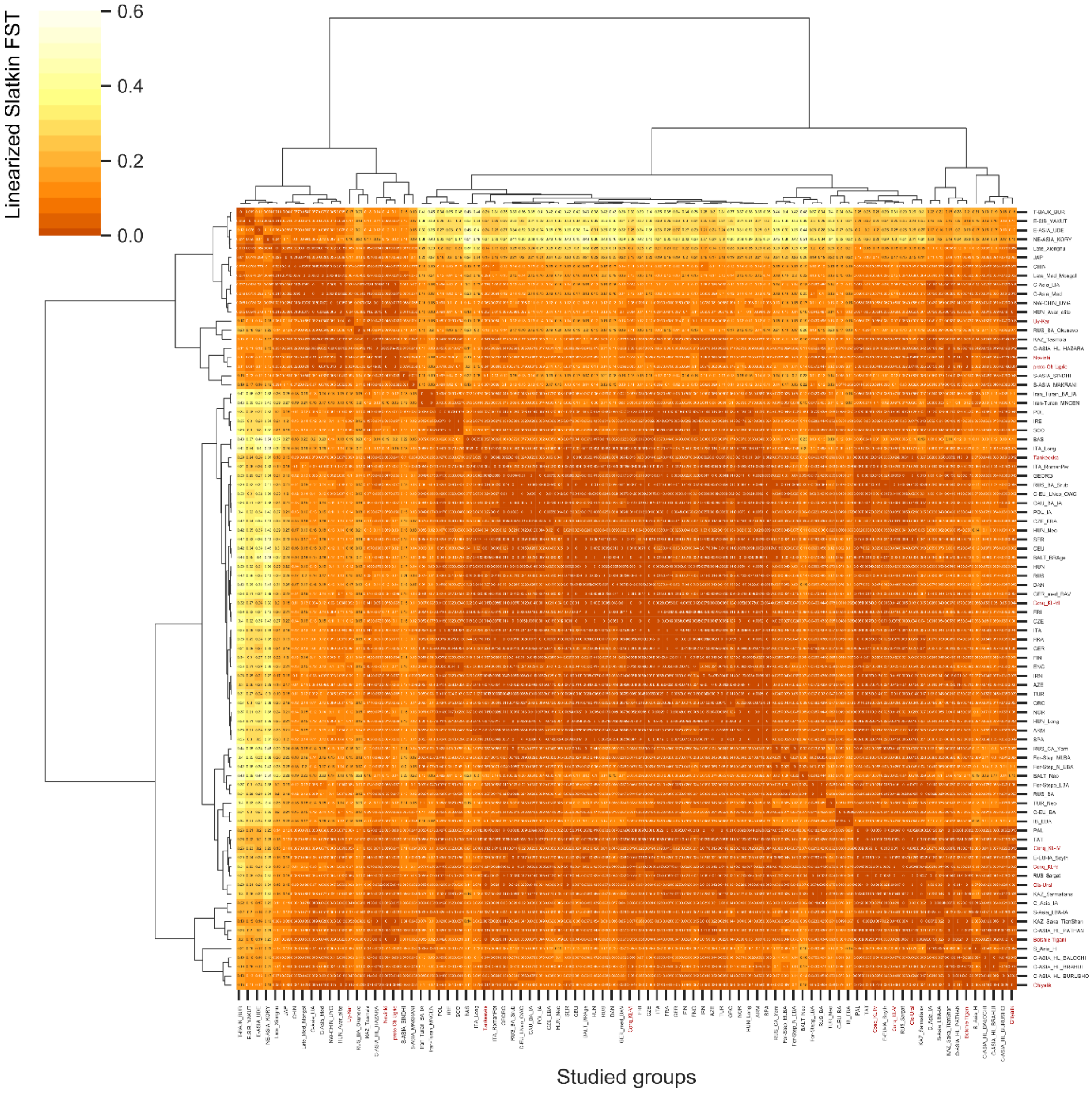


*Figure S38: Heatmap and clustering based on linearized Slatkin $F_{ST}$. For the larger image in PDF format, see Extended Data Figure 2.*

##### 1.4. AMOVA analyses

We performed AMOVA analyses (Analysis of Molecular Variance) in Arlequin, using the 10 formed groups (main text Table 1, Supplementary Material, Table S4, Table S5).

First, we classified these groups into three sets based on the clustering results: 1) Uyelgi+Karanayevo, proto-Ob-Ugric, Novinki; 2) Tankeevka, Cis-Ural, KL-IV, KL-V, KL-VI and 3) Bolshie Tigani, Chiyalik. In this case, the source of variance among sets (4.06%) is more than within the sets (0.83%), and among the sets, we detected a significant difference ($F_{CT}$= 0.04058, p= 0.00782±0.00313), while the difference between groups within sets is not significant ($F_{SC}$= 0.00869, p= 0.15836±0.01353).

Secondly, we formed four sets from the studied groups: 1) Uyelgi+Karanayevo, proto-Ob-Ugric, Novinki; 2) Tankeevka, KL-VI; 3) Cis-Ural, KL-IV, KL-V; 4) Bolshie Tigani, Chiyalik). Then we detected that the source of variance is the same (2.44%) among and within sets; moreover, in this case, the difference is significant between groups within the sets ($F_{CT}$= 0.02439, p= 0.05963+-0.00813) ($F_{SC}$=0.02496, p= 0.00098+-0.00098).

Thus, the AMOVA analyses support the division of the studied groups into three sets.

##### 1.5. MDS analysis based on Slatkin $F_{ST}$ values

We performed MDS (Multidimensional Scaling) based on linearized Slatkin $F_{ST}$ values (SLATKIN 1995) and plotted in 2D. The groups from the Volga-Ural region are mainly South- and Central-Asian archaic (Iron Age, Bronze Age, Medieval period) and modern-day populations. The position of the Novinki and proto-Ob-Ugric groups also shows a closer connection with the eastern population. The outsider position of the Uyelgi+Karanayevo group could be caused by the numerous intra- and intersite genetic connections.

The groups of Hungarian conquerors from the Carpathian Basin (KL-IV-VI) are separated from each other. The KL-IV group is located between the European and Asian populations, the closest groups to it being the Cis-Ural and RUS_Sargat (Iron Age Sargat culture in the Southern Ural). This conqueror group is the closest to our studied groups (Tankeevka and Cis-Ural groups based on dimension 1 and Bolshie Tigani and Chiyalik group based on dimension 2). The KL-IV is near the steppe-origin populations and the KL-VI close to European and Near-Eastern groups (Fig. S39).

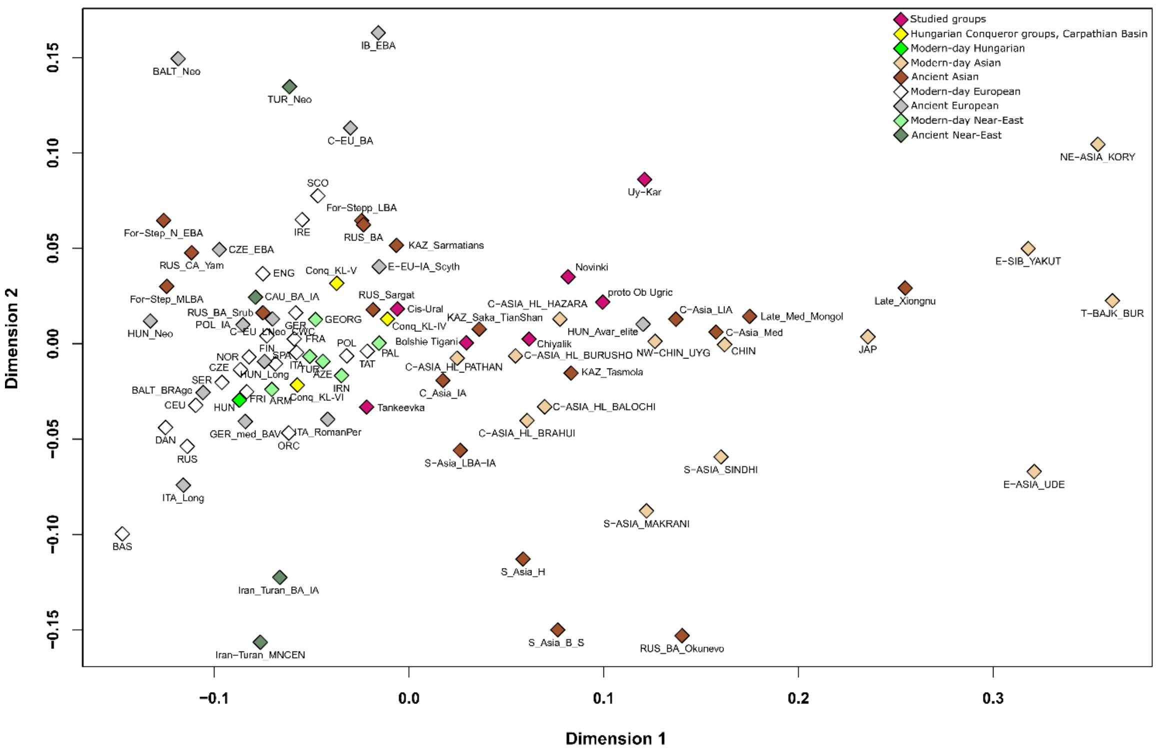


*Figure S39: MDS plot based on whole mitogenome sequences of ancient and present-day populations. The analysis was made using Slatkin $F_{ST}$ values, which are presented in Table S4.*

##### 1.6. Mantel test

We performed Mantel-tests (MANTEL 1967) based on the genetic and geographical distances of the studied groups and the comparative data (Supplementary Material, Table S6).

We that the two variables correlate with each other on Eurasia scale. However, if groups at great geographical distance (like KL-IV-VI and/or proto-Ob-Ugric) were not included in the analysis, the two variables become uncorrelated. Thus, the diversity between the studied groups in the Volga-Ural region is not the result of their spatial distribution, but rather intensive mobility and interaction of populations shaped the early medieval maternal genetic landscape of the Volga-Ural region.

#### 2. Individual-level analyses

##### 2.1. Y-STR network analyses

The median-joining network of the R1a-Z93>Z94>Z2124 (nevgen.org) Y-chromosome subhaplogroup was constructed with 16 STRs (Fig. S40). We used 3 of our samples for this network, which belong to the Bolshie Tigani, and Novinki (two samples from the Brusyany site) groups. According to our results the Novinki samples are related to each other, but the Bolshie Tigany sample clusters distantly, showing no paternal connection between the two groups.

The median-joining network of another R1a subhaplogroup (R1a-Z280>Y4459) was performed based on 15 STRs (Fig. S41). This network contains three studied samples, which represent the Chiyalik culture. The two paternal haplotypes from the Novo Hozyatovo site are identical and are one step away from modern Belorussians and Russians. The sample from the Gulyukovo site is distal from these samples, but it is also located near the Russian individuals.

The haplogroup of the Hungarian King Béla III is R1a-Z282>Z280>> L1280> Y5647 based on his STR profile (OLASZ ET AL. 2018). One of our samples from the Novinki group (Lebyazhinka site) is also included in this subhaplogroup. According to the median-joining network of the subhaplogroup R1a-Z282>Z280>> L1280 (Fig. S42), Béla III and Lebyazhinka samples are close to each other, but several steps apart, which is caused by the difference observed at several loci between them. They are grouped with Eastern-European people (the dataset used for the network analysis can be found in Supplementary Material, Table S10).

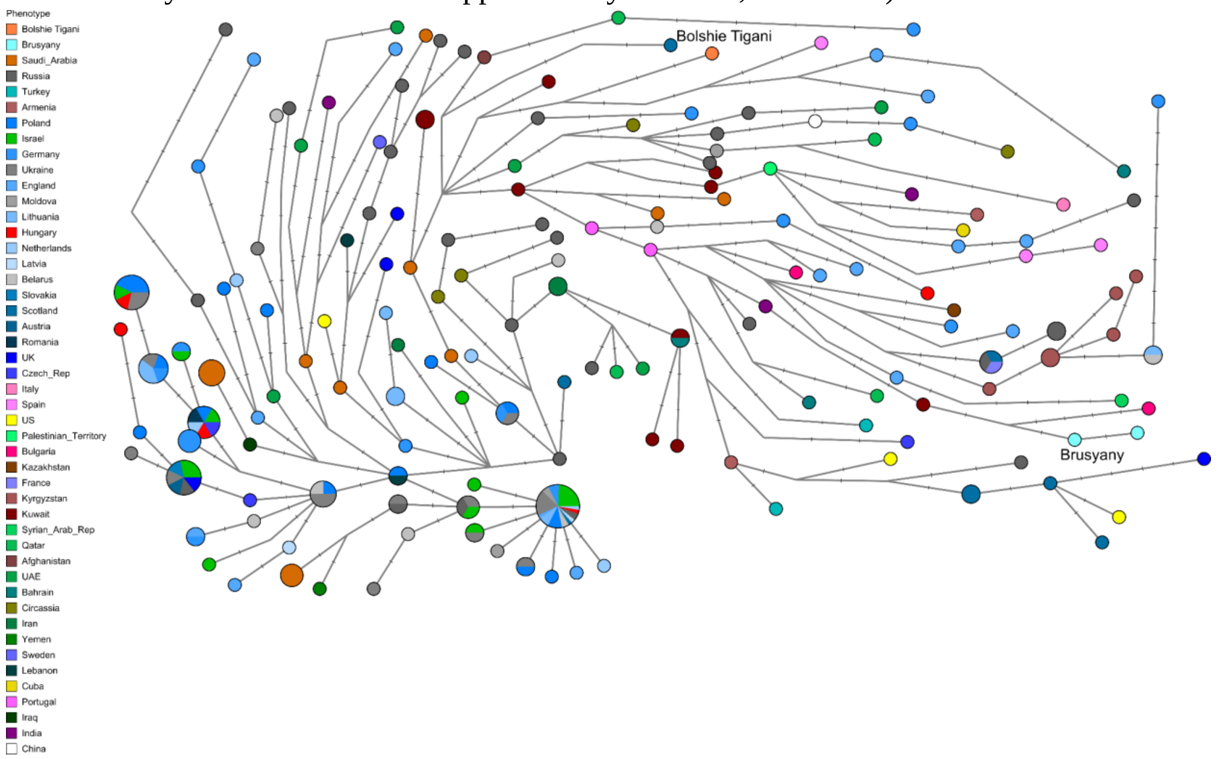


*Figure S40: Median-joining network of R1a Z93>Z94>Z2124 Y-chromosomal subhaplogroup. This network was created using 16 STRs.*

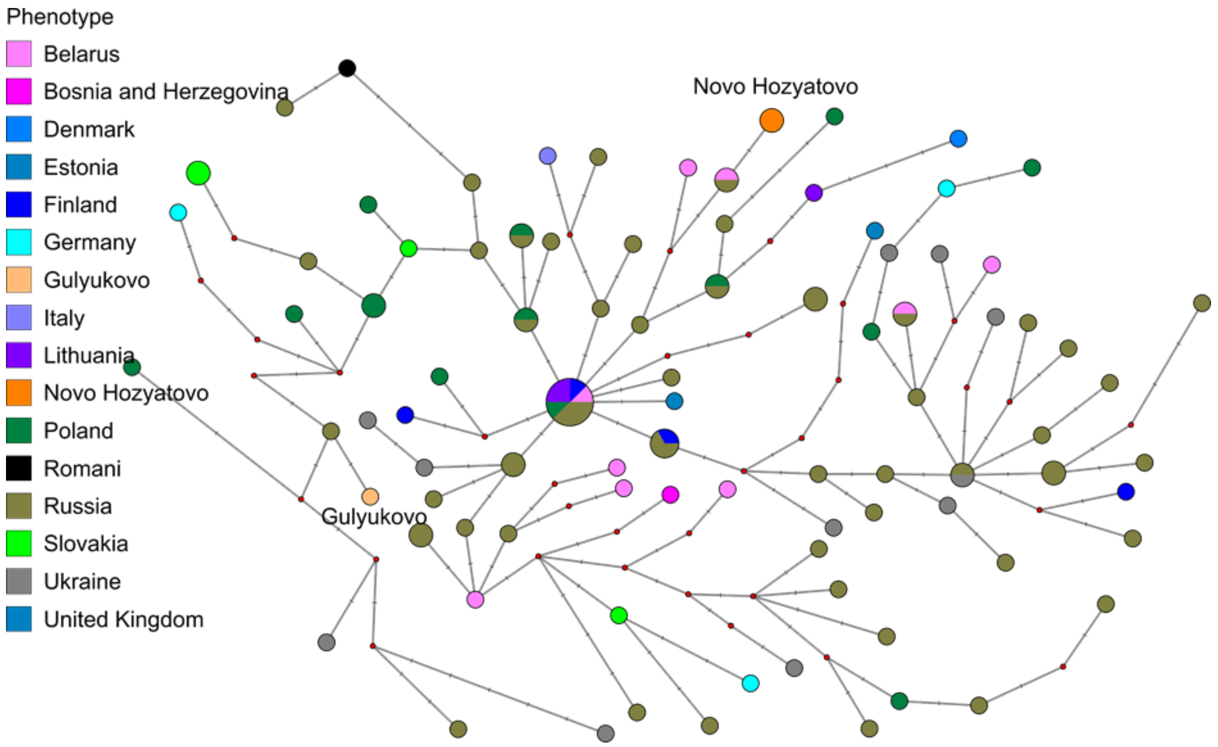


*Figure S41: Median-joining network of R1a-Z280>Y4459 Y-chromosomal subhaplogroup. The analysis was performed based on 15 STRs.*

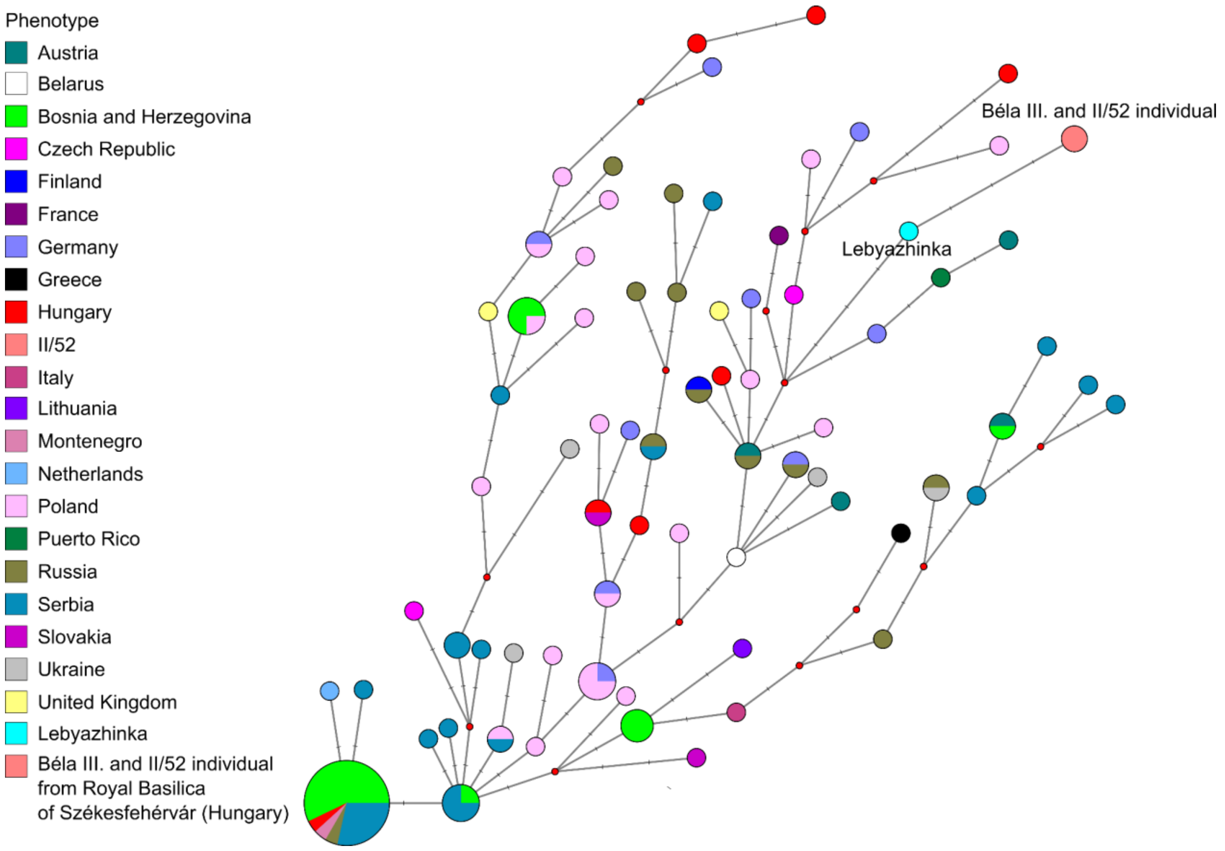


*Figure S42: Median-joining network of R1a-Z282>Z280>L1280 Y-chromosomal subhaplogroup. The network was made using 15 STR loci.*

##### 2.2. Phylogenetic analysis of mitochondrial haplogroups

The median-joining (MJ) network (Fig. S43) based on the mitogenome sequences informs us about the diverse connections of the Hungarians in the CB and the studied VUR groups, which largely clusters as branches (haplogroups) of the mitochondrial phylogenetic tree (www.phylotree.org). In addition, the analysis also shows close intra-site maternal relationships.

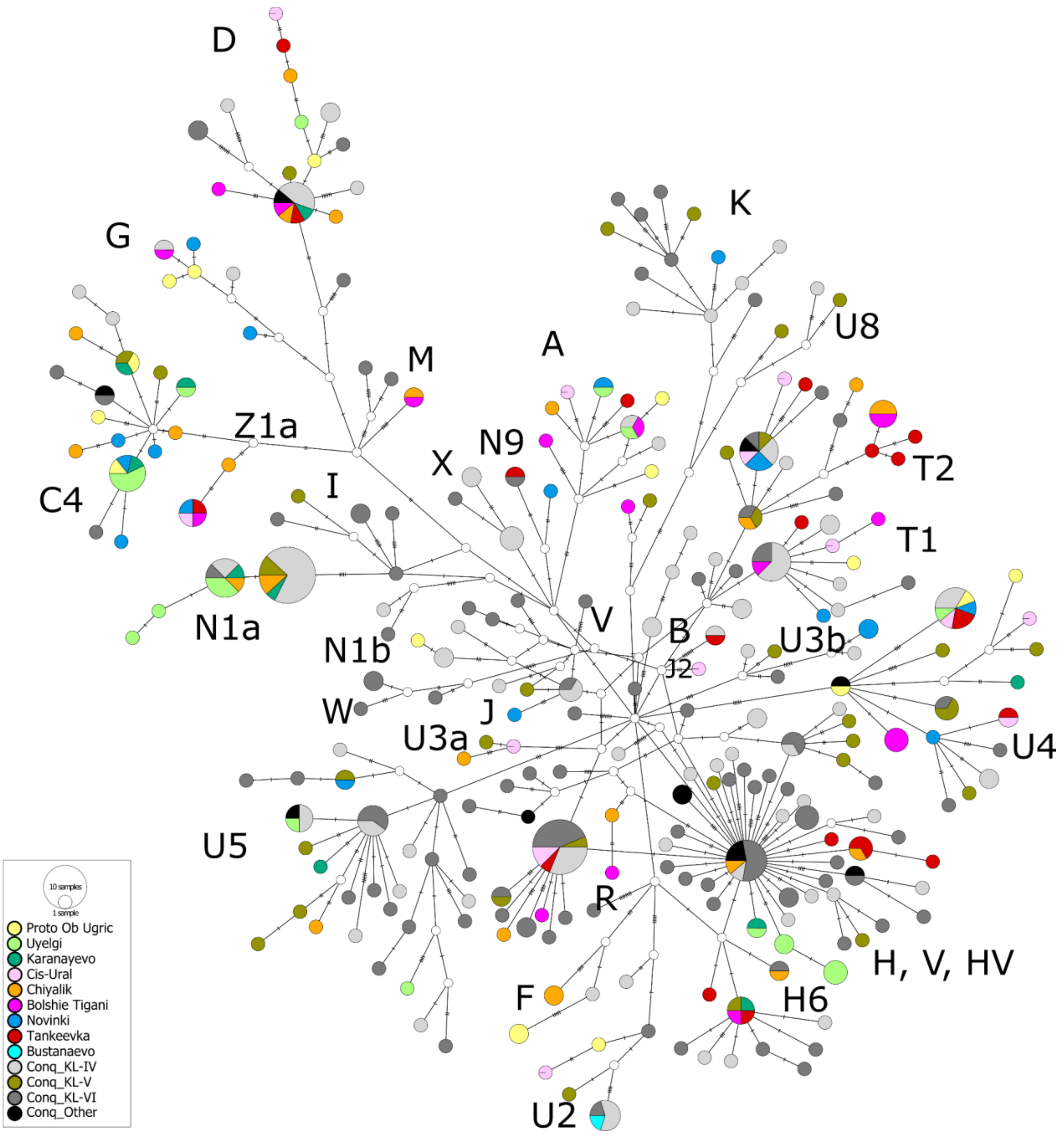


*Figure S43: Median-joining network based on the maternal lines of the studied groups. This analysis shows a diverse system of relationships between our groups and the Conquest period Hungarians. Members of the studied groups are indicated by different colours, major mitochondrial haplogroups are labelled as colored clouds.*

For analysing the maternal relationships, we made neighbour-joining (NJ) phylogenetic trees from those haplogroups (HGs) –detected in the newly analysed 112 samples– that appeared in at least one further site/group of sites associated with the Hungarians (including the Uyelgi, Cis-Ural (CSÁKY ET AL. 2020), and KL-IV-VI groups (NEPARÁCZKI ET AL. 2017, 2018, CSÁKY ET AL. 2020, MAÁR ET AL. 2021)). We highlight below the details of those phylogenetic trees that show the relationships within and between sites/groups. For complete phylogenetic trees, see Extended Data Figure 4. In each figure, the representatives of the groups associated with the Hungarians are highlighted in red. The samples currently examined in this study are marked in red and bold letters.

The haplotype analyses show a close intra-site maternal relationship reflected by the identical mitogenomic sequences at Bolshie Tigani (T2d1b1), Gulyukovo (T2d1b1), Brusyany (U3, T2b24), Ivanov Mis (F1a1c) and Novo Hozyatovo (F1b1e). These people were probably maternally closely related to each other.

The inter-site relationships are more interesting in our investigations because they can connect the studied groups from different regions (Western-Siberia, Volga-Ural region, Carpathian Basin). We detected not only close connections but identical mitogenomes between the studied sites/groups (Fig. S45).

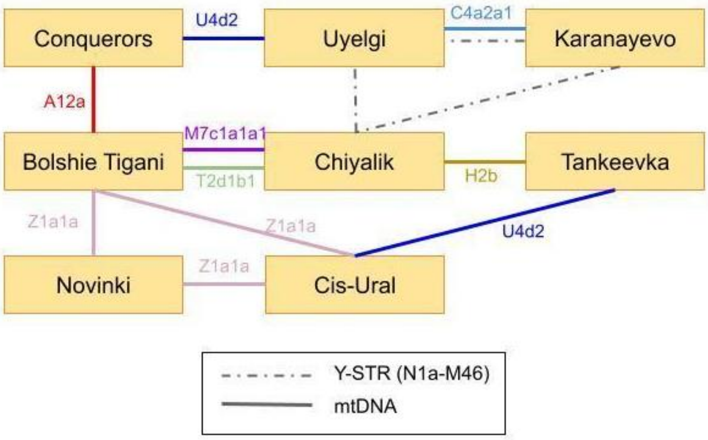


*Figure S44: Identical phylogenetic lineages (haplotypes) between the studied sites/groups. The conquerors group contains in this case the KL-IV and KL-VI groups. Investigated groups from the Volga-Ural region do not show identical maternal connections with representatives of the KL-V group. The proto-Ob-Ugric group does not show such a relationship with the other groups at all.*

##### 2.2.1. Haplogroup A

Seven of the studied individuals are of haplogroup A, belonging to 5 different subhaplogroups:

- A+152+16362: Tankeevka, Shilovka (Novinki group) – this subgroup was detected also in individuals from Uyelgi and Sukhoy Log (Cis-Ural group) (Figure S45B)
- A+152+16362+200: Novo Hozyatovo (Chiyalik group) (Fig. S45A)
- A10: Bolshie Tigani
- A12a: Bolshie Tigani, Vikulovo (proto-Ob-Ugric group) – this subgroup is also present in the Uyelgi site and the KL-IV (Figure S45A)
- A8a1: Ust-Tara (proto-Ob-Ugric group)

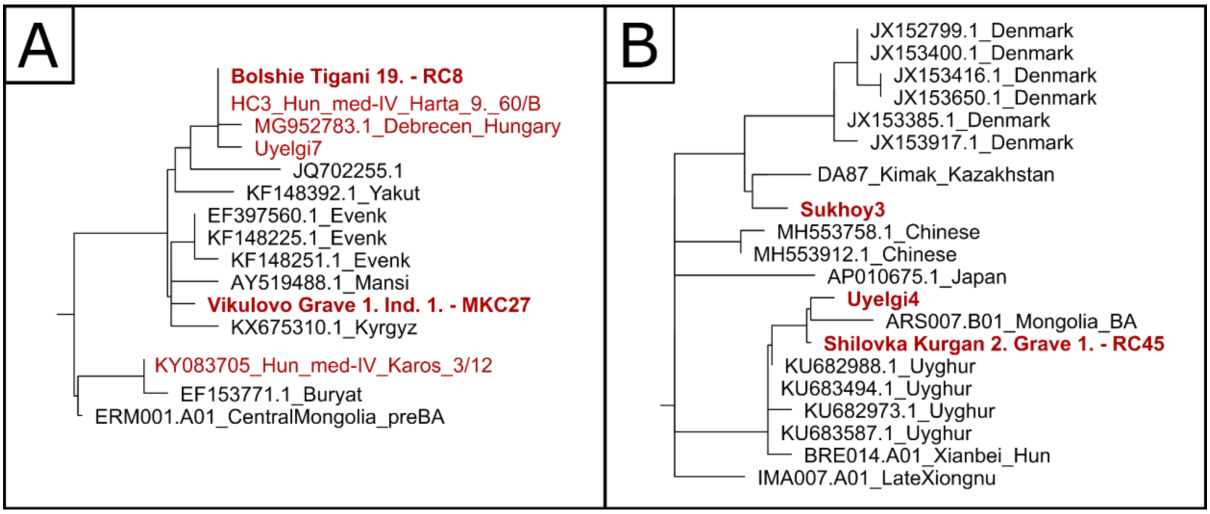


*Figure S45: A: Partial neighbour-joining phylogenetic tree of A12a mitochondrial haplogroup: On this tree, a sample from Bolshie Tigani shares a branch with a Hungarian conqueror, a modern-day Hungarian and a sample from Uyelgi. The conqueror (KL-IV) and a sample from Bolshie Tigani have identical mitochondrial DNA sequences. In the Bolshie Tigani grave 19, a partial horse skeleton and Uralic ceramics accompanied the deceased, together with other archaeological findings with connections to the Hungarians in the Carpathian Basin, thus burial habits, archaeological findings and the biological results support one another. This HG is an example of the maternal lineages of the conquerors coming from the eastern side of the Urals (e.g., Uyelgi), connecting the Volga-Ural region (Bolshie Tigani) with the conquerors of the Carpathian Basin (KL-IV).*

*B: Part of the NJ phylogenetic tree of the mtDNA subhaplogroup A+152+16362: Based on this tree, we detect a connection between samples from the Uyelgi and Novinki group (Shilovka site), which is traced back to the eastern steppe environment.*

##### 2.2.2. Haplogroup C4

We have nine samples and eight subgroups in this HG:

- C4+152: Ivanov-Mis (proto-Ob-Ugric group)
- C4a1a+195: Shilovka (Novinki group)
- C4a1a3: Ust-Tara (proto-Ob-Ugric group)
- C4a1a6: Karanayevo, Novinki (Novinki group) – this subgroup is also detected in the Uyelgi site in several cases ((Figure S47B)
- C4a2a1: Karanayevo – this subgroup is also detected in Uyelgi site and conqueror groups (KL-IX, KL-VI) (Figure S47A)
- C4b: Gulyukovo (Chiyalik group), Karanayevo – this group has also been described in the case of the conquerors (two KL-IV, KL-V) (Figure S46)
- C4b1: Barsov Gorodok (proto-Ob-Ugric group)

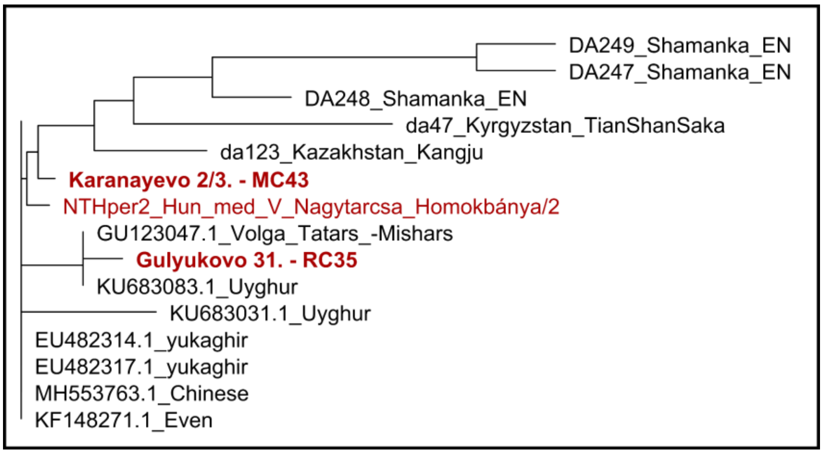


*Figure S46: Part of the NJ phylogenetic tree of C4b mitochondrial HG is signalling a connection between Karanayevo and KL-V conqueror group of an overall Central and East Asia-wide distributed HG.*

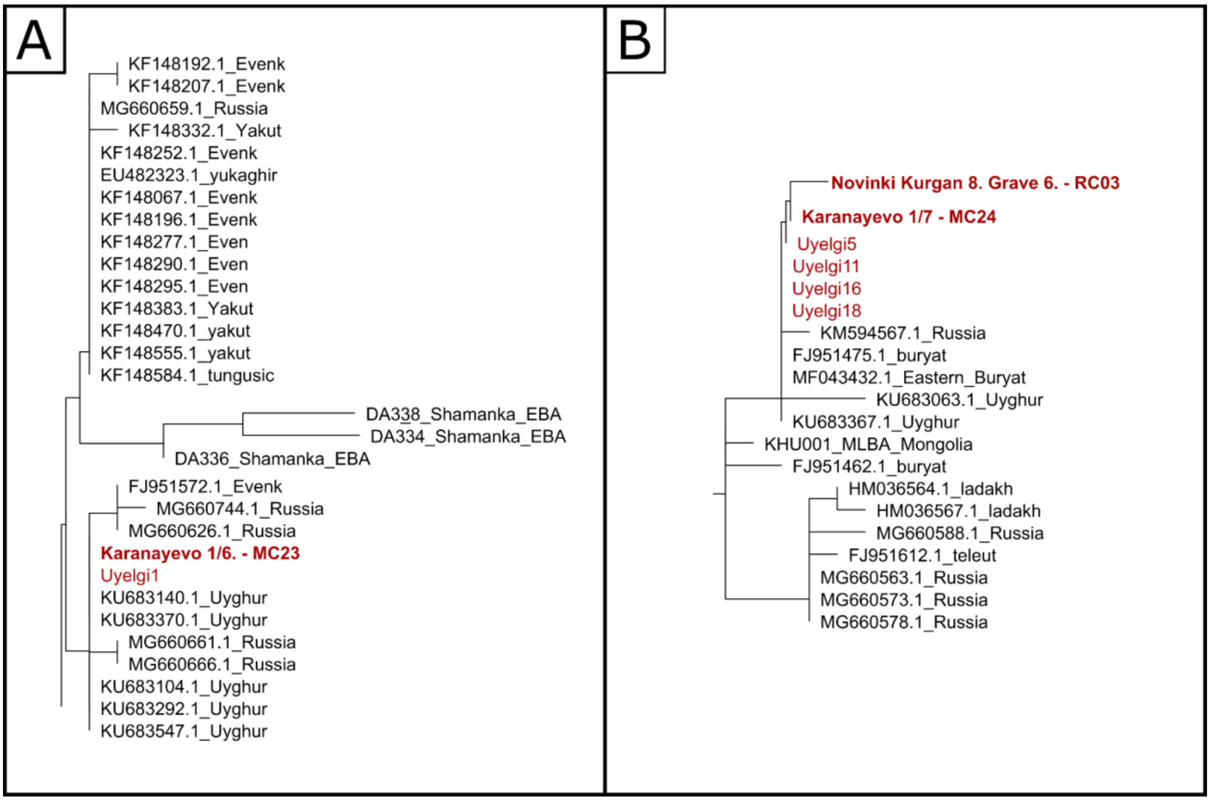


*Figure S47: A: Excerpt from the phylogenetic tree of the C4a2a1 subHG: Samples from Uyelgi and Karanayevo are located on a "steppe branch". They have identical mtDNA sequences, indicating a close maternal connection.*

*B: Part of NJ phylogenetic tree of C4a1a mitochondrial HG. The samples marked in red show close maternal relationships and extensive steppe connections.*

##### 2.2.3. Haplogroup C5

Four samples and three subHG belong to HG C5.

- C5+16093: Malaya Ryazan (Chiyalik group)
- C5a1: Novinki (Novinki group), Gornovo (Chiyalik group) (Figure S48)
- C5c: Gulyukovo (Chiyalik group)

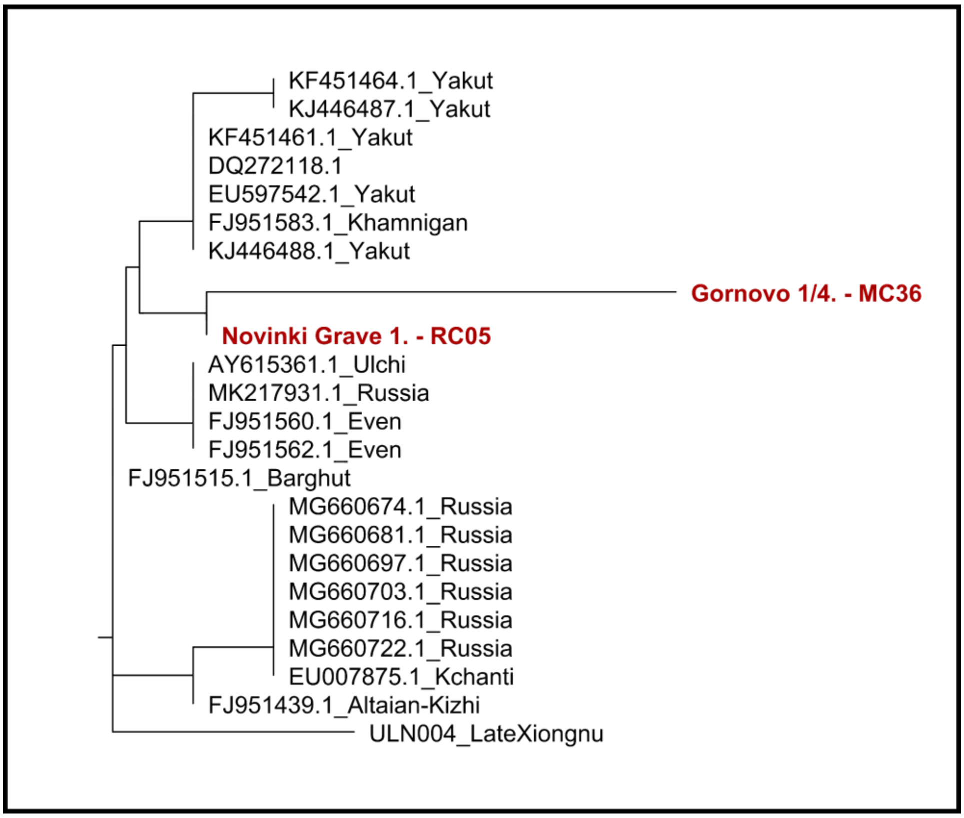


*Figure S48: Partial NJ phylogenetic tree of C5a1 mitochondrial haplogroup shows the connection between the Novinki and Gornovo samples (Novinki and Chiyalik groups) within a Siberian environment*

##### 2.2.4. Haplogroup D4

Nine of our samples belong to six subhaplogroups of this HG:

- D4c2b: Gulyukovo (Chiyalik group) – this subHG points to the Russian Far East
- D4e4: Bolshie Tigani, Karanayevo, Tankeevka, Gulyukovo (Chiyalik group) – this subgroup appears also in conquerors (Figure S49A)
- D4g1b: Bolshie Tigani – this is a branch that contains almost exclusively Chinese and Japanese samples
- D4j2: Tankeevka – this subHG is detected in the Brody site too (Cis-Ural group) (Figure S49B)
- D4j2a: Novo Hozyatovo (Chiyalik group) (Figure S49B)
- D4j4: Panovo (proto-Ob-Ugric group) (Figure S49B)

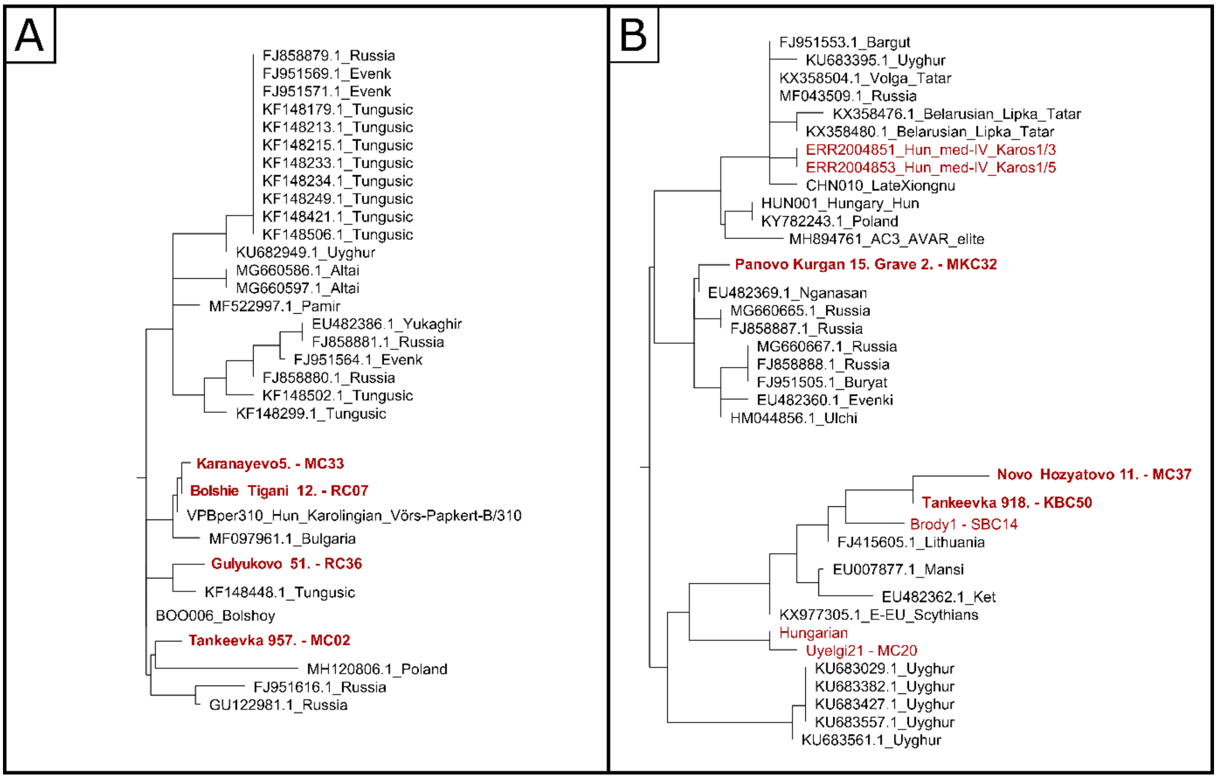


*Figure S49.: A: Partial D4e4 NJ phylogenetic tree. The relationship between the Karanayevo and Bolshie Tigani samples is clear and surprising as well, with a Carolingian period sample from Western Hungary, suggesting a pre-conquest connection between these samples or previously undetected European appearance of this HG.*

*B: Partial NJ phylogenetic tree of D4j mitochondrial subHG. Samples from the Tankeevka, Novo Hozyatovo (Chiyalik group), and Brody (Cis-Ural group) sites are located in a Central-Asian environment on the tree, which are also connected with Uyelgi and a modern-day Hungarian sample. A sample from Panovo (proto-Ob-Ugric group) is on another sub-branch within a Siberian environment.*

##### 2.2.5. Haplogroup G

Five of our samples belong to HG G and these samples fall into 4 subhaplogroups.

- G2a1: Bolshie Tigani, Barsov Gorodok (proto-Ob-Ugric group)
- G2a2a: Brusyany (Novinki group)
- G2a3: Panovo (proto-Ob-Ugric group)
- G3a3: Brusyany (Novinki group)

The studied samples cannot be related to each other based on phylogenetic trees; however, the sample from Brusyany with HG G3a3 is related to a modern-day Hungarian in a steppe environment.

None of these samples are closely related to each other; however, a sample from Brusyany is related to a modern-day Hungarian within a steppe environment.

This HG is represented by G2a1, G2a1d2, and G2a2 subgroups in the conqueror group KL-IV.

##### 2.2.6. Haplogroup H1b

This haplogroup appears at the Bolshie Tigani site and the Cis-Ural group (Sukhoy Log, two H1b2). No relevant relationship was detected between these samples, nor with the conqueror groups. King Béla III. belongs to this haplogroup (NAGY ET AL. 2020, WANG ET AL. 2021). He shows no connection with the Volga-Ural samples, although he connects phylogenetically with an individual from KL-VI (Extended Data Figure 4).

##### 2.2.7. Haplogroup H2

Seven of the studied individuals belong to haplogroup H2 and are in four different subhaplogroups:

- H2a2a: Karanayevo
- H2a2a: Tankeevka
- H2b: three samples from Tankeevka and one from Gulyukovo (Chiyalik group) (Figure S50)

Other subhaplogroups of H2 (H2a1, H2a1c, H2a1n) have been described in the KL-IV.

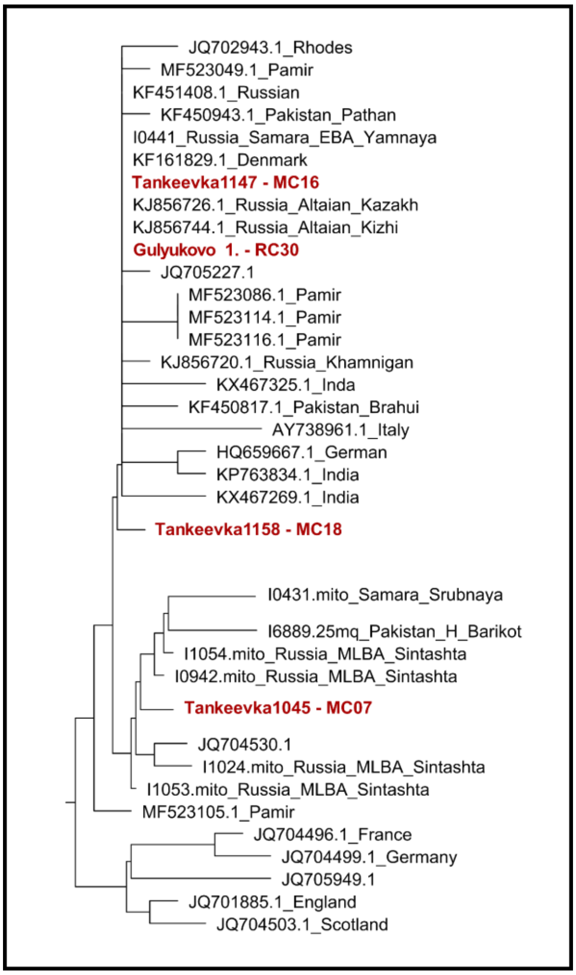


*Figure S50.: Based on the NJ phylogenetic tree of H2b mitochondrial HG, the samples from the Tankeevka and Gulyukovo (Chiyalik group) site show a relationship that can be traced back to the steppe/Central Asia.*

##### 2.2.8. Haplogroup H6a1

We detected two different subhaplogroups of H6a1:

- H6a1a: Bolshie Tigani, Tankeevka
- H6a1b: Karanayevo

Even though the H6a1 haplogroup appears in multiple cases in all conqueror groups (KL-IV-VI), our examined samples show a close relation neither with each other nor with the conquerors.

##### 2.2.9. Haplogroup H13

Two studied samples have mitochondrial HG H13a1d. One from Bolshie Tigani and another from the Gornovo site (Chiyalik group) (Figure S51).

H13 haplogroup is also present in the conquerors (KL-I and KL-VI), but these individuals are not related to our examined samples.

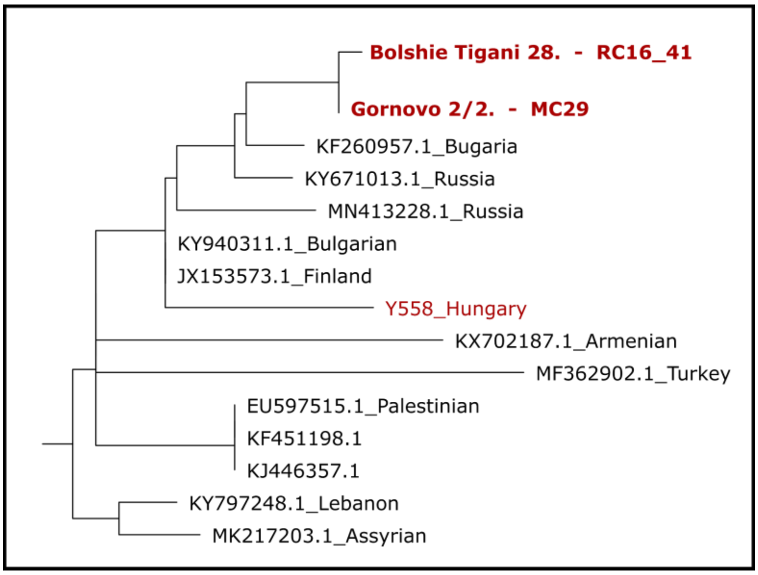


*Figure S51.: Part of H13a1 mitochondrial haplogroup's NJ phylogenetic tree. Two samples from the Bolshie Tigani and Gornovo (Chiyalik group) sites show a connection with each other and with a modern-day Hungarian (Y558_Hungary) as well.*

##### 2.2.10. Haplogroup M7c

We detected an M7c1a1a1 subhaplogroup in the Bolshie Tigani and Gulyukovo (Chiyalik group) sites (Figure S52).

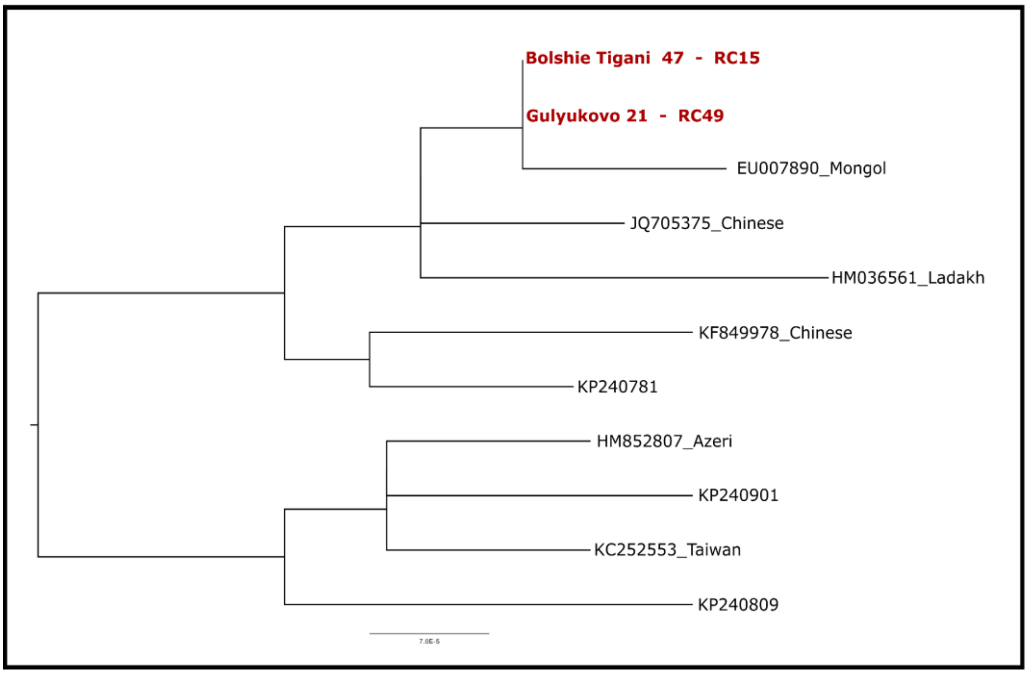


*Figure S52: Neighbour-joining phylogenetic tree of M7c mtDNA HG. The two studied samples have identical positions on this tree, which indicates a clear and close relationship. This maternal line may have entered the study area from the eastern part of the steppe.*

##### 2.2.11. Haplogroup N1a1a1a1

We have five samples in this HG:

- N1a1a1a1a: two samples from Karanayevo, two from Gulyukovo and one from Novo Hozyatovo (Chiyalik group) - this subgroup appears also in conquerors (KL-IV and KL-V) and Uyelgi site (Fig. S53)

Some conquerors belong to the subhaplogroup N1a1a1a1 as well.

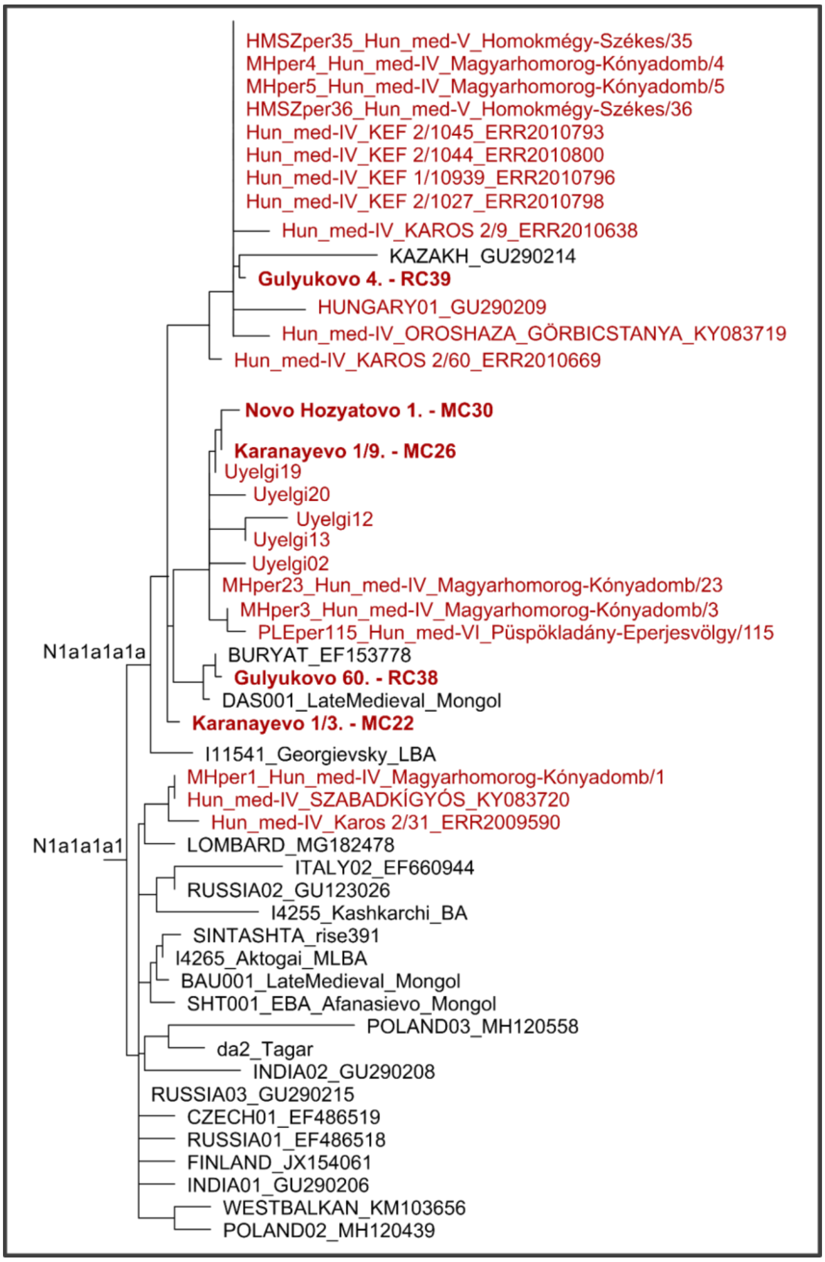


*Figure S53: Partial NJ phylogenetic tree of subhaplogroup N1a1a1a1. Some individuals from KL-IV group belong to N1a1a1a1 subHG. The N1a1a1a1a is splitting into two branches that may have evolved in the Southern Urals. It almost exclusively comprises samples discovered in sites associated with the early Hungarians. Representatives of the KL-IV and KL-VI appear on both N1a1a1a1a branches of the tree, connecting them to the Kushnarenkovo sites in Cis-Urals and Trans-Urals (Uyelgi, Karanayevo) and the Chiyalik culture (Novo Hozyatovo, Gulyukovo) as well.*

##### 2.2.12. Haplogroup T1a1

Our five samples represent the T1a1 haplogroup.

- T1a1: Bolshie Tigani, Tankeevka, Malaya Ryazan (Novinki group), Panovo (proto-Ob-Ugric group) – this subgroup is also present in the Bayanovo site (Cis-Ural group) and the conqueror groups (KL-IV, KL-VI) (Figure S54, Figure S55)
- T1a1d: Bolshie Tigani

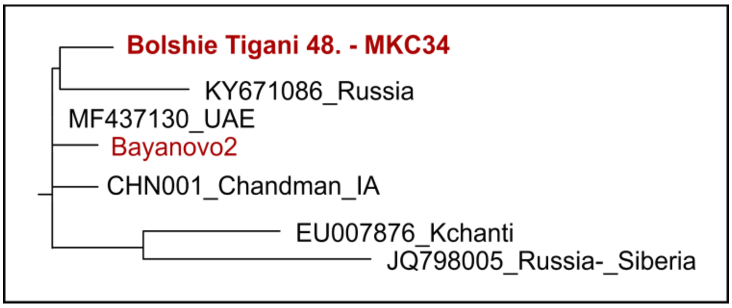


*Figure S54: One part of T1a1 HG's NJ phylogenetic tree: samples from Bolshie Tigani and Bayanovo show a connection.*

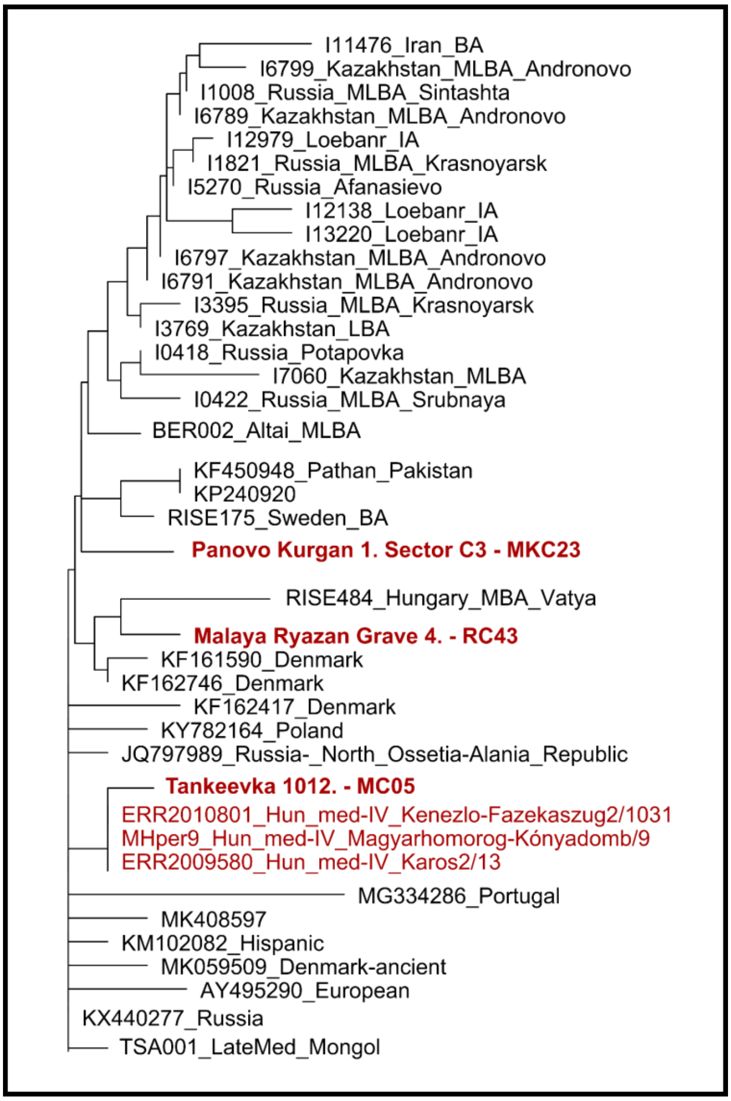


*Figure S55: Second part of T1a1 HG's NJ phylogenetic tree: Tankeevka and three conquerors in the Carpathian Basin (KL-IV) form a sub-branch.*

##### 2.2.13. Haplogroup T2d

Eight of our examined samples belong to the rather uncommon HG T2d (Figure S56).

- T2d1b1: two samples from Bolshie Tigani, three from Tankeevka and two samples from Gulyukovo (Chiyalik group)
- T2d2: Gornovo (Chiyalik group) – the subgroup has also been described in the conquerors (KL-VI)

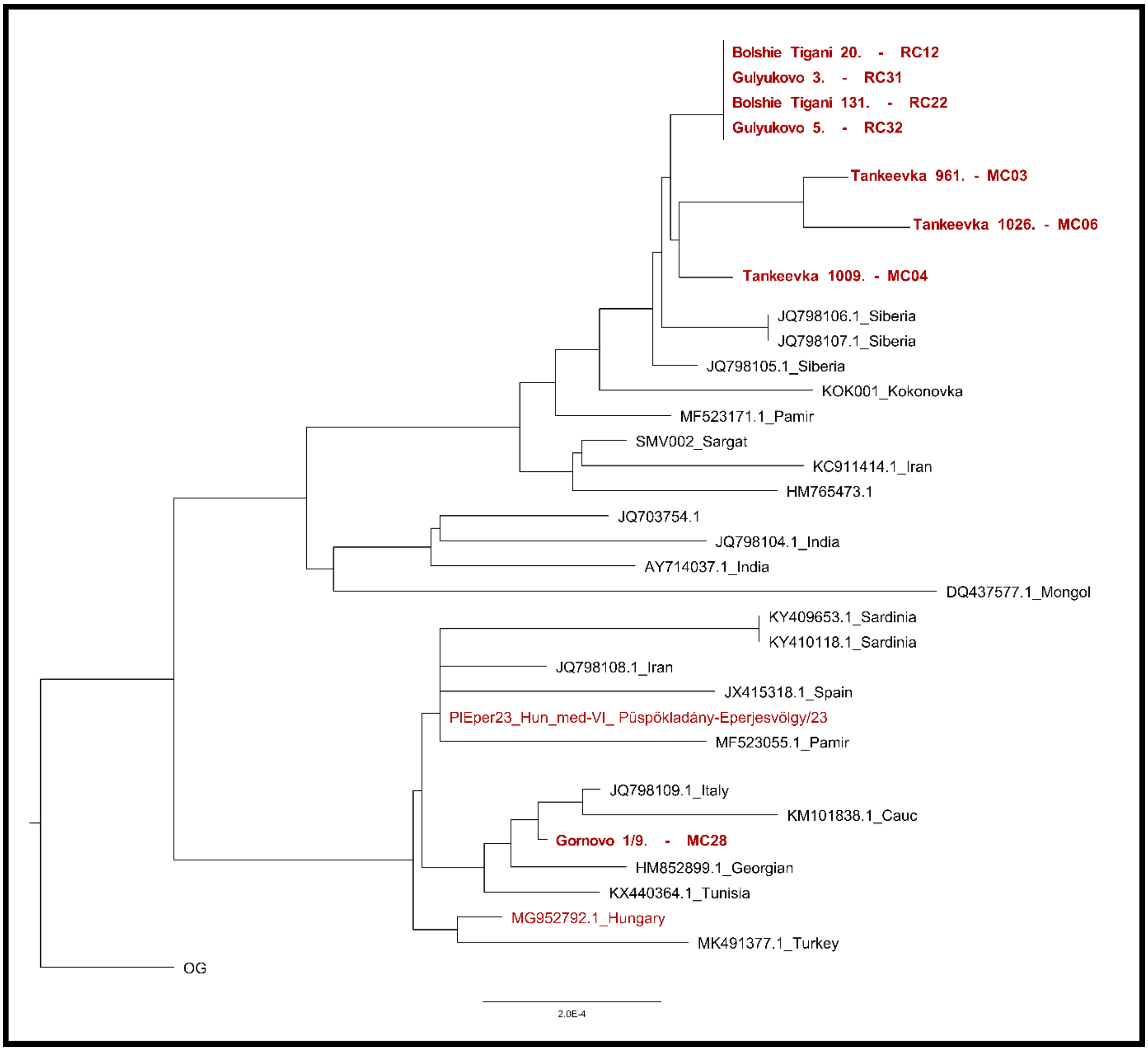


*Figure S56: NJ phylogenetic tree of T2d mitochondrial haplogroup. The samples from the Bolshie Tigani and Gulyukovo (Chiyalik group) sites have identical mtDNA sequences and formed a sub-branch with samples from Tankeevka. These samples form a Siberian sub-branch, while the sample from Gornovo (Chiyalik group) along with a modern-day and conqueror Hungarian branch, together with a phylogeographically undefined sub-branch, is possibly connected to the steppe. In the vicinity of this individual from Gornovo we also find modern-day and conquering Hungarian (KL-VI) individuals.*

##### 2.2.14. Haplogroup U2e1

Two examined samples belong to U2e1 mitochondrial HG:

- U2e1: Panovo (proto-Ob-Ugric group) – this subhaplogroup is also present in the Bartym site (Cis-Ural group) and in KL-VI conqueror group (Figure S57A)
- U2e1b: Bustanaevo – this haplogroup has also been described in conquerors (KL-IV, KL-VI). (Figure S57B)

Other subgroups of this HG (U2e1a1, U2e1b1) were detected in conquerors.

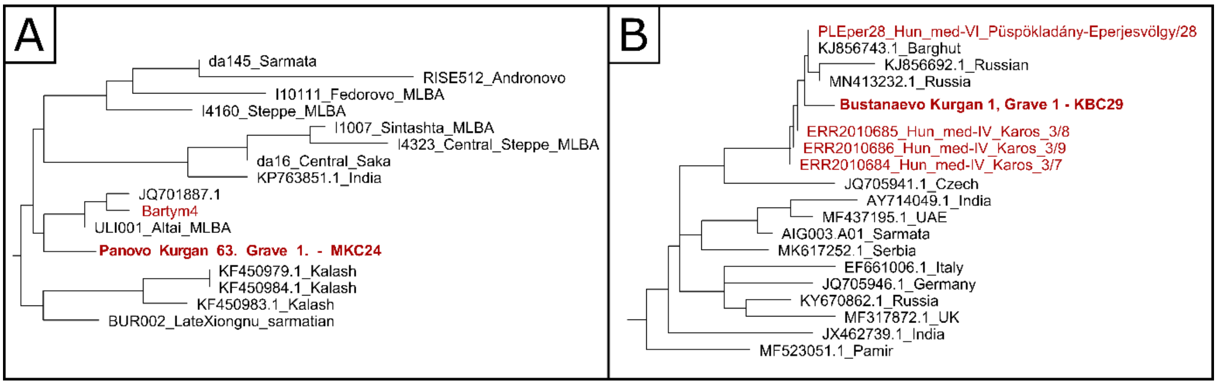


*Figure S57: A: Based on the partial NJ phylogenetic tree of mtDNA HG U2e1 we can see a relatively close relationship between the samples from Panovo (proto-Ob-Ugric) and Bartym (Cis-Ural group) sites within a Central-Asian branch.*

*B: The sample from the Bustanaevo site belongs to the U2e1b mitochondrial HG. Based on the NJ phylogenetic tree of this maternal line the sample from Bustanaevo shows a connection with several conquerors (KL-IV and KL-VI).*

##### 2.2.15. Haplogroup U3

The HG U3 was identified in three tested samples.

- U3a: Gornovo (Chiyalik group) – this sample shows a connection with Near-Eastern samples
- U3b: two samples from Brusyany (Novinki group) – these mitogenomes are identical and are connected with Near-Eastern individuals from around the Caspian Sea.

Several subgroups of the U3 are present in Sukhoy Log (Cis-Ural group) (U3a1) and conquerors (KL-IV-VI) (U3a1b, U3b1b, U3b2, U3b2a, U3b3), but show no closer relationship with the samples we examined.

##### 2.2.16. Haplogroup U4

The subhaplogroups of the U4 mitochondrial HG include 12 samples examined by us.

- U4: Bolshie Tigani
- U4a1d: Tankeevka, Lebyazhinka (Novinki group) – this subHG is also present in Bartym site (Cis-Ural group) (Figure S58)
- U4a2: Bolshie Tigani 2x, Barsov Gorodok (proto-Ob-Ugric group) - the haplogroup has also been described in the conquerors (KL-I, KL-V, KL-VI)
- U4b1a1a: Ivanov Mis (proto-Ob-Ugric group)
- U4b1b1+16311: Karanayevo
- U4d2: Tankeevka 2x, Brusyany (Novinki group), Barsov Gorodok (proto-Ob-Ugric group) - this subgroup also appears at the Uyelgi site, Bartym site (Cis-Ural group), and in the KL-IV conqueror group (Figure S59)

Three samples from the Bolshie Tigani site show a close relationship with each other, and two of them have identical mtDNA sequences.

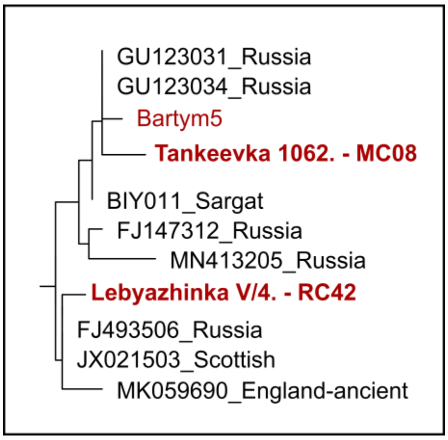


*Figure S58: One part of U4a1d mitochondrial HG's NJ phylogenetic tree: Samples from the Bartym site (Cis-Ural group) and Tankeevka site are grouped together on this tree. The proximity of a Sargat individual may suggest succession between these groups and proto-Hungarians. Sample from the Lebyazhinka site (Novinki group, although connected to the Ugric people, see in Supplementary Material, Chapter A) is on a different branch of the tree.*

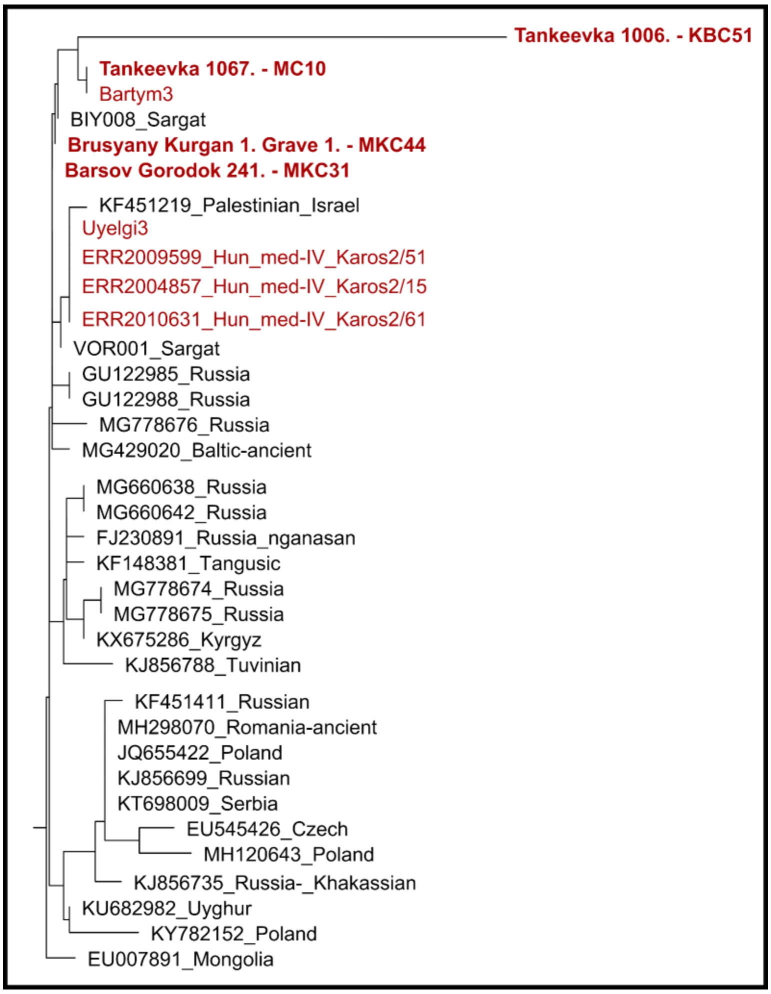


*Figure S59: Partial U4d2 mtDNA HG's NJ phylogenetic tree. The studied samples (Tankeevka, Brusyany (Novinki group), Barsov Gorodok (proto-Ob-Ugric group)) are near individuals from the steppe and Eastern Europe. These samples show a slightly distant relationship between individuals from the Uyelgi site and the conqueror group (KL-IV). There is a sample from Sargat-culture on both branches, which is another argument that some lines have existed in the Volga-Ural region since the Iron Age.*

##### 2.2.17. Haplogroup U5a

Three of our samples belong to HG U5a, but they fall into three different subgroups:

- U5a1d2a1: Novo Hozyatovo (Chiyalik group)
- U5a1g1: Karanayevo
- U5a2a1+152: Brusyany (Novinki group) (Figure S60)

Several subgroups of this HG were also described in all three conqueror groups.

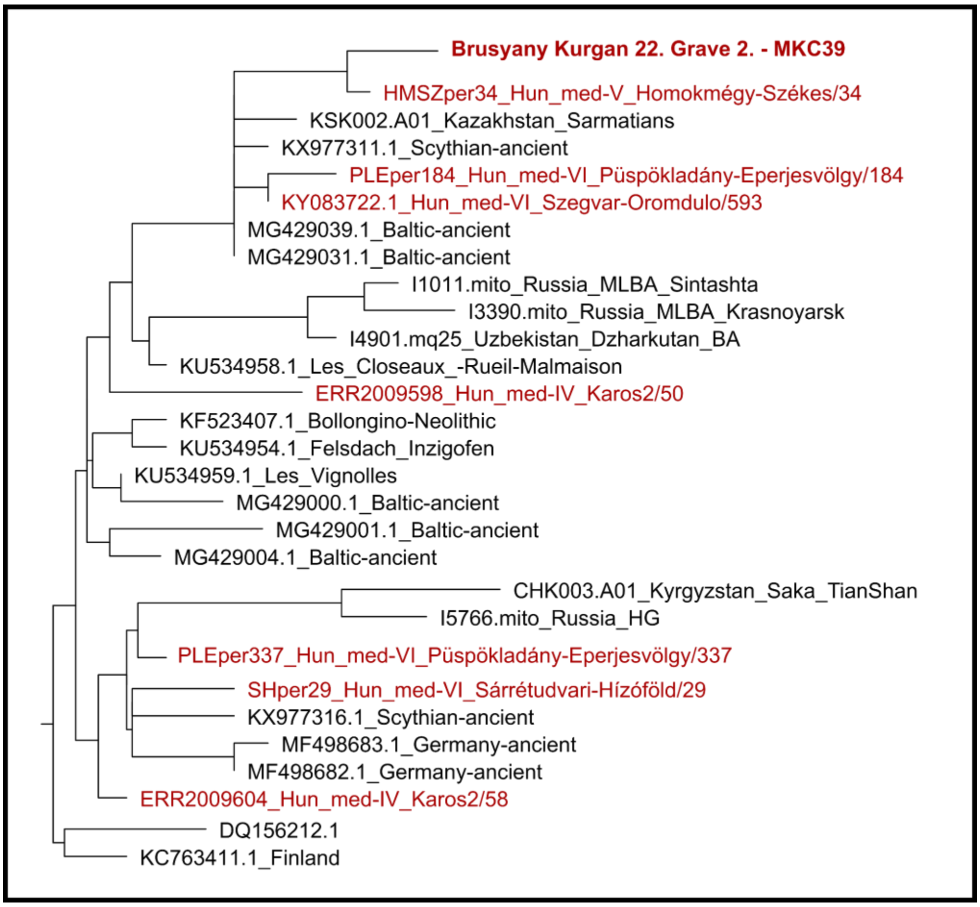


*Figure S60: Neighbour-joining phylogenetic tree of U5a2a1+152 mtDNA haplogroup: Several conquerors appear on this tree next to the Brusyany samples. This maternal line may be a link between individuals from the two different regions regarding HMSZ/34, but other Hungarian cases show rather indirect connections.*

##### 2.2.18. Haplogroup Z1a

Four of our samples belong to two subhaplogroups of this HG:

- Z1a: Gulyukovo (Chiyalik group)
- Z1a1a: Bolshie Tigani, Tankeevka, Mullovka (Novinki group) – this subgroup appears also in Bayanovo (Cis-Ural group) (Figure S61)

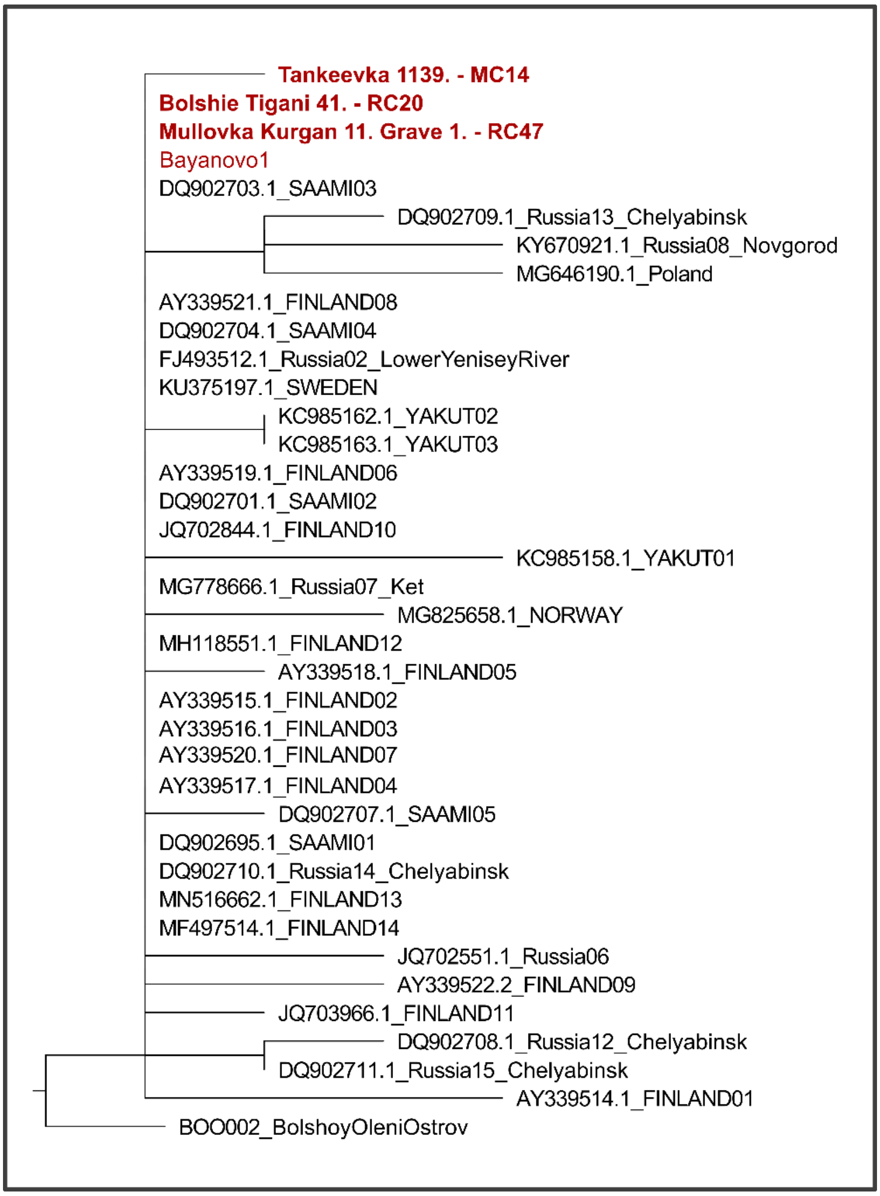


*Figure S61: NJ phylogenetic tree of HG Z1a1a mitochondrial haplogroups: Z1a1a samples from Bolshie Tigani, Mullovka, and Bayanovo have identical mitochondrial sequences. Z1a spread from Siberia through the Volga-Ural region to northern Europe, where it was already present 3500 years ago (LAMNIDIS ET AL. 2018). This tree demonstrates the connection between the Volga-Ural region and northern Europe (LAMNIDIS ET AL. 2018, DER SARKISSIAN ET AL. 2013, MALYARCHUK ET AL. 2010, INGMAN-GYLLENSTEN 2007a), which may also be the result of an interaction that took place later than the previously assumed gene flows (5th-4th millennia BCE and 1st millennium BCE (INGMAN-GYLLENSTEN 2007b).*

#### 3. References to chapter B


CSÁKY ET AL. 2020: Csáky, V., Gerber, D., Szeifert, B., Egyed, B., Stégmár, B., Botalov, S.G., Grudochko, I.V., Matveeva, N.P., Zelenkov, A.S., Sleptsova, A.V., *et al.* (2020) Early medieval genetic data from Ural region evaluated in the light of archaeological evidence of ancient Hungarians. *Sci. Rep.*, **10**.

DER SARKISSIAN ET AL. 2013: Der Sarkissian, C., Balanovsky, O., Brandt, G., Khartanovich, V., Buzhilova, A., Koshel, S., Zaporozhchenko, V., Gronenborn, D., Moiseyev, V., Kolpakov, E., *et al.* (2013) Ancient DNA Reveals Prehistoric Gene-Flow from Siberia in the Complex Human Population History of North East Europe. *PLoS Genet.*, **9**.

INGMAN-GYLLENSTEN 2007a: Ingman, M. and Gyllensten, U. (2007) A recent genetic link between Sami and the Volga-Ural region of Russia. *Eur. J. Hum. Genet.*, **15**, 115–120.

INGMAN-GYLLENSTEN 2007b: Ingman, M. and Gyllensten, U. (2007) Rate variation between mitochondrial domains and adaptive evolution in humans. *Hum. Mol. Genet.*, **16**, 2281–2287.

LAMNIDIS ET AL. 2018: Lamnidis, T.C., Majander, K., Jeong, C., Salmela, E., Wessman, A., Moiseyev, V., Khartanovich, V., Balanovsky, O., Ongyerth, M., Weihmann, A., *et al.* (2018) Ancient Fennoscandian genomes reveal origin and spread of Siberian ancestry in Europe. *Nat. Commun.*, **9**, 5018.

MAÁR ET AL. 2021: Maár, K., Varga, G.I.B., Kovács, B., Schütz, O., Maróti, Z., Kalmár, T., Nyerki, E., Nagy, I., Latinovics, D., Tihanyi, B., *et al.* (2021) Maternal lineages from 10-11th century commoner cemeteries of the Carpathian Basin. *Genes (Basel).*, **12**.

MALYARCHUK ET AL. 2010: Malyarchuk, B., Derenko, M., Denisova, G. and Kravtsova, O. (2010) Mitogenomic diversity in Tatars from the Volga-Ural region of Russia. *Mol. Biol. Evol.*, **27**, 2220–2226.

MANTEL 1967: Mantel, N. (1967) The detection of disease clustering and a generalized regression approach. *Cancer Res.*, 209–220.

NAGY ET AL. 2020: Nagy, P.L., Olasz, J., Neparáczki, E., Rouse, N., Kapuria, K., Cano, S., Chen, H., Di Cristofaro, J., Runfeldt, G., Ekomasova, N., *et al.* (2020) Determination of the phylogenetic origins of the Árpád Dynasty based on Y chromosome sequencing of Béla the Third. *Eur. J. Hum. Genet.*, 164–172.

NEPARÁCZKI ET AL. 2017: Neparáczki, E., Kocsy, K., Tóth, G.E., Maróti, Z., Kalmár, T., Bihari, P., Nagy, I., Pálfi, G., Molnár, E., Raskó, I., *et al.* (2017) Revising mtDNA haplotypes of the ancient Hungarian conquerors with next generation sequencing. *PLoS One*, **12**.

NEPARÁCZKI ET AL. 2018: Neparáczki, E., Maróti, Z., Kalmár, T., Kocsy, K., Maár, K., Bihari, P., Nagy, I., Fóthi, E., Pap, I., Kustár, Á., *et al.* (2018) Mitogenomic data indicate admixture components of Central-Inner Asian and Srubnaya origin in the conquering Hungarians. *PLoS One*, **13**.

OLASZ ET AL. 2018: Olasz, J., Seidenberg, V., Hummel, S., Szentirmay, Z., Szabados, G., Melegh, B. and Kásler, M. (2018) DNA profiling of Hungarian King Béla III and other skeletal remains originating from the Royal Basilica of Székesfehérvár. *Archaeol. Anthropol. Sci.*, 1345-1357.

SLATKIN 1995: Slatkin, M. (1995) A measure of population subdivision based on microsatellite allele frequencies. *Genetics*, **139**, 457–462.

WANG ET AL. 2021: Wang, C.C., Posth, C., Furtwängler, A., Sümegi, K., Bánfai, Z., Kásler, M., Krause, J. and Melegh, B. (2021) Genome-wide autosomal, mtDNA, and Y chromosome analysis of King Bela III of the Hungarian Arpad dynasty. *Sci. Rep.*, **11**.
